# PDGFRα signaling in cardiac fibroblasts modulates quiescence, metabolism and self-renewal, and promotes anatomical and functional repair

**DOI:** 10.1101/225979

**Authors:** Naisana S. Asli, Munira Xaymardan, Ralph Patrick, Nona Farbehi, James Cornwell, Elvira Forte, Ashley J. Waardenberg, Vaibhao Janbandhu, Scott Kesteven, Vashe Chandrakanthan, Helena Malinowska, Henrik Reinhard, Peter Schofield, Daniel Christ, Ishtiaq Ahmed, Romaric Bouveret, Surabhi Srivastava, Rakesh K. Mishra, Jyotsna Dhawan, James J.H. Chong, Robert Nordon, Peter Macdonald, Robert M. Graham, Michael Feneley, Richard P. Harvey

## Abstract

The interstitial and perivascular spaces of the mammalian heart contain a highly interactive tissue community essential for cardiac homeostasis, repair and regeneration. Mesenchymal cells (fibroblasts) are one of the most abundant cell types, playing key roles as sentinels, tissue architects, paracrine signaling hubs and lineage precursors, and are linked to heart disease through their roles in inflammation and fibrosis. Platelet-derived growth factors (PDGFs) are secreted by several cell types involved in cardiac injury and repair, and are recognized mitogens for cardiac fibroblasts and mesenchymal stem cells. However, their roles are complex and investigations of their impact on heart repair have produced contrasting outcomes, leaving therapeutic potential uncertain. Here, we use new approaches and tools, including single cell RNA sequencing, to explore cardiac fibroblast heterogeneity and how PDGF receptor α (PDGFRα) signaling impacts fibroblasts during heart repair. Short-term systemic delivery of PDGF-AB to mice from the time of myocardial infarction (MI) led to enhanced anatomical and functional recovery. Underpinning these benefits was a priming effect, in which PDGF-AB accelerated exit of fibroblasts from quiescence and induced a higher translational biosynthetic capacity in both fibroblasts and macrophages without triggering fibrosis. Our study highlights the significant biosynthetic heterogeneity and plasticity in cardiac fibroblast populations, and suggests a rationale for a novel therapeutic approach to cardiac injury involving controlled stimulation of fibroblast activation.

## INTRODUCTION

In humans, myocardial infarction (MI) is the most common form of acute cardiac injury leading to the death of millions of cardiomyocytes (CM), reduced heart function and high mortality. Therefore, achieving heart regeneration after MI is a major clinical imperative. The hearts of zebrafish and some urodele amphibia show spectacular regeneration through cardiomyocyte (CM) dedifferentiation and proliferation^1^, and a similar capacity exists in neonatal mammalian hearts^2^. However, in the hearts of adult mammals, CM turnover is low^3^ and replacement after MI is thought to be insignificant.

The heart also contains a variety of stromal cell types, including fibroblasts, endocardial, epicardial, coronary and lymphatic vascular cells, nerves, immune cells, and potentially stem and progenitor cells, although a comprehensive understanding of this complex interacting cellular community is lacking. It is intuitive that achieving cardiac regeneration in man will involve preserving and restoring connective tissue and vasculature architecture as well as CMs. Fibroblasts are one of the most abundant cardiac cell types, acting as major sensing and signaling hubs through their progenitor, paracrine, mechanical and electrical functions^4–7^, and are conspicuously linked to heart disease through their roles in inflammation and fibrosis. After MI, stromal fibroblasts proliferate and differentiate into myofibroblasts, which stabilize the infarct through secretion of a collagen-rich matrix^7^. A common feature of regenerating tissues is that timely resolution of inflammation is critical for limiting fibrosis and allowing tissue replacement^8^. However, in the injured heart, inhibition of inflammation can lead to arrest of repair and autoimmunity, while inhibition of fibrosis can promote functional recovery^9^, albeit at the cost of a predisposition to cardiac rupture^10^. Thus, cellular interactions driving inflammation and fibrosis are finely balanced, and a deeper understanding of relevant pathways will be needed to point the way to novel regenerative targets.

We previously defined a population of cardiac fibroblast colony-forming cells (termed colony forming units - fibroblast; cCFU-F) that exhibit stem cell qualities, including long-term self-renewal and multi-lineage differentiation capacity *in vitro*, akin to those of bone marrow (BM) mesenchymal stem cells (MSCs)^11,12^. In the BM, accumulating evidence show that cells enriched for MSC colony-forming units correspond to multipotent osteogenic/chondrogenic/stromal lineage progenitors *in vivo*, and that their descendants support different stages of hemopoiesis through niche functions^13^. A potentially analogous population in skeletal muscle supports satellite stem cell activation, and gives rise to fibro-adipogenic cells in pathology^14^. We hypothesize, therefore, that aspects of fibroblast activation, expansion, differentiation and/or renewal are governed by stem cell principles^11,12,15^

Virtually all cardiac fibroblasts express Platelet-Derived Growth Factor Receptor α (PDGFRα)^16,17^ and PDGF ligands are expressed by multiple cell types involved in the injury response. In mammals, there are 4 PDGF ligands which form 5 dimeric species (AA, AB, BB, CC, DD) that bind with different affinities to homo- and/or heterodimers of the two cognate receptors (αα, αβ, ββ). The roles of PDGF signaling in the heart are complex. Genetic studies show that the PDGF-BB/PDGFRβ signaling axis is involved in both developmental and adult coronary angiogenesis, principally for recruitment of perivascular cells^18^ but also acts in induction of pro-angiogenic signaling from CM under stress conditions^19^. PDGFRα is essential for formation of interstitial fibroblasts from epicardium in development^20^, and has been implicated in proliferation, migration and differentiation of a variety of mesenchymal cell types including MSCs and cardiac fibroblasts^21^. Constitutively active mutants of PDGFRα lead to organ-wide fibrosis and cancer^22^, while PDGFR inhibition in humans can cause lethal cardiomyopathies^23^. In preclinical models, the therapeutic impact of delivering different PDGF ligands before or at the time of MI has been tested based principally on a pro-angiogenic rationale; however, this has yielded diverse outcomes from improved cardiac repair to frank fibrosis and heart failure^24–28^. Thus, how PDGF ligands and receptors function in cardiac homeostasis and injury repair is not well understood, and the therapeutic potential of modulating PDGF/PDGFR signaling remains unclear.

Here, we use new approaches and tools, including scRNAseq, to investigate fibroblast heterogeneity, and the impact of manipulating PDGFRα signaling on cardiac fibroblasts *in vitro* and *in vivo*. Short-term systemic delivery of PDGF-AB to mice from the time of MI led to significantly improved anatomical and functional repair. Enhanced PDGFRα signaling stimulated the rate of cell cycle engagement of fibroblasts exiting quiescence, and induced a higher ribosomal biosynthetic state in both fibroblasts and macrophages *in vivo*, without triggering fibrosis. By contrast, inhibition of PDGFRα signaling in fibroblasts induced quiescence and reduced self-renewal, associated with major changes in the transcriptome and potentially chromatin structure. Our results show a pro-reparative role for enhanced PDGFRα signaling in the heart involving controlled fibroblasts and macrophage activation.

## RESULTS

### Modulation of PDGFR signaling on cell cycle dynamics in vitro

Our previous work has shown that there are two major resting fibroblasts sub-sets which express PDGFRα (a generic fibroblast marker^6,16^) but are distinguishable by high versus low expression of the stem cell marker SCA1^11,12^ (Fig. S1A). The SCA1^+^PDGFRα^+^ (S^+^P^+^) population is enriched for cCFU-F colony-forming units^11,12^, attesting to functional differences between these populations. We recently profiled cardiac fibroblasts using scRNAseq and cell clustering revealed two major fibroblast sub-populations expressing high and low levels of the *Sca1* gene^16^.

We first explored how PDGF signaling influenced the cell cycle of S^+^P^+^ cells grown *in vitro* under MSC conditions. Among PDGF ligands tested (AA, AB, BB), PDGF-AB was the most potent in stimulating colony formation (see below), so we used it in these experiments. Single cell lineage tracking of freshly isolated S^+^P^+^ cells showed that ~60% underwent cell division in generation (gen) 0 and, of these, 60% divided at least once more (gens >0; Fig. 1A). In gen 0, PDGF-AB strongly stimulated the probability of cell cycle entry and the overall proportion of cells dividing (Fig. 1B), and increased EdU incorporation over the first 24 hrs (Fig. 1C). Total cell yield was increased at the end of passage P0 (Fig. 1D), with no change in cell death (Fig. S1B). By contrast, the small molecule PDGFRα/β inhibitor, AG1296, had no significant effect on the rate of cell cycle entry or the proportion of cells dividing in gen 0 (Fig. 1B). In gens >0, PDGF-AB also stimulated the probability of cell division, although to a lesser extent than in gen 0, whereas AG1296 slightly decreased the probability, with accompanying changes in the proportion of cells dividing (Fig. 1B).

**Figure 1:**
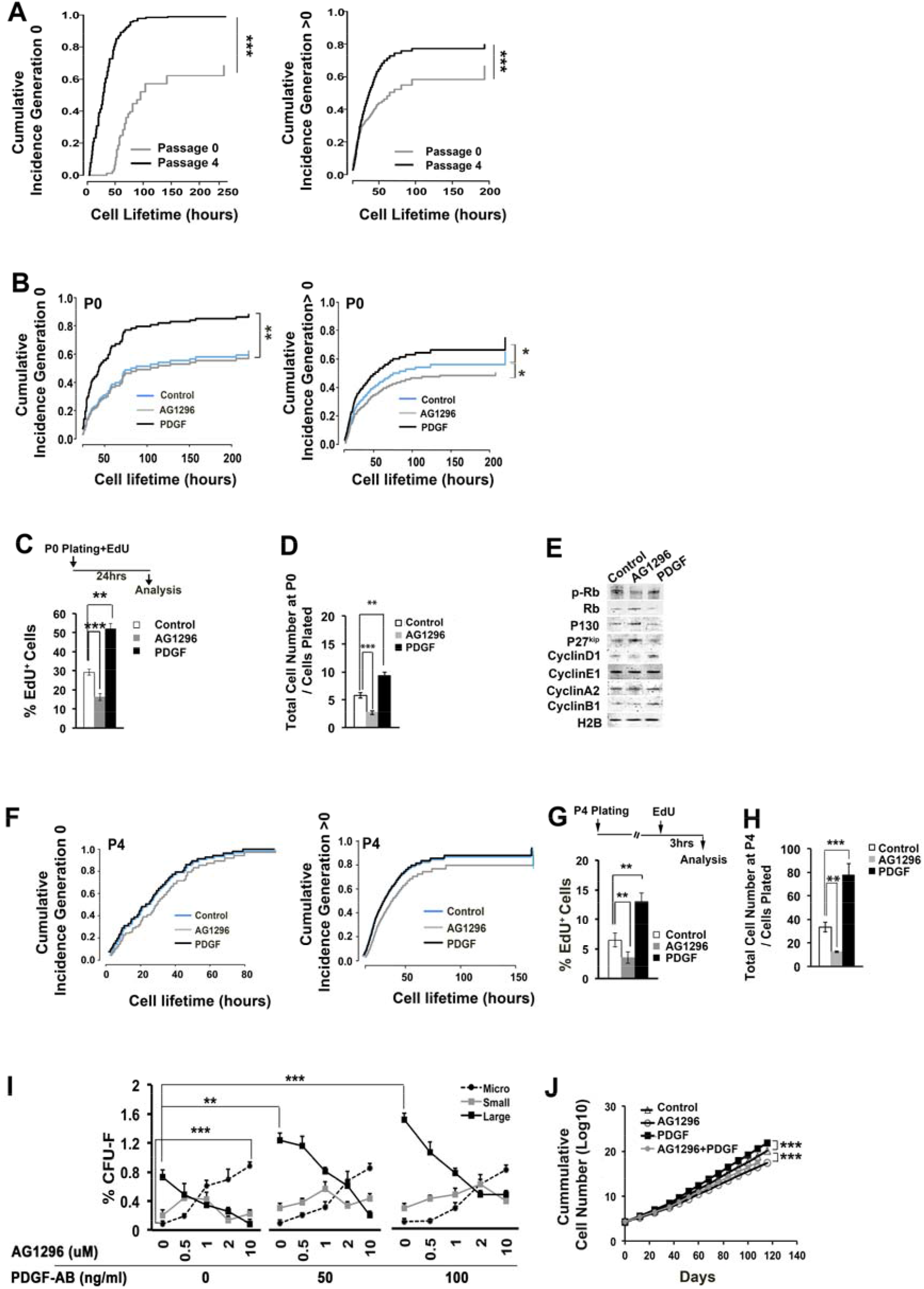
Role of PDGFRα signaling in cCFU-F cultures. (A) Accumulated proportion of cells undergoing division over time in freshly plated S^+^P^+^ cells across P0, and established cCFU-F cultures re-plated at the P4, scored separately in generation 0 (whose birth was not observed) and generations >0. (B) Single cell analysis of cumulative cell division in freshly plated (P0) S^+^P^+^ cells with and without PDGF-AB and AG1296, scored separately in generations 0 and >0. (C) EdU incorporation in S^+^P^+^ cultures with 24hr pulse. (D) Total cell number in cCFU-F cultures treated with vehicle (control), AG1296 (10μM) or PDGF-AB (100ng/ml) scored at the end of P0. (E) Western blot analysis of indicated cell cycle regulators in S^+^P^+^ cultures at P0. (F) Live cell tracking analysis of the accumulated proportion of cells undergoing division over time across P4 cells treated with vehicle, AG1296 or PDGF-AB, scored separately in generation 0 and generations >0. (G,H) EdU incorporation and total cell number, respectively, arising from treatment of cCFU-F cultures across P4 with AG1296 or PDGF-AB. (I) Colony analysis (large, small, micro-colonies) in S^+^P^+^ cells scored at P0 (day 12), with indicated concentrations of PDGF-AB ligand and PDGFR signaling inhibitor AG1296. (J) Long-term cultures of S^+^P^+^ cells with or without PDGF-AB (100ng/ml) and AG1296 (10μM). All comparisons: *p<0.05; ** p<0.01; *** p<0.001.

Western blotting of protein extracts harvested from ligand and inhibitor treated cultures at the end of P0 showed reciprocal changes in both positive and negative cell cycle regulators, including Rb, p130, cyclins G1 and D1, and the cyclin-dependent kinase inhibitor p27^kip1^ (Fig. 1E).

We repeated these experiments on cCFU-F cultures at P4, which are fully adapted to the *in vitro* culture. After re-plating, 100% of cells divided in gen 0 and ~80% divided again (gens >0; Fig. 1F). However, neither PDGF-AB nor inhibitor had any impact on the probability of cell cycle entry and proportion of cells dividing (Fig. 1F), or cell death (Fig. S1B), even though there were observable changes in cell cycle regulators (Fig. S1C).

These data reveal that the dominant effect of increased PDGF signaling in S^+^P^+^ cell cultures was to increase the probability of cell cycle entry and the proportion of cells initially dividing. However, PDGF-AB had only a minor effect on the subsequent rate of cell cycle progression, and no effect once cultures were fully established. Paradoxically, ligand and inhibitor treatment of P4 cells still showed a pronounced increase and decrease, respectively, in EdU incorporation and total cell number at the end of P4 (Fig. 1G,H). We hypothesized, therefore, that PDGFR signaling was impacting the dynamics of self-renewal.

### Impact of modulating PDGF signaling on cCFU-F self-renewal

In our colony assay, around 1 in 120 S^+^P^+^ cells form large colonies^11^. We have also previously shown using single cell tracking of cCFU-F cultures derived from *Pdgfa^GFP/+^* mice (expressing a nuclear-localized fusion between histone H2B and eGFP under control of the *Pdgfra* locus^11^), that GFP^+^ cells are capable of expansive self-renewal (probability of division generating two GFP^+^ daughters >0.5)^29^. Furthermore, the size of cCFU-F colonies is a proxy for their clonal growth, with only large colony cells capable of undergoing self-renewal over many passages (>200 days)^11^. During culture there is a degree of spontaneous differentiation of cells into large flat myofibroblast-like cells, which rarely divide (Fig. S1D) - this occurs infrequently in large colonies, whereas in micro-colonies, which do not re-plate, virtually all cells are differentiated^11^. These experiments show that cCFU-F exist in a spectrum of cell states correlating with their self-renewal ability and propensity for spontaneous differentiation.

Using the colony readout, we explored the impact of altering PDGF signaling. S^+^P^+^ cells were isolated from healthy adult hearts and cultured in a cCFU-F assay with and without increasing doses of PDGF-AB ligand and/or AG1296 inhibitor until the end of P0 (day 12). PDGF-AB increased the number of large colonies ~2-fold in a dose-dependent manner (Fig. 1I), as well as cell yield, as noted above (Fig. 1D). AG1296 decreased the number of large colonies ~3-8-fold and cell yield. The addition of increasing amounts of a specific ligand-blocking PDGFRα monoclonal antibody (mAb) produced the same colony profile as increasing doses of AG1296 (Fig. S1E), confirming the role of PDGFRα in these effects.

We asked whether inhibition of PDGFRα signaling was accompanied by a permanent change in cell state, e.g. whether they became committed to differentiate. We cloned large, small and micro colonies from inhibitor and vehicle-treated P0 cultures, and clone pools were re-plated without inhibitor, continuing this clonal analysis through secondary (2°), tertiary (3°) and quaternary (4°) assays (Fig. S1F). All inhibitor-treated colony types gave rise to large 2° colonies after re-plating in the absence of inhibitor - even micro-colonies, which, when taken from control cultures, do not replate (Fig. S1F). Further passage of clone pools restored colony forming ability to that typical of control colony types (Fig. S1F). Thus, PDGFRα inhibition does not appear to irreversibly commit cCFU-F cultures to differentiate and its effects on colony number are fully reversible over time. Consistently, bulk S^+^P^+^ cell cultures could be passaged in the presence of the inhibitor without exhaustion for >120 days (P13), albeit that inhibited cultures showed a lower rate of self-renewal than control cultures, whereas PDGF-AB ligand-treated cultures showed a higher rate of self-renewal (Fig. 1J). Therefore, PDGF-AB promotes, and PDGFRα inhibitor suppresses, colony formation and self-renewal.

### Short-term PDGF-AB treatment promotes heart repair

The specific impact of increasing or decreasing fibroblast signaling states *in vivo* is not known. Therefore, we asked how exogenous PDGF-AB influenced heart repair in a murine MI model in which left coronary artery was permanently occluded. A previous rat study delivered PDGF-AB via intra-myocardial injection 1-2 days prior to induction of MI, and improvements in cardiac parameters (scar size, ST-elevation, fractional shortening) and exercise tolerance were shown^30^, hypothesized to be via an angiogenic mechanism. However, the impact on fibroblasts was not considered, the model has no clinical correlate and no benefit was seen if PDGF-AB was injected at the time of MI^31^. Here, we delivered PDGF-AB via mini-pump to wild-type (WT) mice over a 5-day period from the time of MI, which led to a ~40% increase in serum PDGF-AB levels (Fig. S2A). At 28 days post-MI, PDGF-AB-treated hearts showed a ~50% reduction in scar area and normalized heart/body weight ratio (Fig. 2A-C). Echocardiographic analysis revealed a ~50% improvement in ejection fraction (EF) as early as day 5, a benefit maintained at day 28 post-MI, in PDGF-AB compared to PBS-treated hearts (Fig. 2D). Left ventricular (LV) fractional area change (FAC) was also improved at the ventricular apex, which, in MI+PBS hearts, was virtually akinetic (Fig. 2E,F; Fig. S2B). PDGF-AB treatment also significantly improved LV end systolic (ES) and end diastolic (ED) areas at days 5 and 28 post-MI at all four short-axis echocardiographic planes surveyed (Fig. 2G,H; Fig. S2C,D).

**Figure 2:**
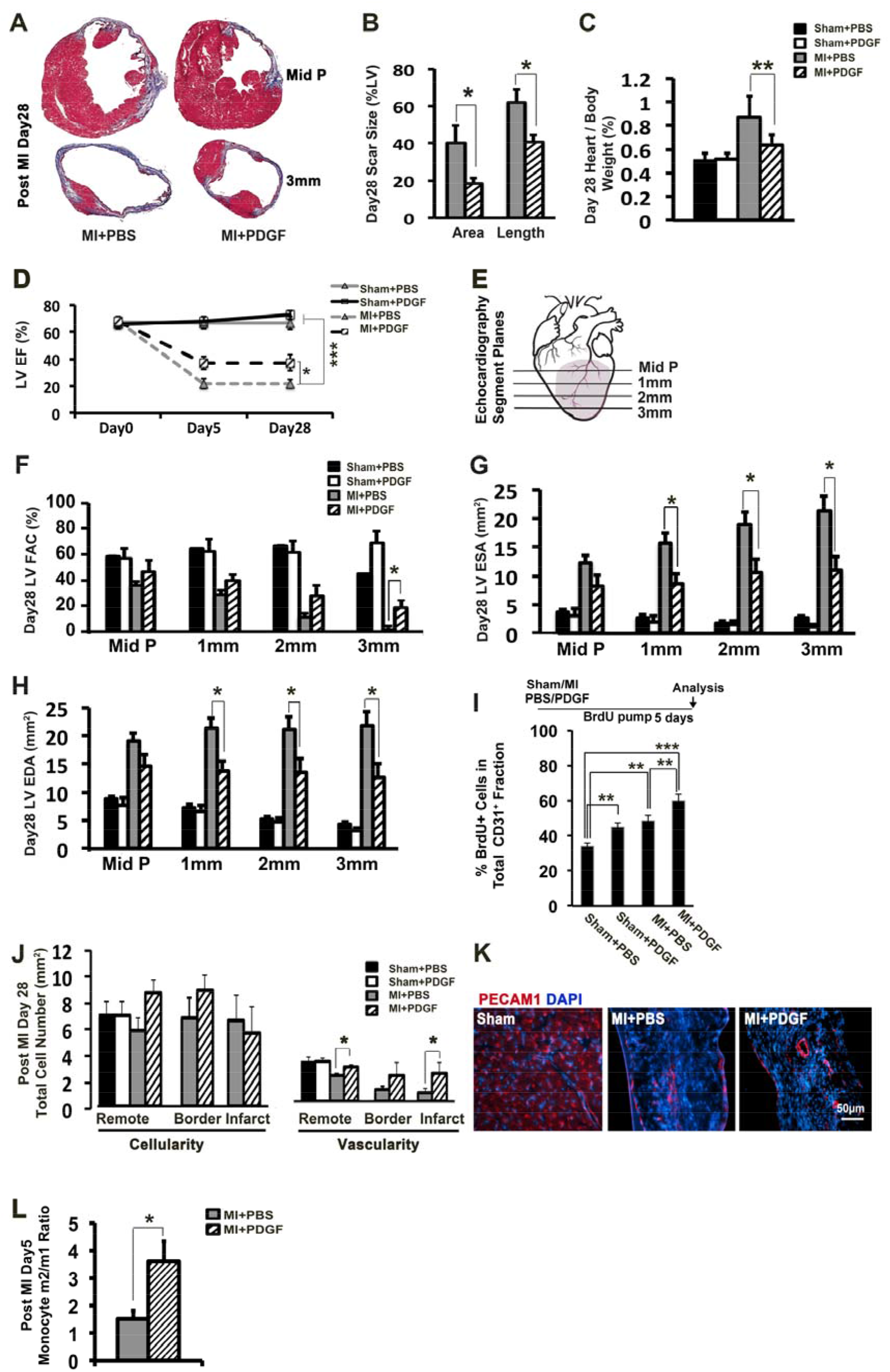
PDGF-AB enhancement of cardiac anatomical and function repair in MI hearts. (A) Distribution of scar (Masson’s Trichrome staining) in PBS and PDGF-AB-treated hearts 28 days post-MI at mid-papillary (Mid P) and 3 mm below Mid P levels (see Fig. 2E). (B) Scar area and length in PBS and PDGF-AB MI hearts. (C) Heart Weight/Body Weight ratio at day 28 sham/MI hearts. (D) LV ejection fraction (EF) in sham/MI hearts treated with PBS or PDGF-AB at 0 (baseline), 5 and 28 days. (E) Cartoon showing short axis planes for echocardiographic analyses. Mid P: mid-papillary. (F) Percent LV fractional area change (FAC) at different echocardiographic planes at day 28. (G,H) LV end systolic area (ESA) and diastolic area (EDA) respectively, at different echocardiographic planes at day 28. (I) BrdU uptake in CD31^+^ endothelial cells in sham/MI hearts with PBS or PDGF-AB treatments. (J) Total cellularity (DAPI-stained nuclei/mm^3^) and vascularity (CD31^+^ microvessels/mm^3^) in remote, border and infarct zones of day 28 sham/MI hearts. (K) Immunostaining showing increased caliber of CD31/PECAM1^+^ vessels in the infarct zone of PDGF-AB-treated day 28 post-MI hearts. (L) Quantification of flow cytometric analysis of m2/m1 monocyte ratios in PBS and PDGF-AB-treated MI hearts at day 5. All comparisons: * p<0.05; ** p<0.01; *** p<0.001.

We assayed dying cells (TUNEL assay^+^) at 6 hr post-MI and found no difference between PBS and PDGF-AB-treated hearts (Fig. S2E), suggesting that rescue of cell death in the immediate aftermath of MI is not the principal mechanism of benefit. We surveyed non-CM populations and found that PDGF-AB increased BrdU uptake in CD31^+^ vascular endothelial cells (ECs) at 5 days post-MI (Fig. 2I). This was also evident in PDGF-AB-treated sham hearts, showing that PDGF-AB can stimulate BrdU incorporation in ECs in the absence of injury^32^. Whereas there was no significant increase in vascular density in PDGF-AB-treated uninjured hearts, in treated MI hearts there was an increase in vascular density in remote as well as the border and infarct zones (Fig. 2J). Vessel caliber within the scar was increased at 28 days post-MI (Fig. 2K).

We also looked at monocytes and found that the relative number of pro-inflammatory m1 monocytes was decreased at day-5 (Fig. S2F,G), leading to an increase in the Ly-6C^low^/Ly-6C^high^ (m2/m1) monocyte ratio (Fig. 2L) known to reflect a pro-reparative monocyte milieu^4^. Thus, PDGF-AB can significantly improve cardiac functional repair when delivered from the time of MI, potentially impacting fibroblasts and multiple other cell types directly or indirectly.

### Pdgfra-GFP discriminates between resting and activated fibroblasts in vivo

To begin to dissect the mechanism of enhanced functional repair, we focused for the remaining parts of this study on PDGFRα^+^ cardiac fibroblasts as a direct target of PDGF-AB. We first investigated *Pdgfra^nGFP/+^* knockin mice as a new tool for discriminating between resting and activated PDGFRα^+^ fibroblasts *in vivo*^11^. In sham hearts, virtually all S^+^P^+^ and S^−^P^+^ fibroblasts were GFP^high^, as assessed by flow cytometry (Fig. 3A, left panels). On tissue sections, GFP^bright^ cells were found embedded within a collagen VI-rich matrix in the *tunica adventia* of arterioles and distributed throughout the myocardial interstitium close to isolectin^+^ microvessels (Fig. S3A-D), consistent with previous studies on fibroblast localization^7^. However, neither CMs nor ECs showed detectable GFP (see Discussion). GFP^bright^ cells were low or negative for PDGFRβ and other pericyte markers including CD146 and NG2 (Fig. S3E-G; I), and were exclusively external to the collagen IV^+^ basal lamina surrounding vascular endothelial cells (Fig. S3H; 1500 cells scored), indicating that they are not pericytes^27^.

**Figure 3:**
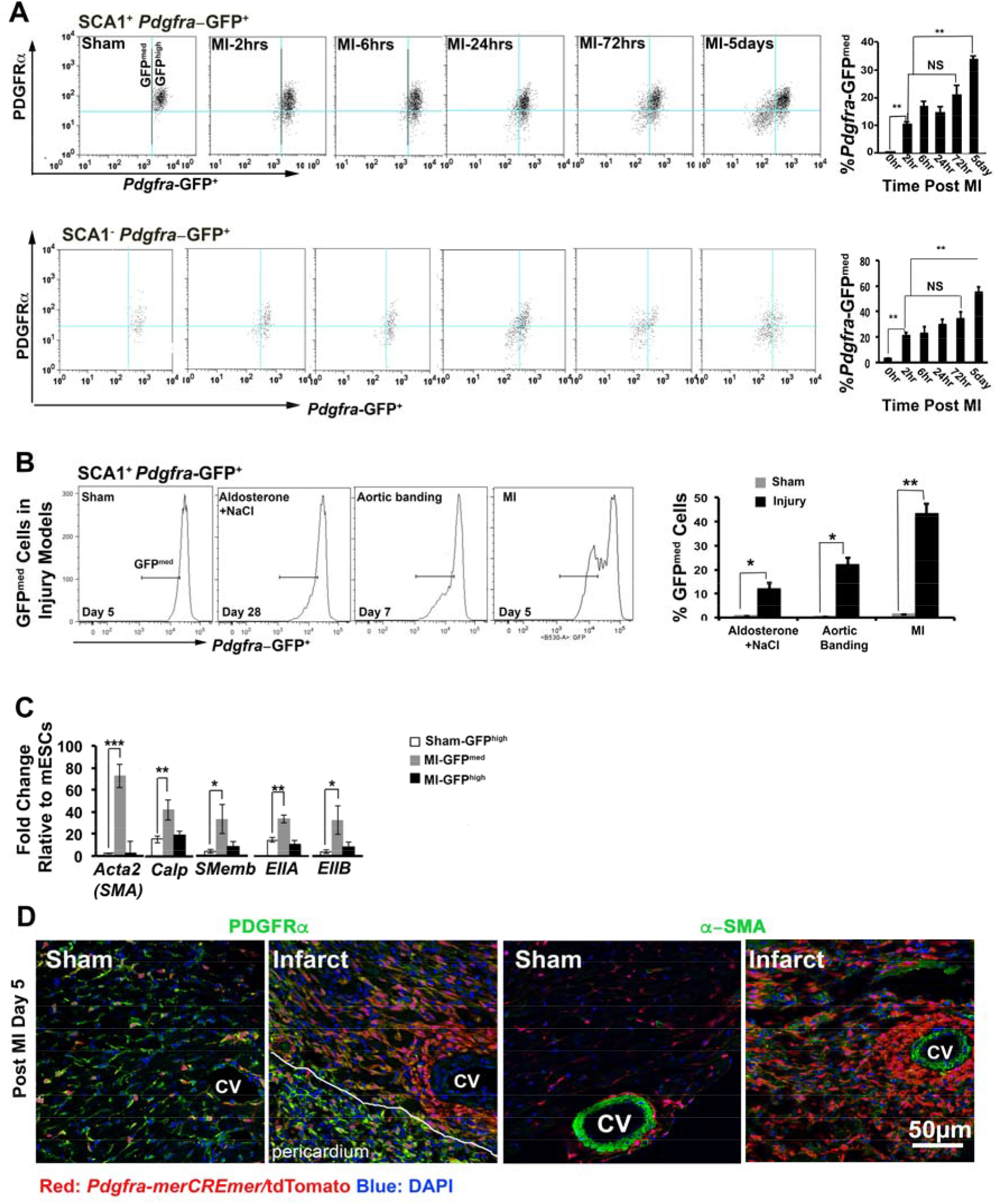
Dynamics of *Pdgfra*-GFP^medium^ myofibroblasts after MI. (A) Time course flow cytometry analysis of SCA1^+^/*Pdgfra*-GFP^+^ and SCA1^−^/*Pdgfra*-GFP^+^ fibroblasts, displaying PDGFRα and GFP expression, with quantification of GFP^medium^ myofibroblasts. (B) Flow cytometry showing GFP^medium^ cell population (bar) in SCA1^+^/*Pdgfra*-GFP^+^ fibroblasts from *Pdgfra*^GFP/+^ mice subjected to sham operation and disease models including aldosterone plus high salt treatment, aortic banding and MI, with graph showing percentage of the SCA1^+^*Pdgfra*-GFP^medium^ quantified in respective assays. (C) qRT-PCR analysis of activated fibroblast markers in total GFP^medium^ and GFP^high^ cells at 5 days post-sham or MI surgeries. (D) Lineage tracing of PDGFR^+^ fibroblasts at 5 days post-sham/MI using tamoxifen-treated progenies of *Pdgfra^merCREmer/+^* crossed to *Rosa26^tdTomato/+^* reporter mice. Panels display endogenous fluorescence of tdTomato-Red, and immunofluorescence staining of PDGFRα and SMA. They highlight tdTomato^+^ cells brightly stained for PDGFRα but SMA^−^ in the sham hearts and tdTomato^+^PDGFRα^dull^SMA^+^ cells in the infarct area. cv: coronary vessel. All comparisons: * p<0.05; ** p<0.01; *** p<0.001.

Post-MI, flow cytometry revealed accumulation of a GFP^medium^ population in *Pdgfra*-GFP hearts between 2 hrs and 5 days (Fig. 3A). These were negative for the hematopoietic lineage cell surface marker CD45 (Fig. S3J), consistent with a local origin, and by day 5 post-MI most had become negative for cell surface PDGFRα (Fig. 3A). S^+^GFP^medium^ cells were also seen when cardiac fibrosis was induced by aldosterone/salt treatment or aortic banding (Fig. 3B). We determined that S^+^GFP^medium^ cells encompassed activated fibroblast populations. We recently profiled cardiac fibroblasts in sham and days 3 and 7 post-MI hearts using scRNAseq data^16^ and observed down-regulation of *GFP, Pdgfra* and other stem cell markers (*Sca1, Cd34, Cd90/Thy1*) with injury, and up-regulation of contractile markers such as *Acta2* (encoding αSMA) and the matricellular marker *Postn* (encoding Periostin), in fibroblasts as they activate, proliferate and differentiate after MI^16^. scRNAseq data allowed us to formally define the main population of activated fibroblasts (F-Act), a novel activated fibroblast population (F-WntX) present in sham hearts, a distinct pre-proliferative state (F-CI) and proliferating fibroblasts (F-Cyc) which predominate at MI-day 3, and multiple myofibroblast populations at MI-day 7. qRT-PCR confirmed higher expression of activation markers in S^+^GFP^medium^ cells relative to S^+^GFP^high^ cells at 5 days post-MI, including a ~70-fold increase in expression of *Acta2* and down-regulation of several cardiac transcription factor genes that mark resting fibroblasts^33^ (Fig. 3C; Fig. S3K). When plated, GFP^high^ cells gave rise to large cCFU-F colonies which were PDGFRα^bright^αSMA^−^, whereas GFP^medium^ cells only gave rise to micro-colonies which were PDGFRα^dim^αSMA^+^ (Fig. S3L,M).

We confirmed that resting *Pdgfra*^+^ fibroblasts give rise to activated fibroblasts by lineage tracing using tamoxifen (tam)-inducible *pdgfra^merCremer/+^* CRE recombinase driver mice crossed to a *Rosa26^TdTomato^* CRE reporter mice (Fig. 3D). 14 days after tam treatment and 5 days post-sham, virtually all lineage tagged (tdTomato^+^) cells were PDGFRα^bright^αSMA^−^, as expected of resting fibroblasts. Post-MI, the infarct region contained abundant, large, tdTomato^+^PDGFRα^dim^αSMA^high^ cells, findings consistent with recent fibroblast lineage tracing studies^34–36^.

### Flux between resting and activated fibroblasts in PDGF-AB-treated MI hearts

We quantified GFP^high^ and GFP^medium^ fractions at day 5-post sham or MI. A GFP^medium^ population was not detected in the absence of MI (Fig. 3A; Fig. 4A). Consistent with time-course evaluation (Fig. 3A), GFP^medium^ cells accumulated in MI+PBS animals (Fig. 4A), and this could be partially inhibited by treatment with the PDGFRα/β signaling antagonist Imatinib (Fig. 4A), or systemic injection of blocking anti-PDGFRα antibody (Fig. 4B), demonstrating that normal fibroblast activation *in vivo* is dependent at least in part on endogenous PDGFRα signaling.

**Figure 4:**
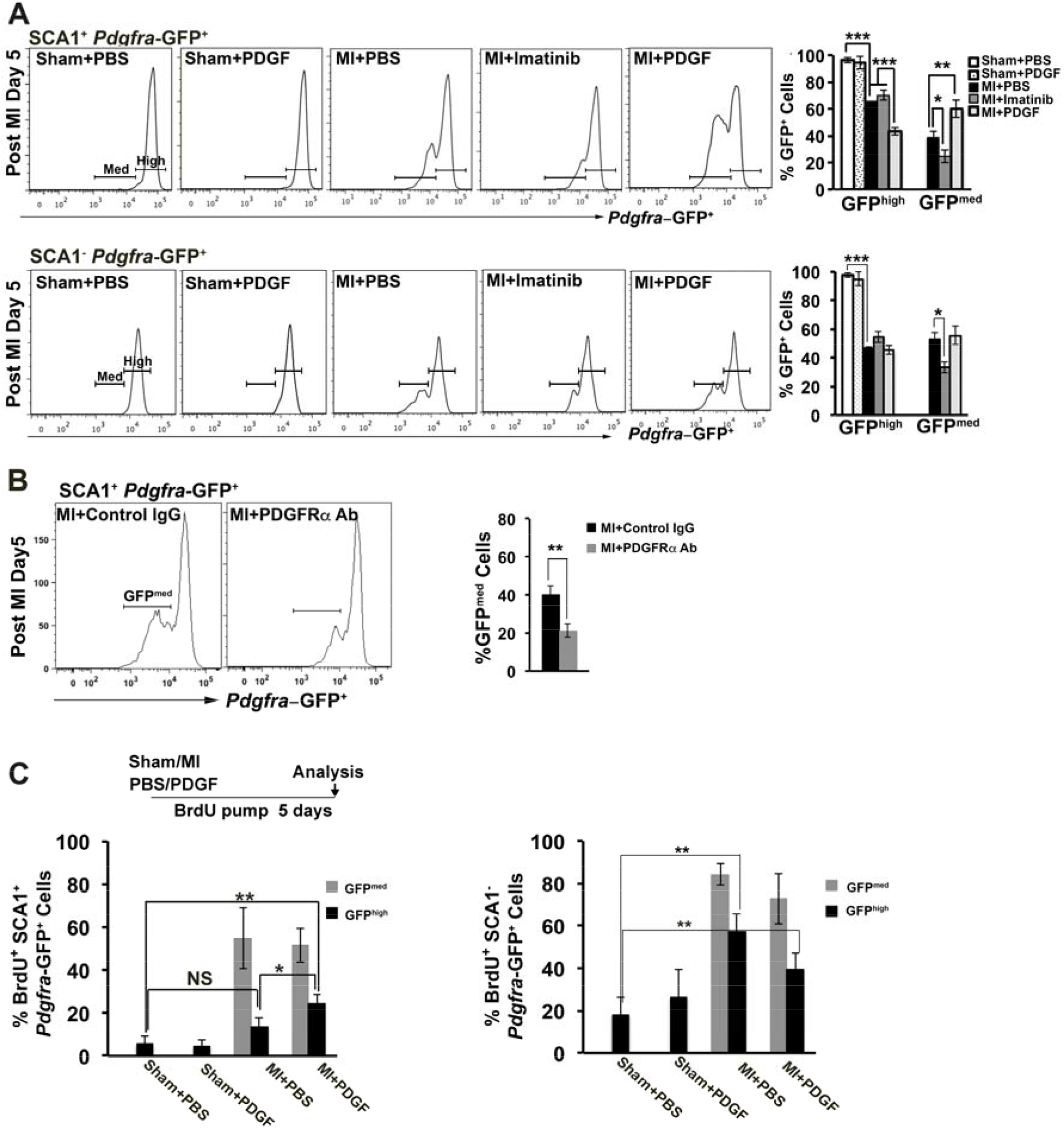
Analysis of proliferation and cell flux in the fibroblast lineage in sham/MI hearts with PDGF-AB and Imatinib treatments. (A) Flow cytometry quantifying GFP^high^ fibroblasts (High) and GFP^medium^ myofibroblasts (Med) from day 5 post-sham/MI hearts treated with PBS, PDGF-AB or Imatinib in both SCA1^+^ and SCA1^−^ fibroblast sub-fractions. (B) Flow cytometric analysis of SCA1^+^*Pdgfra*^−^GFP^+^ cardiac fibroblasts at day 5 post-MI after treatment with control IgG antibody or blocking with monoclonal PDGFRα antibody (Ab), with quantification of GFP^medium^ myofibroblasts (bar). (C) Accumulated BrdU incorporation in GFP^high^ fibroblasts and GFP^medium^ myofibroblasts in both SCA1^+^ and SCA1^−^ populations at 5 days post-sham/MI after treatment with PBS or PDGF-AB. All comparisons: * p<0.05; ** p<0.01; *** p<0.001.

We noted that activated GFP^medium^ cells formed at the expense of resting GFP^high^ fibroblasts, and this was evident in both SCA1^+^ and SCA1^−^ sub-fractions (see bar plots in Fig. 4A). This is consistent with the currently-held view that activated fibroblasts derive almost exclusively from resident resting fibroblasts^37^. This effect was also observed in fibroblast populations defined by scRNAseq data^16^. Based on our flow data, 30-50% of resting fibroblasts transit to an activated (GFP^medium^) state by MI-day 5.

PDGF-AB treatment of uninjured hearts did not lead to activation of fibroblasts (Fig. 4A). Strikingly, however, PDGF-AB treatment increased the number of S^+^GFP^medium^ fibroblasts by ~50% at MI-day 5. Interestingly, S^−^GFP^medium^ cells were not increased, even though Imatinib inhibited their accumulation post-MI. Thus, PDGF-AB targets S^+^P^+^ cells and increases the activated fibroblast pool after injury.

To assess proliferation in the different fibroblast populations over the first 5 days, BrdU was delivered (along with vehicle, PDGF-AB or inhibitor) continuously from the time of MI via mini-pump (Fig. 4C). A basal level of BrdU incorporation was seen in GFP^high^ cells in the sham condition (higher in the S^−^GFP^high^ population); however, PDGF-AB did not stimulate this level, suggesting that exogenous PDGF-AB is not mitogenic for fibroblasts in uninjured hearts. After MI, BrdU incorporation was increased in GFP^high^ cells and PDGF-AB slightly stimulated incorporation into the S^+^GFP^high^ population. We were surprised by this finding, given that scRNAseq clearly assigns pre-proliferative and proliferative fibroblasts an activated status (high levels of *Acta2* and *Postn*, and lower levels of *Pdgfra* and *Sca1*, relative to resting fibroblasts)^16^. It is possible that some proliferating fibroblasts retain high GFP expression due to the stability of H2B-eGFP compared to PDGFRα protein; alternatively, some activated and divided fibroblast may regress to a resting (GFP^high^) state^16,38^.

GFP^medium^ cells showed higher BrdU incorporation than GFP^high^ cells, consistent with the known division of activated fibroblasts in response to injury-induced mitogens over the first 2-4 days post-MI^16,34^ (Fig. 4C). However, PDGF-AB did not stimulate an increase in BrdU incorporation in GFP^medium^ cells, demonstrating that exogenous PDGF-AB does not act as a conventional mitogen for activated fibroblast lineage cells. Thus, PDGF-AB promotes activation of cardiac fibroblasts without stimulating DNA synthesis, consistent with our *in vitro* findings (Fig. 1A-D).

### Biosythetic profiling of cardiac fibroblasts

Taking an alternative approach to profiling fibroblast activation, we stained freshly isolated S^+^P^+^ cells with DNA (7AAD) and RNA (Pyronin Y) dyes and analyzed uptake by flow cytometry. Pyronin Y staining largely reflects ribosomal RNA and ribosomal protein mRNA content, and serves as a marker of the biosynthetic state^39^. In uninjured and sham-operated hearts, virtually all freshly isolated S^+^P^+^ cells had 2n DNA content, with 60% exhibiting a very low RNA state consistent with quiescence (G_0_) (Fig. 5A). The remainder showed a continuum of higher biosynthetic states. By day 5 post-MI (after the peak in fibroblast proliferation^34,35^), there was a significant reduction in the proportion of cells in G_0_ from ~60% to ~40%, showing that a large number of quiescent fibroblasts become biosynthetically activated after MI (Fig. 5A). Activation was completely inhibited by Imatinib treatment, demonstrating again that normal fibroblast activation *in vivo* requires PDGFR signaling.

**Figure 5:**
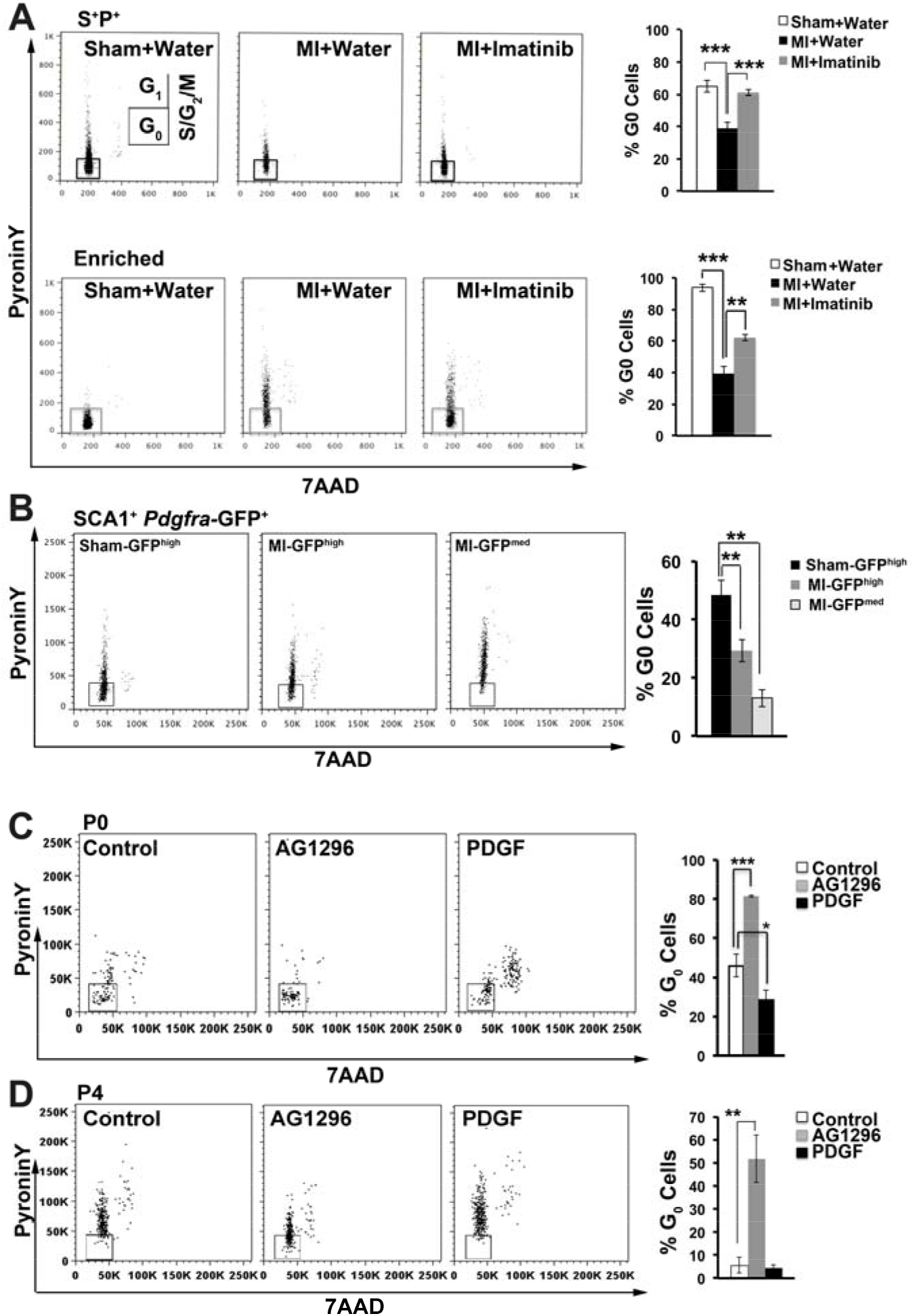
Cell cycle analyses of fresh and cultured fibroblasts after PDGFRα ligand and inhibitor treatments. (A) PyroninY/7AAD cell cycle analyses of freshly isolated S^+^P^+^ (top panels) and colony-enriched fibroblasts (bottom panels) at day 5 post-Sham/MI. Insert shows cell cycle compartments. Data also shown from MI mice treated with Imatinib or water control. Percent G_0_ cells (boxed) indicated. (B) Cell cycle analysis of S^+^*Pdgfra*-GFP^+^ fibroblasts (flow-purified GFP^high^ and GFP^medium^ fractions) from *Pdgfra*^GFP/+^ mice subject to sham/MI. Graph indicates percent G_0_ cells (boxed). (C) Cell cycle analyses of freshly plated S^+^P^+^ cultures with and without PDGF-AB and AG1296 at P0. (D) Cell cycle analysis of P4 S^+^P^+^ cultures with and without AG1296 or PDGF-AB. All comparisons: * p<0.05; ** p<0.01; *** p<0.001.

PyroninY/7AAD analysis of fibroblasts from *Pdgfra*-GFP mice revealed similar heterogeneity in biosynthetic states among SCA1^+^GFP^high^ cells (Fig. 5B). S^+^GFP^medium^ cells also showed a broad biosynthetic spectrum, although, as expected for activated cells, very few (~10%) were in G_0_ and overall there was a shift to higher states in these cells.

To examine the biosynthetic state of cCFU-F, S^+^P^+^ cells were further fractionated to CD90^+^PDGFRβ^low^ cells, which resulted in a >10-fold enrichment in large colony-forming cells^33^. Virtually all cells in this enriched fraction were in G_0_ (Fig. 5A, lower panels); however, >50% became activated after MI, which was partially inhibited by Imatinib. Thus, similar to long-lived stem cells, cCFU-F reside in G_0_, with a major shift towards higher biosynthetic states as they become activated after MI.

We applied the PyroninY/7AAD assay to cCFU-F cultures. When freshly isolated S^+^P^+^ cells were cultured until the end of P0, ~46% remained in G_0_ (Fig. 5C). Because 60% of P0 cells divide after plating (Fig. 1A), many of these dividing cells must retain or re-assume a low biosynthetic state. This delayed adaption to cell culture likely accounts for the slower initial rate of self-renewal of cCFU-F cultures, which show an inflection point at ~P4 (48 days)^11^ (Figs. 1F; see also Fig. 6I). This is consistent with the slow activation of stem cells from quiescence^40^, although we cannot discount a contribution from cell stress during adaptation to cell culture. Strikingly, however, cultures treated with PDGF-AB ligand across P0 showed a decrease in the proportion of cells in G_0_ from ~46% to ~29%, whereas AG1296 increased the proportion of cells in G_0_ from ~46% to ~81% (Fig. 5C). Even in P4 cultures, in which virtually all cells were fully activated (RNA^medium-high^), treatment with AG1296 led to ~50% of cells being drawn into G_0_ (Fig. 5D), with an associated reduction in cell yield (Fig. 1H). Thus, in cCFU-F cultures, PDGFRα signaling promotes exit from quiescence. Conversely, inhibition of PDGFRα signaling induces quiescence. These results align with our observation that PDGFRα signaling promotes exit of freshly-plated fibroblasts from quiescence *in vitro*.

**Figure 6:**
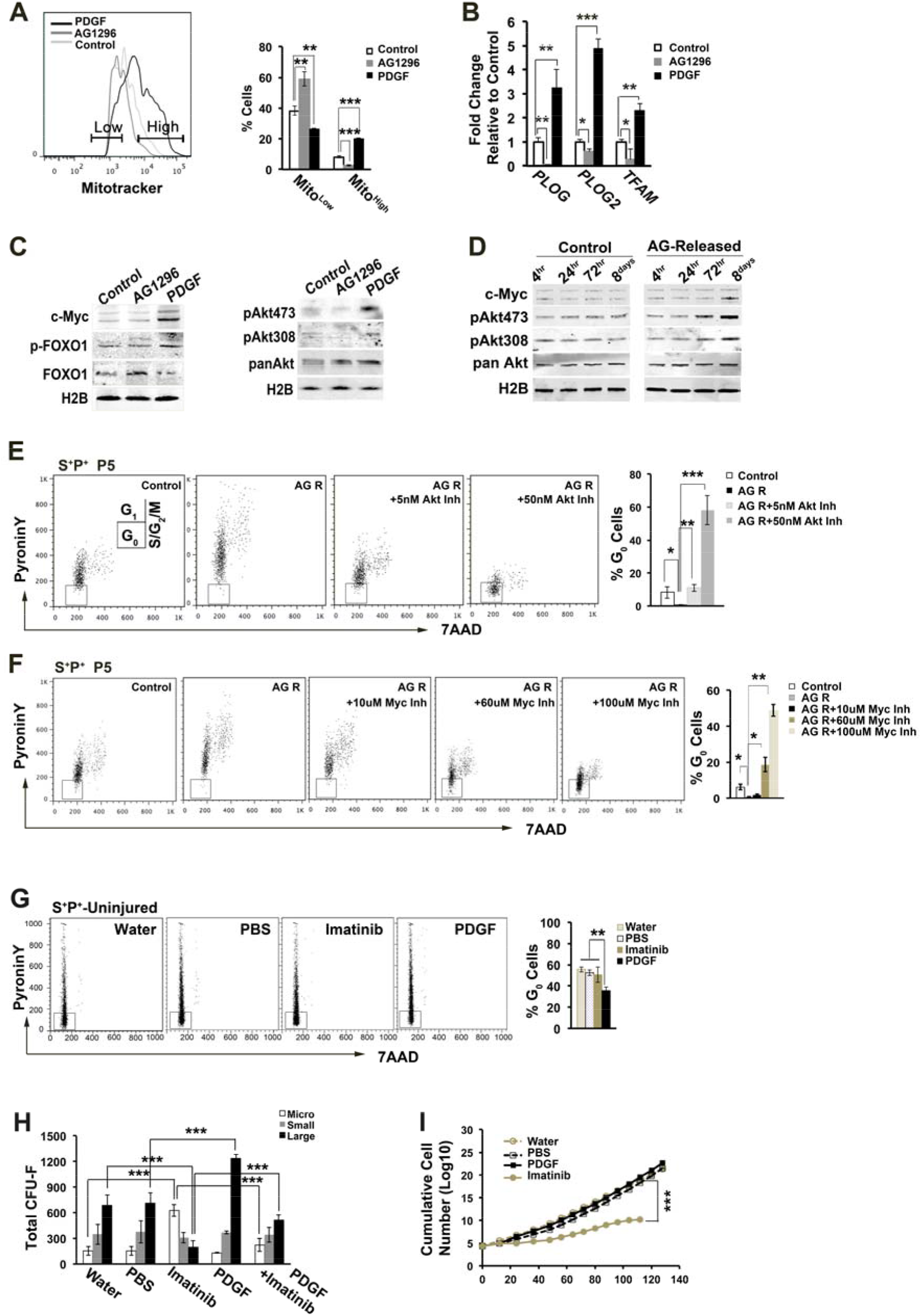
Metabolic and cell cycle states of S^+^P^+^ cultures – essential role of AKT and cMYC in cell cycle reentry after PDGFR inhibition. (A) Quantification of mitochondrial mass in S^+^P^+^ cells cultured to P0 by Mitotracker® Green FM (Mito) staining, with or without PDGF-AB (100ng/ml) and AG1296 (10μM), showing quantification of Mito^low^ and Mito^high^ fractions. (B) qRT-PCR analysis of genes for mitochondrial polymerase enzymes (POLG; POLG2) and transcription factor (TFAM). (C,D) Western blot analyses of indicated cMYC and AKT pathway proteins in PDGF-AB and AG1296-treated S^+^P^+^ cultures at P0 (C), and after AG1296 inhibition across P4 and release from inhibition for a further passage (AG-Released; D). (E,F) Cell cycle analysis of uninhibited cultures (P5) compared to cultures treated across P4 with AG1296 and released (AG R), with and without increasing doses of AKT and cMYC inhibitors. Graphs in E and F show proportion of cells in G_0_. (G) Cell cycle analysis after *in vivo* treatment of uninjured mice with Imatinib and/or PDGF-AB ligand relative to vehicle controls at day5. (H) Total cCFU-F (micro-, small, large) per heart from mice receiving PDGF-AB and/or Imatinib treatments (I) Long-term growth curves of S^+^P^+^ cultures after *in vivo* treatments. All comparisons: * p<0.05; ** p<0.01; *** p<0.001.

These data complement our analysis of fibroblast activation using *Pdgfra*-GFP mice. They highlight the significant heterogeneity in the biosynthetic state of ventricular cardiac fibroblasts even at rest, with many cells, including cCFU-F, being in G_0_ (quiescence), whereas others, while dormant, exhibit a spectrum of biosynthetic states. PDGF signaling stimulates exit from quiescence, whereas inhibition induces quiescence.

### Exit from PDGFRα-inhibition-induced quiescence requires AKT and cMYC

To see whether the biosynthetic changes in fibroblasts during activation correlate with metabolic indicators, we probed mitochondrial and metabolic signaling pathways in cultured S^+^P^+^ cells. Using MitoTracker Green staining, we found that PDGF-AB increased the proportion of cells with higher mitochondrial mass, whereas AG1296 increased the proportion of cells with a lower mass (Fig. 6A). These effects were accompanied by changes in transcripts encoding mitochondrial polymerases (POLG, POLG2) and the transcription factor, TFAM (Fig. 6B). Thus, increased biosynthetic activity induced by PDGF-AB drives mitochondrial biosynthetic networks, with likely changes in energy substrate use.

PDGFRα signaling is known to activate the PI3K/AKT/FOXO and c-Myc pathways involved in G1 checkpoint control, cell growth, mitochondrial biosynthesis and stem cell function^41–44^. Important to this study, nuclear AKT and cMYC collaborate in ribosomal biogenesis^45^. Treatment of cCFU-F cultures with PDGF-AB until the end of P0 activated the AKT and cMYC pathways, as seen by increases in the active phosphorylated forms of AKT (pAKT473, pAKT308) and in AKT-dependent phosphorylation of FOXO (which excludes it from the nucleus and marks it for degradation), and increased cMYC levels (Fig. 6C). Conversely, AG1296 led to a decrease in pAKT and pFOXO, and an increase in total FOXO, consistent with the observed induction of quiescence^42^.

We discovered using the Pyronin Y/7AAD assay that if inhibitor-treated P4 cultures (~60% of cells in G_0_) were released from inhibitor for a further passage (P5) there was a significant upwards surge in biosynthetic state and only ~1% of cells remained in G_0_ (condition AG-R versus control; Fig. 6E; Fig. S4A). This may be due to the increased expression of *Pdgfra* with inhibitor (data not shown). Time course Western blot analysis of AG-R cultures across P5 showed accumulation of pAKT and cMYC (Fig. 6D). We exploited this assay to ask whether the AKT and cMYC pathways were required for exit from the G_0_-like state imposed by PDGFR inhibition. At the concentrations used, neither the AKT inhibitor (LY294002) nor cMYC inhibitor (10058-F4) had any impact on steady state growth or the PyroninY/7AAD profile of control cultures (Fig. S4B,C and data not shown). However, exit from the G_0_-like state induced by PDGFR inhibition was completely blocked in a dose-dependent manner by both drugs (Fig. 6E,F). Thus, PDGFR signaling in cCFU-F cultures governs metabolic and biosynthetic states, with inhibition inducing a low biosynthetic and network state resembling G_0_/quiescence, release from which requires AKT and cMYC.

### Transcriptome analysis of cCFU-F cultures

Microarray and principal component analysis of cCFU-F cultures at P0 showed that inhibitor-treated cells were transcriptionally distinct from those of untreated and ligand-treated cultures, which clustered together (Fig. S5A). The top gene ontology (GO) terms associated with genes downregulated by inhibitor were *nucleosome assembly* (driven largely by down-regulation of histone genes), as well as *epigenetic regulation of gene expression, DNA methylation, cell proliferation, migration* and *adhesion, negative regulation of apoptosis, angiogenesis* and other signaling terms (Fig. S5B), consistent with a repressed signaling, proliferative, transcriptional and epigenetic state in inhibitor-treated cultures.

There were many genes up-regulated with inhibitor, associated with terms for *cell adhesion* and *extracellular matrix*, as well as *negative regulation of Wnt signaling* (a positive driver of fibrosis) and a variety of tissue developmental processes (Fig. S6B), consistent with findings that quiescence involves many active network states controlling metabolism, proliferation and differentiation^46^.

### Impact of modulating PDGFRα signaling on cCFU-F in uninjured hearts

Although *in vivo* delivery of PDGF-AB did not activate cardiac fibroblasts in the absence of injury using *Pdgfra*-GFP as the readout (Fig. 4A), PDGF-AB may induce metabolic and biosynthetic changes in resting fibroblasts that could relate to the increased fibroblast activation seen during MI. We specifically assessed this using PyroninY/7AAD staining. Whereas Imatinib treatment of uninjured mice for 5 days had no effect on the staining profile of isolated S^+^P^+^ fibroblasts, *in vivo* PDGF-AB treatment shifted the profile significantly, inducing exit of ~35% of cells from G_0_ (Fig. 6G). We also assayed colony formation after *in vivo* treatments. PDGF-AB induced a ~70% increase in large colony formation from S^+^P^+^ cells of uninjured hearts, whereas Imatinib severely inhibited colony numbers, with PDGF-AB partially mitigating the effects of Imatinib (Fig. 6H). Imatinib significantly inhibited long-term self-renewal of cCFU-F cultures (assayed in the absence of inhibitor; Fig. 6I). Identical results were seen after injection of blocking PDGFRα mAb to uninjured mice (Fig. S4D,E). These colony and long-term growth profiles closely resembled those obtained after ligand and inhibitor treatment of S^+^P^+^ cultures *in vitro* (Fig. 1I,J). Our data show that PDGF-AB can metabolically prime cardiac fibroblasts *in vivo* independently of injury.

### Single cell transcriptome analysis of cardiac fibroblasts after in vivo PDGF treatment

It was important to establish whether fibroblast activation occurring in PDGF-AB-treated sham and MI hearts was a trigger for myofibroblast differentiation and fibrosis. scRNAseq can significantly extend our view of cell state and lineage heterogeneity in complex biological systems. We therefore performed scRNAseq on the total cardiac interstitial cell population (TIP) of hearts at days 3 and 7 post-sham or MI surgery, with PBS or PDGF-AB treatment. Days 3 and 7 were chosen to align with our previous scRNAseq study^16^. We profiled over 45,758 cells after quality control filtering and performed unbiased clustering using the Seurat R package^47^, identifying 24 sub-populations after removal of hybrid or contaminating (erythroid) cells, with data plotted on t-SNE dimensionality reduction plots (Fig. 7A)^16^. All major TIP populations corresponded to those we previously characterized at days 3 and 7 post-sham and MI^16^; however, we did not observe any major population-level differences between the PBS and PDGF conditions at MI-days 3 or 7 (Fig. S6A), confirmed by Differential Proportion Analysis (DPA) (Fig. S6D)^16^.

**Figure 7:**
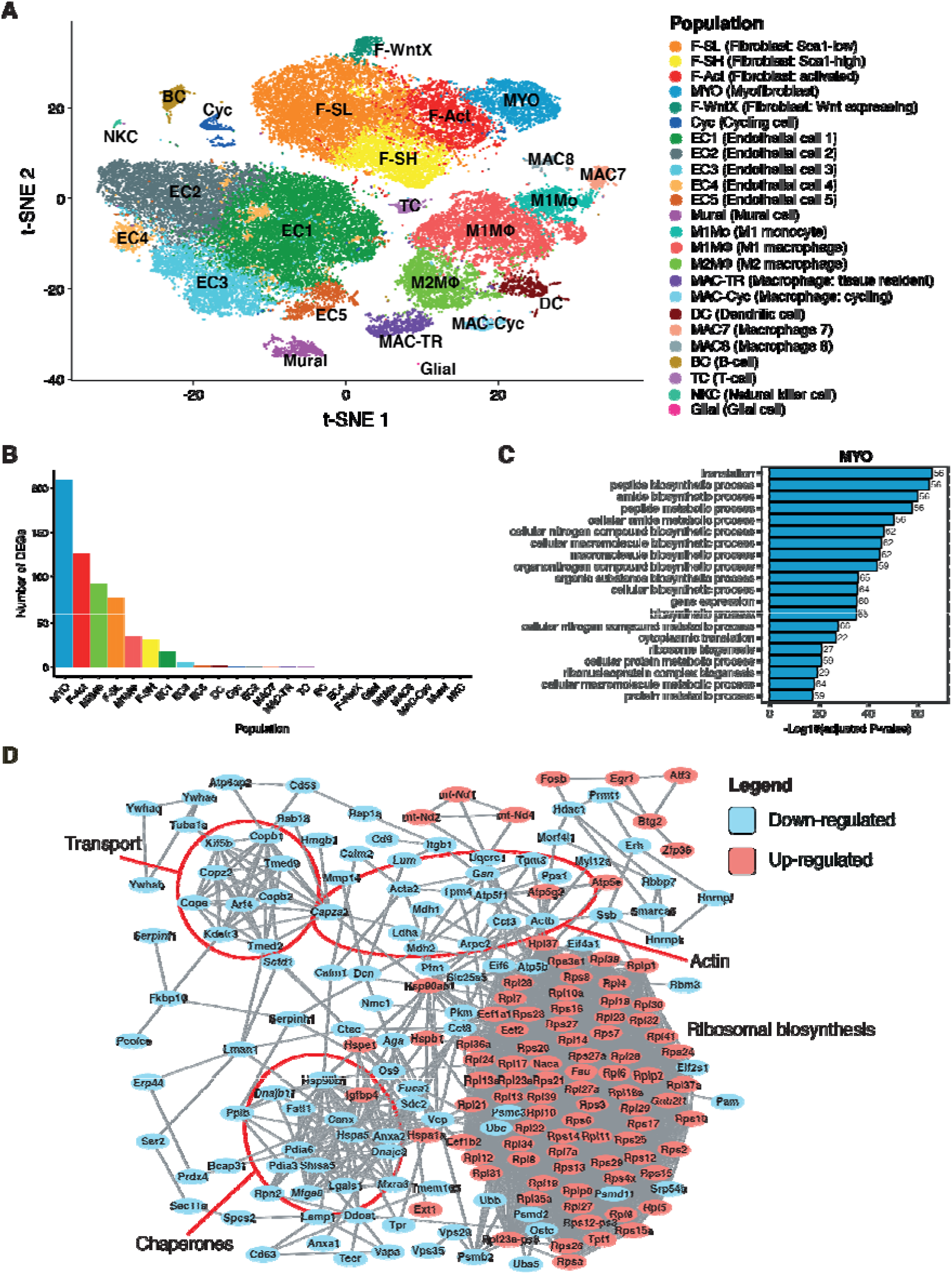
Single-cell RNA-seq of total interstitial population (TIP) cells from PDGF-AB and PBS treated, injured vs. uninjured hearts. (A) t-SNE plot of aggregate of cells across conditions showing cells coloured according to identified populations. (B) Number of population-specific differentially expressed genes (DEGs) between PDGF-AB and PBS conditions at MI-day 7 (MAST, P_adj_ < 1e-05, log2 fold-change > 0.25). (C) Top 20 over-represented GO biological process terms for DEGs up-regulated in MYO at MI-day 7 with a PDGF-AB vs. PBS comparison. (D) STRING high-confidence network of differentially expressed genes between PDGF-AB and PBS conditions at MI-day 7 within sub-populations MYO, F-Act, M2MΦ and F-SL. Genes (nodes) are coloured according to up-regulated in PDGF-AB (red) or down-regulated (blue).

Nonetheless, we noted a slight but significant increase in the number of unique molecular identifiers (UMIs) in PDGF-AB-treated TIP cells across all experiments (Wilcoxon rank-sum test; P < 1e-07 for all experiments), including in PDGF-AB-treated sham hearts (Fig. S6C). There was a similar trend for the number of genes detected per-cell (with the exception of MI-day 7) (Fig. S6D), overall suggesting an increased level of transcription in TIP cells from PDGF-AB treatment.

Differential expression (DE) analysis using MAST^48^ was applied to detect population-specific DE genes between PDGF-AB and PBS conditions. The largest number of DE genes were detected at MI-day 7 (Fig. 7B; Fig. S7A-D) and myofibroblasts (MYO) showed the largest number of DE genes (>200; Fig. 7B). The number of DE genes decreased progressively in activated fibroblasts (F-Act), M2 macrophages (M2MΦ), *Sca1*^low^ resting fibroblasts (F-SL) and other populations. The change in gene expression between PBS and PDGF-AB conditions was relatively modest, with most genes showing a 0.25-0.5 (log2) fold-change. Nevertheless, gene ontology (GO) over-representation testing with PANTHER^49^ showed that PDGF-AB up-regulated genes were overwhelmingly associated with components of the translational machinery (Fig. 7C; Fig. S7E-G; Supplementary Tables 1-4). Protein:protein association network analysis using STRING^50^ showed that up-regulated DE genes encoded a tightly interconnected network of ribosomal and ribosome-connected proteins, the latter including numerous heat shock protein chaperones, Gnb2l1 (involved in ribosome quality control), translation initiation factors Eif1 (involved in ribosome biosynthesis), Eif2s1 and Eif4a1, translation elongation factors 1α and 1β, nascent polypeptide-associated complex protein alpha (Naca), which inhibits nascent proteins from being inappropriately shunted to the endoplasmic reticulum, and Ostc, a subunit of an oligosaccharyltransferase which transfers high manose oligosaccharides to nascent proteins (Fig. 7D). Complex I and V mitochondrial protein genes mt-Nd1/2/4 were also up-regulated. The vast preponderance of ribosomal protein genes in the network is highly suggestive of activated PI3K/AKT/mTOR signaling and c-Myc activity, which are the main pathways collaborating to regulate ribosomal biosynthesis^45^. Ribosomal RNA synthesis is likely also increased by PDGF-AB, as shown by Pyronin Y staining in uninjured hearts (Fig. 6G), although this would not be detected in our scRNA-seq study since this was performed on polyadenylated RNA. The increased expression of transcription factor genes *Klf4, Nfix, Egr1, Fosb, Atf3* and *Zbtb20* may account for the increased global transcription in PDGF-AB-treated hearts.

Collagen genes were not up-regulated by PDGF-AB and indeed most DE genes related to ECM were down-regulated in MYO (Fig. S7H), suggesting that PDGF-AB does not trigger activated fibroblast lineage cells to differentiate into myofibroblasts. Other down-regulated gene modules included those for COPI retrograde golgi-to-endoplasmic reticulum transport (e.g. *Copb1, Copb2, Copx2, Cope, Arf4, Scdf1*), actin and actin filament regulatory proteins (e.g. *Acta2, Actb, Tpm3/4, Gsn, Arpc2, Capza2*), and a distinct group of molecular chaperones (e.g. *Hspa5, Hsp90b1, Dnajb11, Dnajc3, Pdia3/6*). Overall, the signature for PDGF-AB primed fibroblast lineage cells is one of elevated protein synthesis capacity, seen most strongly in MYO, as well as other fibroblast and macrophage populations. There was no evidence that activated cells possessed an enhanced ECM or general secretory phenotype, or had differentiated to myofibroblasts disproportionately.

## DISCUSSION

Replacing CMs lost to ischemic injury will be necessary for achieving heart regeneration in humans. However, it is intuitive that the diverse connective tissue and vascular populations of the heart, which form a complex interactive cellular community contributing to organ architecture, viability and function, will also need to be restored^38^. In pro-regenerative animals, fibroblasts, vessels, immune cells and nerves are all essential for heart regeneration^51^. Therefore, characterizing the heterogeneity in these cells, and understanding and unlocking their paracrine signaling and pro-regenerative functions, is a research and therapeutic priority.

PDGF ligands are expressed by many tissues involved in cardiac injury and repair. Virtually all resting fibroblasts express the PDGFRα gene and protein, making them a likely target of cognate ligands^16,17^. The *Pdgfrb* gene is also expressed in cardiac fibroblast lineage cells in healthy and injured adult hearts, including vascular mural cells^16^, although we found only low to absent levels of PDGFRβ protein in *Pdgfra*-GFP^high^ resting fibroblasts by immunofluorescence (Fig. S3G). This is likely because PDGFRβ acts downstream of PDGFRα principally in recruitment of perivascular cells in remodeling vascular networks^18^, although in adult hearts it is also expressed on CMs under stress conditions where it may elicit pro-angiogenic effects^19,52^.

PDGF ligands have been implicated as mitogens for mesenchymal cells and MSCs, and chronic activation of PDGFR signaling has been associated with fibrosis and cancer in different settings^22^. However, various PDGFs have also been tested for their impact on heart injury based on a pro-angiogenic rationale. Different ligands and delivery approaches have led to diverse outcomes, from improved heart repair after MI, to frank fibrosis and heart failure in uninjured mice^22,24,27,28^. Only one study has assessed the impact of PDGF-AB, which was shown to have beneficial effects when directly injected into the myocardium 1-2 days prior to experimental MI^30^; however, this model does not have a clinical correlate and a single injection of PDGF-AB at the time of MI had no effect on outcomes^31^. These diverse findings highlight the lack of understanding of how different PDGF signaling pathways regulate the responses of the heart to injury, and the therapeutic potential of altering PDGFR signaling *in vivo* remains obscure.

We hypothesized that cardiac fibroblasts and their cCFU-F component are primary targets of PDGF ligands induced in the injury environment. Whereas inhibition of PDGFRα or β signaling has been shown to decrease collagen deposition after heart injury previously^27^, suggesting roles in fibrosis, their specific roles on fibroblast activation and differentiation have not been documented. Here, we set out to explore in detail the role of PDGFRα signaling on fibroblast heterogeneity, activation, proliferation and differentiation *in vitro* and *in vivo* using complementary approaches. Delivery of PDGF-AB via osmotic mini-pump for 5 days from the time of MI led to significant positive impacts on cardiac anatomical and functional repair including increased ejection fraction, and reduced end systolic and diastolic dimensions, and scar. Given the severity of infarcts imposed in this mouse study (mean EF ~20%), the degree of function improvement is noteworthy, and would be highly significant in the clinical setting. Our cellular and molecular analyses have demonstrated that the main impact of augmented PDGF-AB signaling in fibroblasts was to promote oxidative metabolism (increased mitochondrial mass and transcription factor expression) and increase ribosomal biosynthetic state, exit from G_0_ and increasing self-renewal capacity, without influencing fibroblast proliferation or fibrosis. We propose, therefore, that stimulation of PDGFRα with exogenous PDGF-AB ligand in this therapy setting significantly changes their metabolic and biosynthetic state, with associated changes in the dynamics and extent of intracellular and paracrine signaling contributing to heart repair.

We first used *Pdgfra^GFP/+^* knock-in mice as a novel tool for quantification of fibroblast activation during heart injury and repair by flow cytometry. The majority of resting fibroblasts were GFP^high^, whereas GFP^medium^ cells were identified as activated fibroblasts, aligning with the down-regulation of numerous stem cell markers including *Pdgfa* in activated fibroblast lineage cells, as shown by scRNAseq^16^. It is likely that at day 5 post-MI *Pdgfra*-GFP^medium^ cells contain a mixture of activated, pre-proliferative and proliferative fibroblasts (F-Act, F-CI, F-Cyc) and early myofibroblasts (MYO)^16^. Using PDGFR inhibitors and a specific PDGFRα blocking antibody, we found that PDGFRα signaling was essential for fibroblasts activation and/or differentiation *in vivo* using the *Pdgfra*-GFP readout, consistent with previous studies^27^. On the other hand, PDGF-AB treatment promoted fibroblast activation, leading to a ~50% increase in GFP^medium^ cells at the expense of GFP^high^ cells. However, PDGF-AB did not lead to increased BrdU incorporation in *Pdgfra*-GFP^medium^ cells.

We are aware that the increase in S^+^GFP^medium^ cells after PDGF-AB treatment may be exaggerated if there was a negative feedback from PDGFRα activation to *Pdgfra* transcription. However, this seems unlikely as we found that PDGF-AB treatment of cCFU-F cultures led to increased *Pdgfra* expression (data not shown). Furthermore, PyroninY/7AAD staining has provided a complementary and quantitative view of fibroblast activation/biosynthetic state, in this case reflecting cellular ribosomal (r)RNA content. Using this approach, we found that in uninjured hearts fibroblasts exist in a continuum of states with around half expressing extremely low levels of rRNA that we and others have interpreted as G_0_/quiescence^39^. Like other stem cell populations, cCFU-F, which are enriched in the SCA1^high^ subset of resting fibroblasts, were exclusively in quiescence, although after MI a significant proportion of them (~30%) became activated. As expected, GFP^medium^ cells induced during MI were virtually all activated. Importantly, even in the absence of injury, Pyronin Y staining showed that PDGF-AB was able to stimulate a substantial exit of fibroblasts from quiescence – this occurred without increasing BrdU incorporation or activation/differentiation using the *Pdgfra*-GFP readout. It was also accompanied by an increase in large colony formation and long-term self-renewal, demonstrating the functional impact of the effect. These data demonstrate a biosynthetic priming effect of PDGF-AB on fibroblasts.

We recently profiled cardiac fibroblasts by scRNAseq, revealing substantial fibroblast heterogeneity and non-linear cell dynamics^16^. We defined two major resting fibroblast populations expressing high and low levels of *Sca1*, and a novel activated fibroblast lineage (F-WntX) present in uninjured hearts. We also formally defined the major activated fibroblast population (F-Act), as well as pre-proliferative (F-CI) and proliferative fibroblast (F-Cyc) states, and at least 3 myofibroblast states, which appear after MI.

A key question is whether acute exogenous PDGF-AB delivery stimulates myofibrogenesis, although this seems unlikely given the pro-reparative effects elicited. We used scRNAseq to address this issue. We performed scRNAseq on TIP cells at day 7 post-MI to align with our previous analyses^16^. However, this showed no increase in the relative numbers of resting or activated fibroblasts (F-SH, F-SL, F-Act, F-Cyc), or myofibroblasts (MYO), compared to PBS-treated controls. Nonetheless, MYO, as well as other resting and activated fibroblasts (F-SL, F-SH, F-Act), and M1 and M2 macrophages, showed an increase in ribosomal biosynthetic state, without enhancement of secretory or ECM signatures. This likely occurs in response to increased PI3K/ATK/mTOR and cMYC signaling which are activated downstream of PDGFRα (Fig. 6C,D) and known to regulate ribosomal RNA biogenesis among other anabolic processes^45^. The ribosomal biosynthetic network signature has similarities to that induced in pre-proliferative fibroblasts (F-CI) that we defined previously at MI-day 3 using scRNAseq^16^. F-CI cells appear to exist in a cell trajectory continuum between activated (F-Act) and proliferative (F-Cyc) fibroblasts, and show strong upregulation of genes for ribosome biosynthesis and translation yet lack proliferative and myofibroblast ECM signatures^53^. For example, 57% of genes induced by PDGF-AB in MYO overlap with genes upregulated in F-CI compared to F-Act. Thus, multiple lines of evidence suggest that acute PDGF-AB treatment leads principally to an increase in ribosomal biosynthetic state in different fibroblast populations, as well as in macrophages, however without triggering fibrosis. The priming effect was minimal in ECs. It is possible that the accelerated resolution of inflammation observed in PDGF-AB-treated MI hearts (increased M2/M1 macrophage ratio) is related to the increased macrophage biosynthetic state.

Manipulation of PDGFRα signaling *in vitro* supported the above model. In freshly plated S^+^P^+^ cells, PDGF-AB treatment led to an increased probability of cell cycle reentry from the quiescent state over the first few cell cycles, and subsequently increased oxidative metabolism, colony formation and self-renewal, without effects on the rate of proliferation (cell cycle length) or commitment to differentiation. PDGFR inhibition had the opposite effect. Overall, our findings extend previous work showing that PDGF can act as a *competence* factor during re-entry of serum-starved immortalized fibroblasts to the cell cycle^54^. The primed state of PDGF-AB-induced cardiac fibroblasts may have similarities to G(Alert), a reversible, activated state induced in skeletal muscle stem cells by remote injury^40^. In common with G(Alert), PDGF-AB-treated S^+^P^+^ fibroblasts showed an increased rate of cell cycle re-entry *in vitro*. However, concomitant changes in cell size, mitochondrial content and activation of the AKT/mTOR pathway (other features of G(Alert)) were not observed in S^+^P^+^ cells after hind limb ischemic injury (data not shown), albeit that effects on minor populations (eg. cCFU-F) may have been masked.

PDGFRα blockade in both freshly plated and P4 S^+^P^+^ cells inhibited cell cycle reentry and induced a reversible low rRNA state akin to quiescence. This was accompanied by decreased colony formation and rate of self-renewal. Exit from this state required AKT and cMYC, which are known to be involved in G_0_/G_1_ checkpoint control and metabolic state, notably ribosomal biosynthesis. Inhibition led to altered expression of cell cycle regulators and to profound changes in the transcriptome. Down-regulation of expression of epigenetic pathways after PDGFRα inhibition suggests that the induced quiescence is accompanied by a reorganisation of chromatin, as seen in endogenous stem cell populations^55^. In addition to down-regulated genes, hundreds of genes were up-regulated in PDGFRα-inhibited cells, consistent with findings that maintenance of quiescence involves active programs^46^. The quiescence induced in cCFU-F cultures *in vitro* by PDGFR inhibition may resemble the quiescent state observed in ~50% of GFP^+^ fibroblasts in healthy hearts. Importantly, when we inhibited PDGFR signaling in uninjured hearts, even though the rRNA spectrum did not change, there was a decrease in colony formation and the rate of self-renewal *ex vivo* was strongly and permanently inhibited. Thus, fine control of PDGFR signaling is necessary for maintaining fibroblast metabolic and niche characteristics, as well as activation/differentiation of resting fibroblasts after injury. We have not yet analysed the precise impact of inhibiting endogenous PDGFR signaling on fibroblast activation versus differentiation during MI at single cell resolution, although this will be important to study using scRNAseq and functional assays.

Overall, our studies suggest that a dynamic continuum of cell metabolic and biosynthetic states underpins stromal fibroblast behavior, proliferation and lineage differentiation. This heterogeneity is reflected in the spectrum of colony forming ability, self-renewal, mitochondrial content and ribosomal biosynthetic states. Different matrix and vascular niche environments likely set up the continuum, which could be influenced by hypoxic, signaling or biomechanical gradients. These states are plastic, as seen after induction of MI, and PDGFRα stimulation or inhibition. Our studies reinforce the link between metabolism, and stem and progenitor cell states, and support the notion that a proportion of cardiac fibroblasts, those at the lower end of the ribosome biosynthetic spectrum, have stem cell-like properties^11,12^.

The pro-reparative effects of PDGF-AB on MI hearts likely involve cellular targets other than fibroblasts. Our data suggest that this could include ECs and macrophage precursors, and direct effects on CMs cannot be excluded^19^. PDGFRα expression has been reported on a minority of ECs^32,56^, although we did not see *Pdgfra*-GFP expression in ECs (or CMs) in healthy and injured hearts on sections (Fig. S3A-C and data not shown); however, flow cytometry has shown a minor sub-population of ECs with *Pdgfra*-GFP^low^ expression (data not shown). We found that PDGF-AB increased BrdU incorporation in EC in sham as well as MI hearts, and increased vascular density in remote as well as border and infarct regions after MI, suggesting a direct impact on ECs. While the identity and progenitor status of *Pdgfra*-GFP^low^ cells will be interesting to study, it is noteworthy that the injury response also includes pro-angiogenic paracrine effects from CMs via PDGFRβ^19^, as well as from myofibroblasts^16^. Additional work will be needed to fully understand the *in vivo* cellular targets of different PDGF ligands, and the complex inter-cellular communications participating in the PDGF-AB model of augmented cardiac repair.

There remains a strong clinical and economic case for the development of new regenerative therapies including modulators of fibrosis, and interestingly several existing cardiac standard-of-care drugs, such as statins, act, in part, as anti-fibrotics^4^. We have recently tested the impact of PDGF-AB therapy on cardiac repair after ischemia-reperfusion injury in the more clinically relevant pig model, finding significant improvements in anatomical and functional outcomes, as in the mouse MI model (JJHC, MX, NSA, MPF, RMG, RPH and colleagues, manuscript submitted).

In contrast to previous efforts to affect cardiac regeneration via cell therapy, implantation of bioengineered CM sheets, or cellular reprogramming by delivery of transcription factors^57,58^ or potent signaling intermediates^59,60^, the potential to stimulate the heart non-invasively by the simple parenteral administration of a soluble growth factor such as PDGF-AB, or matrix factors^61^, that activate endogenous pro-regenerative pathways, is clinically attractive. This is particularly the case in the postacute MI setting when invasive interventions are poorly tolerated, limiting the translatability of many of the therapeutic pro-regenerative approaches currently under evaluation.

## Author Contributions

N.S.A., M.X., V.C. and R.P.H conceived of the study; All authors designed and conducted experiments, analyzed data and contributed to the paper; R.P.H, R.M.G., M.F. and R.N. provided supervision; RPH provided financial support; N.S.A., M.X. and R.P.H. wrote the paper.

## Acknowledgements

We thank Elizabeth Kelly for technical assistance. We acknowledge support from the Australian Stem Cell Centre (including P082, S4M1), Australia India Strategic Research Fund (BF020084, BF050024), Australian Research Council Strategic Initiative in Stem Cell Science (SR110001002), Australian Academy of Science Visiting Scholar’s Program, Foundation Leducq Transatlantic Networks of Excellence in Cardiovascular Research (13CVD01, 15CVD03), New South Wales Cardiovascular Research Network (100711), James Kirby Foundation, and National Health and Medical Research Council of Australia (NHMRC; 354400, 573732, 1074386). RPH held an Australia Fellowship and Senior Principal Research Fellowship from the NHMRC (573705, 1118576). JJHC was supported by a Future Leader Fellowship (ID 100463) from the National Heart Foundation of Australia and a Sydney Medical School Foundation Fellowship.

## References

1 Kikuchi, K. et al. Primary contribution to zebrafish heart regeneration by gata4(+) cardiomyocytes. Nature 464, 601–605, doi:10.1038/nature08804 (2010).

2 Porrello, E. R. et al. Transient regenerative potential of the neonatal mouse heart. Science 331, 1078–1080, doi:10.1126/science.1200708 (2011).

3 Bergmann, O. et al. Dynamics of Cell Generation and Turnover in the Human Heart. Cell 161, 1566–1575, doi: 10.1016/j.cell.2015.05.026 (2015).

4 Gourdie, R. G., Dimmeler, S. & Kohl, P. Novel therapeutic strategies targeting fibroblasts and fibrosis in heart disease. Nature REviews Drug Discovery 15, 620–638, doi:10.1038/nrd.2016.89 (2016).

5 Shinde, A. V. & Frangogiannis, N. G. Fibroblasts in myocardial infarction: a role in inflammation and repair. Journal of Molecular and Cellular Cardiology 70, 74–82, doi:10.1016/j.yjmcc.2013.11.015 (2014).

6 Tallquist, M. D. & Molkentin, J. D. Redefining the identity of cardiac fibroblasts. Nature Reviews Cardiology 14, 484–491, doi:10.1038/nrcardio.2017.57 (2017).

7 Travers, J. G., Kamal, F. A., Robbins, J., Yutzey, K. E. & Blaxall, B. C. Cardiac Fibrosis: The Fibroblast Awakens. Circulation Research 118, 1021–1040, doi: 10.1161/CIRCRESAHA.115.306565 (2016).

8 Mescher, A. L. Macrophages and fibroblasts during inflammation and tissue repair in models of organ regeneration. Regeneration (Oxf) 4, 39–53, doi:10.1002/reg2.77 (2017).

9 Kaur, H. et al. Targeted Ablation of Periostin-Expressing Activated Fibroblasts Prevents Adverse Cardiac Remodeling in Mice. Circulation Research 118, 1906–1917, doi:10.1161/CIRCRESAHA.116.308643 (2016).

10 Oka, T. et al. Genetic manipulation of periostin expression reveals a role in cardiac hypertrophy and ventricular remodeling. Circulation Research 101, 313–321, doi:10.1161/CIRCRESAHA.107.149047 (2007).

11 Chong, J. J. et al. Adult cardiac-resident MSC-like stem cells with a proepicardial origin. Cell Stem Cell 9, 527–540, doi:10.1016/j.stem.2011.10.002 (2011).

12 Noseda, M. et al. PDGFRalpha demarcates the cardiogenic clonogenic Sca1 + stem/progenitor cell in adult murine myocardium. Nature Communications 6, 6930, doi:10.1038/ncomms7930 (2015).

13 Kfoury, Y. & Scadden, D. T. Mesenchymal cell contributions to the stem cell niche. Cell stem cell 16, 239–253, doi:10.1016/j.stem.2015.02.019 (2015).

14 Heredia, J. E. et al. Type 2 innate signals stimulate fibro/adipogenic progenitors to facilitate muscle regeneration. Cell 153, 376–388, doi:10.1016/j.cell.2013.02.053 (2013).

15 Cornwell, J. A., Nordon, R. E. & Harvey, R. P. Analysis of cardiac stem cell self-renewal dynamics in serum-free medium by single cell lineage tracking. Stem Cell Research 28, 115–124, doi:10.1016/j.scr.2018.02.004 (2018).

16 Farbehi, N. et al. Single-cell expression profiling reveals dynamic flux of cardiac stromal, vascular and immune cells in health and injury. eLife in press 26th March 2019 (2019).

17 Ivey, M. J. & Tallquist, M. D. Defining the Cardiac Fibroblast. Circulation Journal 80, 2269–2276, doi:10.1253/circj.CJ-16-1003 (2016).

18 Hellstrom, M., Kalen, M., Lindahl, P., Abramsson, A. & Betsholtz, C. Role of PDGF-B and PDGFR-beta in recruitment of vascular smooth muscle cells and pericytes during embryonic blood vessel formation in the mouse. Development 126, 3047–3055 (1999).

19 Chintalgattu, V. et al. Cardiomyocyte PDGFR-beta signaling is an essential component of the mouse cardiac response to load-induced stress. Journal of Clinical Investigation 120, 472–484, doi:10.1172/JCI39434 (2010).

20 Smith, C. L., Baek, S. T., Sung, C. Y. & Tallquist, M. D. Epicardial-derived cell epithelial-to-mesenchymal transition and fate specification require PDGF receptor signaling. Circ Res 108, e15–26, doi:10.1161/CIRCRESAHA.110.235531 (2011).

21 Andrae, J., Gallini, R. & Betsholtz, C. Role of platelet-derived growth factors in physiology and medicine. Genes and Development 22, 1276–1312, doi:10.1101/gad.1653708 (2008).

22 Olson, L. E. & Soriano, P. Increased PDGFRalpha activation disrupts connective tissue development and drives systemic fibrosis. Developmental Cell 16, 303–313, doi:10.1016/j.devcel.2008.12.003 (2009).

23 Savi, M. et al. Imatinib mesylate-induced cardiomyopathy involves resident cardiac progenitors. Pharmacology Research 127, 15–25, doi:10.1016/j.phrs.2017.09.020 (2018).

24 Gallini, R., Lindblom, P., Bondjers, C., Betsholtz, C. & Andrae, J. PDGF-A and PDGF-B induces cardiac fibrosis in transgenic mice. Experimenal Cell Research 349, 282–290, doi:10.1016/j.yexcr.2016.10.022 (2016).

25 Hsieh, P. C., MacGillivray, C., Gannon, J., Cruz, F. U. & Lee, R. T. Local controlled intramyocardial delivery of platelet-derived growth factor improves postinfarction ventricular function without pulmonary toxicity. Circulation 114, 637–644, doi: 10.1161/CIRCULATIONAHA.106.639831 (2006).

26 Edelberg, J. M. et al. Platelet-derived growth factor-AB limits the extent of myocardial infarction in a rat model: feasibility of restoring impaired angiogenic capacity in the aging heart. Circulation 105, 608–613 (2002).

27 Zymek, P. et al. The role of platelet-derived growth factor signaling in healing myocardial infarcts. Journal of the American College of Cardiology 48, 2315–2323, doi: 10.1016/j.jacc.2006.07.060 (2006).

28 Gallini, R., Huusko, J., Yla-Herttuala, S., Betsholtz, C. & Andrae, J. Isoform-Specific Modulation of Inflammation Induced by Adenoviral Mediated Delivery of Platelet-Derived Growth Factors in the Adult Mouse Heart. PLoS One 11, e0160930, doi:10.1371/journal.pone.0160930 (2016).

29 Cornwell, J. A. et al. Quantifying intrinsic and extrinsic control of single-cell fates in cancer and stem/progenitor cell pedigrees with competing risks analysis. Scientific Reports 6, 27100, doi:10.1038/srep27100 (2016).

30 Zheng, J. et al. Platelet-derived growth factor improves cardiac function in a rodent myocardial infarction model. Coron Artery Dis 15, 59–64 (2004).

31 Edelberg, J. M., Xaymardan, M., Lee, S. & Hong, M. K. Protection of Cardiac Myocardium. International Patent Number WO 2004/045531 (2004).

32 Edelberg, J. M. et al. PDGF mediates cardiac microvascular communication. J Clin Invest 102, 837–843, doi:10.1172/JCI3058 (1998).

33 Furtado, M. B. et al. Cardiogenic genes expressed in cardiac fibroblasts contribute to heart development and repair. Circulation Research 114, 1422–1434, doi:10.1161/CIRCRESAHA.114.302530 (2014).

34 Fu, X. et al. Specialized fibroblast differentiated states underlie scar formation in the infarcted mouse heart. Journal of Clinical Investigation 128, 2127–2143, doi:10.1172/JCI98215 (2018).

35 Ivey, M. J. et al. Resident fibroblast expansion during cardiac growth and remodeling. Journal of Molecular and Cellular Cardiology 114, 161–174, doi:10.1016/j.yjmcc.2017.11.012 (2018).

36 Moore-Morris, T. et al. Resident fibroblast lineages mediate pressure overload-induced cardiac fibrosis. Journal of Clinical Investigation 124, 2921–2934, doi:10.1172/JCI74783 (2014).

37 Moore-Morris, T. et al. Infarct Fibroblasts Do Not Derive From Bone Marrow Lineages. Circ Res 122, 583–590, doi:10.1161/CIRCRESAHA.117.311490 (2018).

38 Kanisicak, O. et al. Genetic lineage tracing defines myofibroblast origin and function in the injured heart. Nature Communications 7, 12260, doi: 10.1038/ncomms12260 (2016).

39 Kozma, S. C. & Thomas, G. Regulation of cell size in growth, development and human disease: PI3K, PKB and S6K. Bioessays 24, 65–71, doi: 10.1002/bies.10031 (2002).

40 Rodgers, J. T. et al. mTORC1 controls the adaptive transition of quiescent stem cells from G0 to G(Alert). Nature 510, 393–396, doi:10.1038/nature13255 (2014).

41 Demoulin, J. B. & Essaghir, A. PDGF receptor signaling networks in normal and cancer cells. Cytokine and Growth Factor Reviews 25, 273–283, doi: 10.1016/j.cytogfr.2014.03.003 (2014).

42 Tothova, Z. et al. FoxOs are critical mediators of hematopoietic stem cell resistance to physiologic oxidative stress. Cell 128, 325–339, doi: 10.1016/j.cell.2007.01.003 (2007).

43 Chiariello, M., Marinissen, M. J. & Gutkind, J. S. Regulation of c-myc expression by PDGF through Rho GTPases. Nat Cell Biol 3, 580–586, doi:10.1038/35078555 (2001).

44 Wilson, A. et al. c-Myc controls the balance between hematopoietic stem cell self-renewal and differentiation. Genes Dev 18, 2747–2763, doi:10.1101/gad.313104 (2004).

45 Davis, W. J., Lehmann, P. Z. & Li, W. Nuclear PI3K signaling in cell growth and tumorigenesis. Front Cell Dev Biol 3, 24, doi: 10.3389/fcell.2015.00024 (2015).

46 Chen, B. R. et al. Quiescent fibroblasts are more active in mounting robust inflammatory responses than proliferative fibroblasts. PLoS One 7, e49232, doi:10.1371/journal.pone.0049232 (2012).

47 Butler, A., Hoffman, P., Smibert, P., Papalexi, E. & Satija, R. Integrating single-cell transcriptomic data across different conditions, technologies, and species. Nat Biotechnol 36, 411–420, doi:10.1038/nbt.4096 (2018).

48 Finak, G. et al. MAST: a flexible statistical framework for assessing transcriptional changes and characterizing heterogeneity in single-cell RNA sequencing data. Genome Biol 16, 278, doi: 10.1186/s13059-015-0844-5 (2015).

49 Mi, H. et al. PANTHER version 11: expanded annotation data from Gene Ontology and Reactome pathways, and data analysis tool enhancements. Nucleic Acids Res 45, D183–D189, doi:10.1093/nar/gkw1138 (2017).

50 Szklarczyk, D. et al. The STRING database in 2017: quality-controlled protein-protein association networks, made broadly accessible. Nucleic Acids Res 45, D362–D368, doi:10.1093/nar/gkw937 (2017).

51 Vivien, C. J., Hudson, J. E. & Porrello, E. R. Evolution, comparative biology and ontogeny of vertebrate heart regeneration. NPJ Regenerative Medicine 1, 16012, doi:10.1038/npjregenmed.2016.12 (2016).

52 Hsieh, P. C., Davis, M. E., Gannon, J., MacGillivray, C. & Lee, R. T. Controlled delivery of PDGF-BB for myocardial protection using injectable self-assembling peptide nanofibers. J Clin Invest 116, 237–248, doi:10.1172/JCI25878 (2006).

53 Fonoudi, H. et al. A Universal and Robust Integrated Platform for the Scalable Production of Human Cardiomyocytes From Pluripotent Stem Cells. Stem Cells Transl Med 4, 1482–1494, doi:10.5966/sctm.2014-0275 (2015).

54 Jones, S. M. & Kazlauskas, A. Growth-factor-dependent mitogenesis requires two distinct phases of signalling. Nature Cell Biology 3, 165–172, doi: 10.1038/35055073 (2001).

55 Cheung, T. H. & Rando, T. A. Molecular regulation of stem cell quiescence. Nature Reviews Molecular and Cellular Biology 14, 329–340, doi:10.1038/nrm3591 (2013).

56 Chong, J. J. et al. Progenitor cells identified by PDGFR-alpha expression in the developing and diseased human heart. Stem Cells Dev 22, 1932–1943, doi:10.1089/scd.2012.0542 (2013).

57 Xiao, Y., Leach, J., Wang, J. & Martin, J. F. Hippo/Yap Signaling in Cardiac Development and Regeneration. Curr Treat Options Cardiovasc Med 18, 38, doi:10.1007/s11936-016-0461-y (2016).

58 Srivastava, D. & DeWitt, N. In Vivo Cellular Reprogramming: The Next Generation. Cell 166, 1386–1396, doi:10.1016/j.cell.2016.08.055 (2016).

59 D’Uva, G. et al. ERBB2 triggers mammalian heart regeneration by promoting cardiomyocyte dedifferentiation and proliferation. Nat Cell Biol 17, 627–638, doi: 10.1038/ncb3149 (2015).

60 Mohamed, T. M. A. et al. Regulation of Cell Cycle to Stimulate Adult Cardiomyocyte Proliferation and Cardiac Regeneration. Cell 173, 104–116 e112, doi: 10.1016/j.cell.2018.02.014 (2018).

61 Bassat, E. et al. The extracellular matrix protein agrin promotes heart regeneration in mice. Nature 547, 179–184, doi:10.1038/nature22978 (2017).

