## Supplementary material TEXT+Figures for "PDGFRα signaling in cardiac fibroblasts modulates quiescence, metabolism and self-renewal, and promotes anatomical and functional repair"

**Supplementary Files List**

1. **Supplementary figures and legend**
2. **Detailed methods and materials**
3. **Key resources table (kits and reagents)**
4. **References of the supplementary materials**

**Supplementary figures and legends**

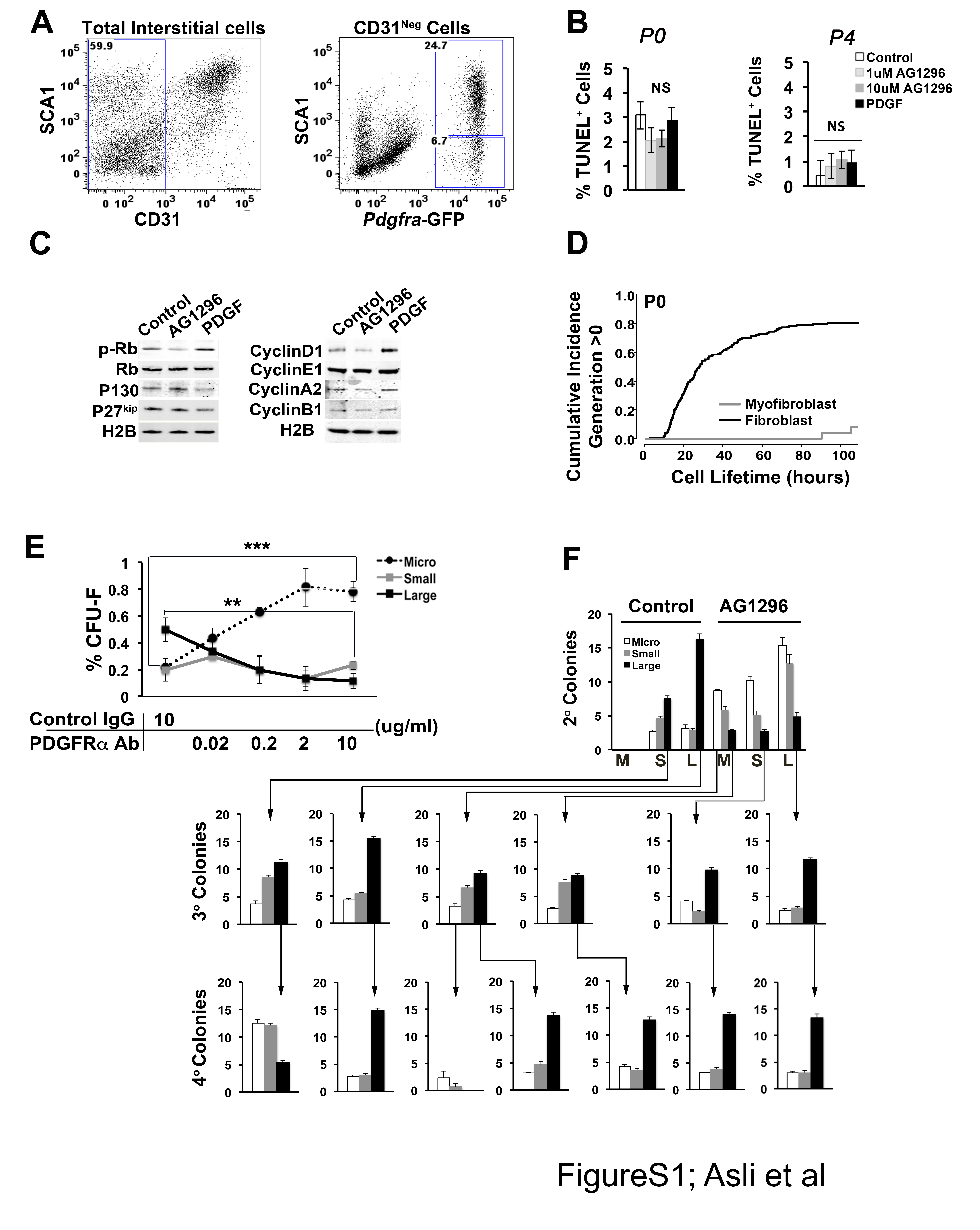

**Figure S1: Cell cycle, cell death and colony-forming assays in AG1296 and PDGF-AB-treated cultures**

(A) Flow cytometry plots demonstrating the distribution of SCA1 and CD31 cell surface expression within total cardiac interstitial cells and *Pdgfra*-GFP^+^ fibroblasts. Cardiac fibroblasts are composed of SCA1^high^ and SCA1^low^ sub-compartments. (B) Percentage of TUNEL-positive cells in S^+^P^+^ cultures treated throughout P0 and P4, respectively, with vehicle (control), AG1296 or PDGF-AB (100ng/ml). (C) Western blot analysis of cell cycle-associated proteins after treatment of S^+^P^+^ cultures across P4 with AG1296 or PDGF-AB. (D) Accumulated proportion of plated S^+^P^+^ cells showing fibroblastic or myofibroblastic morphologies undergoing cell division (generations >0). (E) cCFU-F analysis of S^+^P^+^ cells in the presence of control IgG antibody or blocking anti-PDGFRα monoclonal antibody, scored at the end of P0. (F) Selected data from a serial clonal cCFU-F analysis of pooled large (L), small (S) and micro-colonies (M) initially arising from freshly plated S^+^P^+^ cells cultured in vehicle (control) or AG1296 until the end of P0. Graphs show secondary (2^o^), tertiary (3^o^) and quaternary (4^o^) colony assays after cloned primary colonies were passaged until the end of P1, P2 and P3 in the absence of inhibitor.

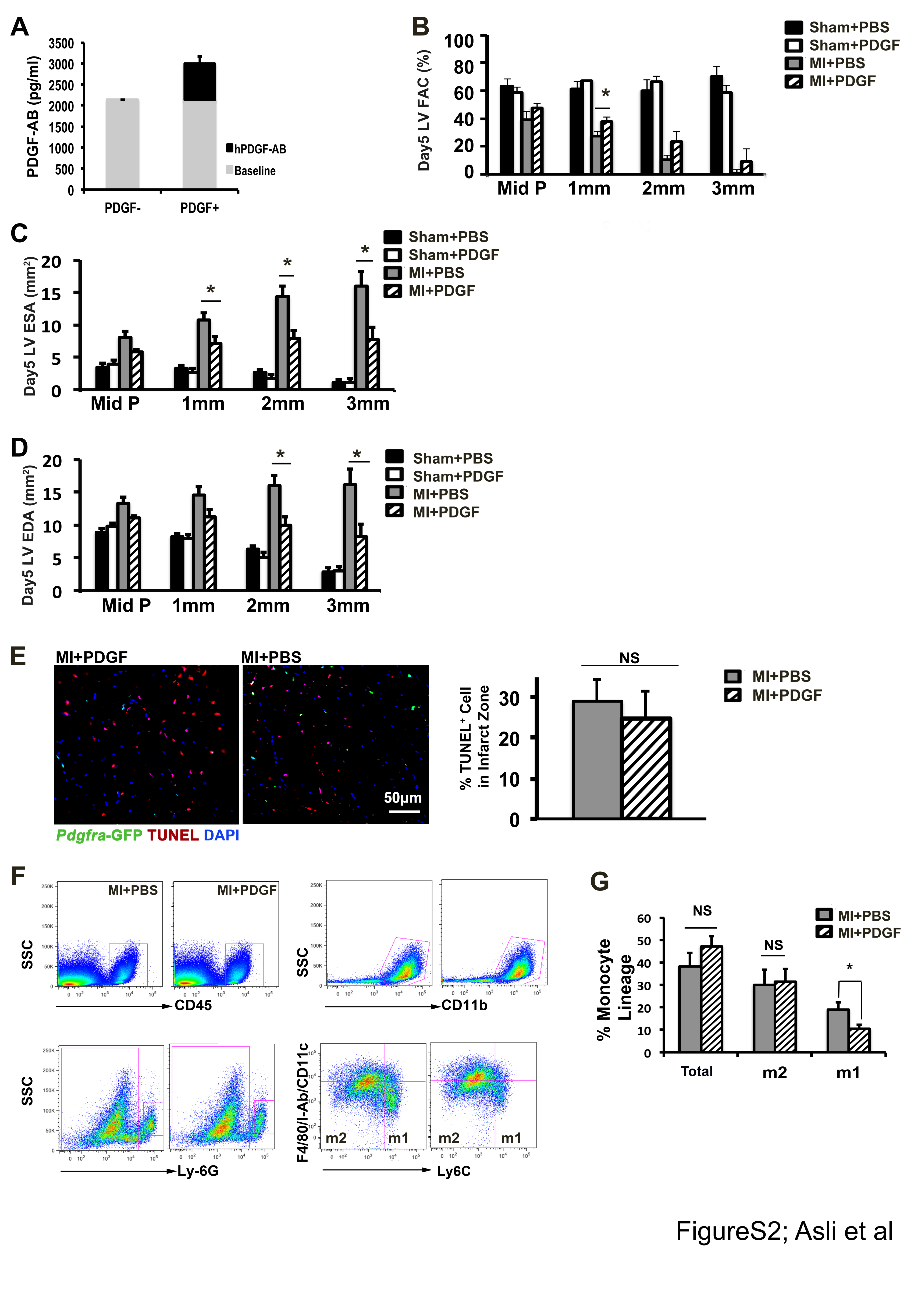

**Figure S2: Quantification of serum PDGF-AB, additional cardiac functional analyses, monocyte quantification strategy and TUNEL studies.**

(A) Levels of mouse (baseline) and human (hPDGF-AB) detected in mice untreated or treated with exogenous human recombinant PDGF-AB, analysed by ELISA. (B-D) Quantification of functional parameters in MI+PDGF-AB hearts at day 5 post-surgery, compared to sham+PBS, sham+PDGF-AB and MI+PBS controls – fractional area change (FAC), left ventricular end systolic area (LV ESA), and left ventricular end diastolic area (LV EDA), respectively, at different echocardiographic planes (see Fig. 2E). (E) Quantification of the TUNEL positive cells in the initial infarct region of MI+PDGF and MI+PBS hearts 6 hours post-infarct. (F) Flow cytometric analysis strategy using side scatter (SSC) and indicated antibodies applied to total cardiac interstitial cells from control MI+PBS and MI+PDGF-AB hearts used to define m1 and m2 monocyte fractions (see Fig. 2L). (G) Quantification of total monocyte lineage population as well as m1 and m2 monocytes from control MI+PBS and MI+PDGF-AB hearts. All comparisons: * p<0.05; *** p<0.001; NS: non-significant.

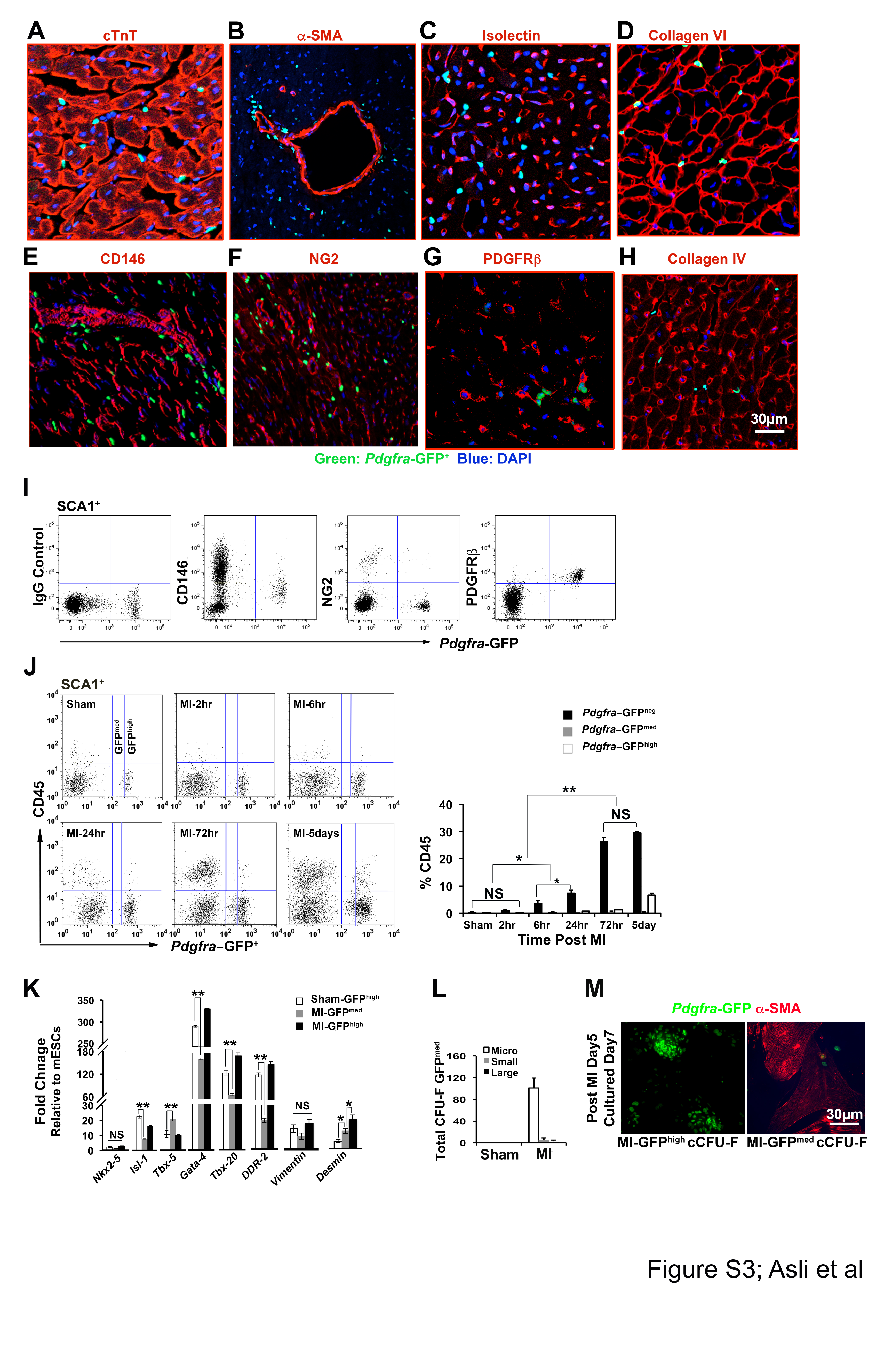

**Figure S3: Colocalisation of the *Pdfra*GFP/+ cells and vascular/pericyte markers in the myocardium. Flow cytometric and gene expression analysis of various markers in SCA1^+^ *Pdgfra*-GFP^+^ fibroblasts in control and treated hearts.**

(A-H) Localization of GFP^+^ stromal cells in sections of hearts from *Pdgfra*^GFP/+^ knock-in mice relative to indicated markers detected by immunofluorescence: cTnT: cardiac troponin T; α-SMA: α-smooth muscle actin. (I) Flow cytometric analysis of indicated pericyte markers on cardiac SCA1^+^ *Pdgfra*-GFP^+^ cells. (J) Flow cytometric analysis of CD45 and GFP expression in freshly isolated SCA1^+^ cardiac interstitial cells from *Pdgfra*^GFP/+^ mice in a time-course post myocardial infarction (MI), compared to cells from sham-operated mice. Graph indicates percent CD45^+^ cells in GFP^negative^, GFP^medium^ and GFP^high^ cardiac interstitial populations. (K) Quantitative RT-PCR analysis of indicated cardiac transcription factors and fibroblast markers in flow-purified GFP^medium^ and GFP^high^ cells from *Pdgfra*^GFP/+^ mice subject to sham operation or MI. Fold change is relative to expression in undifferentiated mouse embryonic stem cells (mESCs). Most cardiac fibroblasts express cardiac transcription factor genes *Isl1,* *Gata4* and *Tbx20*, reflecting their epicardial origins and cardiac identity (Furtado, Costa et al. 2014, Noseda, Harada et al. 2015). However, S^+^GFP^medium^ myofibroblasts showed reduced expression of these genes and of fibroblast markers vimentin and DDR-2. (L) cCFU-F analysis of GFP^medium^ population from day 5 post-sham and MI mice as defined in Fig. 3A. S^+^GFP^medium^ cells formed only micro-colonies, which are typically composed of differentiated myofibroblast-like cells (Chong, Chandrakanthan et al. 2011, Cornwell, Nordon et al. 2018). All comparisons: * p<0.05; ** p<0.01. NS: non-significant. (M) Micrographs showing GFP fluorescence and α-SMOOTH MUSCLE ACTIN (α-SMA) immunofluorescence staining in 7 day cCFU-F cultures of flow-purified GFP^medium^ and GFP^high^ cells from *Pdgfra*^GFP/+^ mice subject to MI (cells harvested at day 5 post-MI). S^+^GFP^medium^ cells resembled myofibroblasts (large GFP^dim^SMA^high^ cells with evidence of actin stress fibres), whereas GFP^high^ cells were spindle-shaped, remained GFP^bright^ and were predominantly SMA^-^.

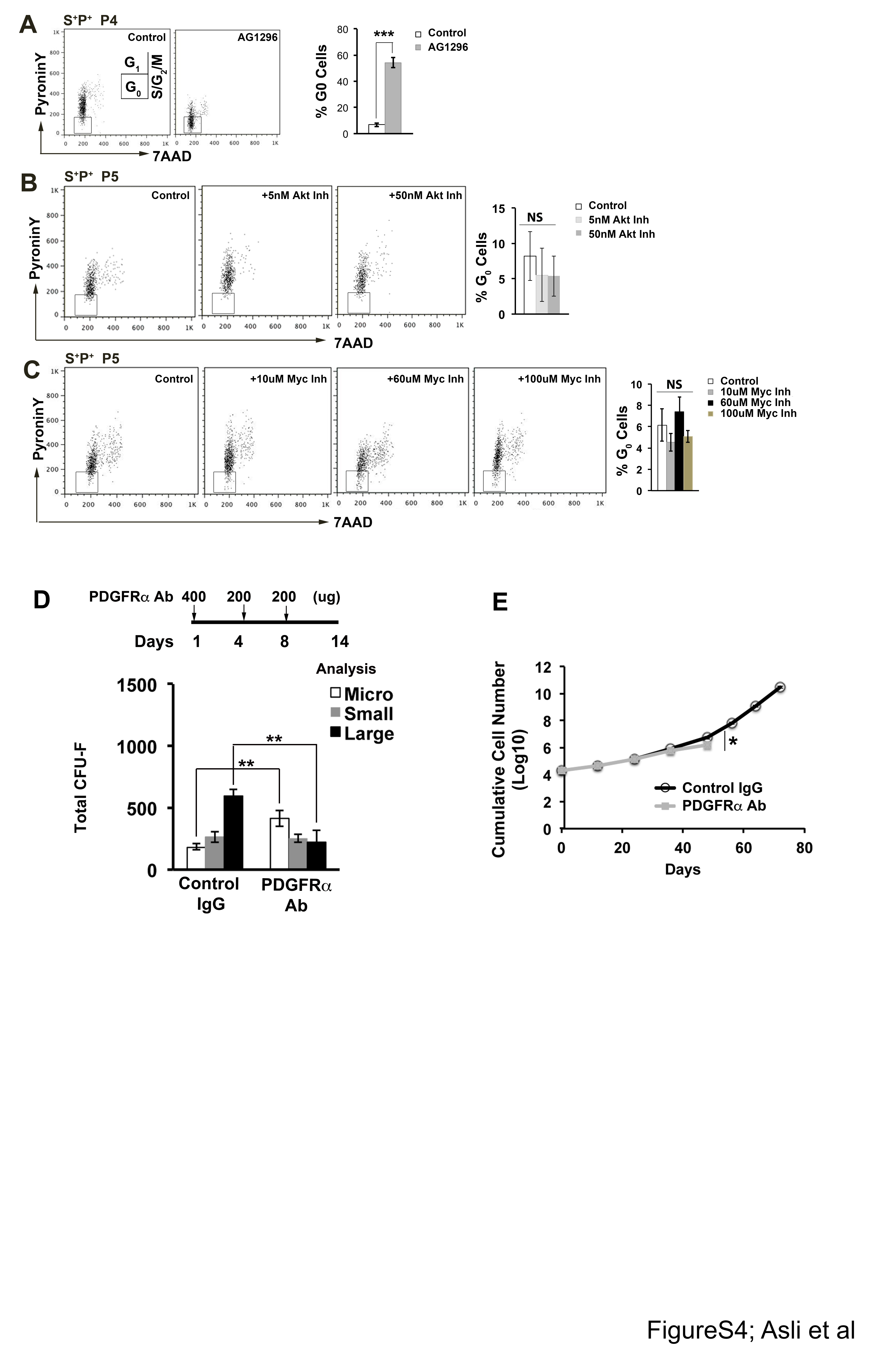

**Figure S4: Inhibition of PDGFRα *in vivo* using blocking anti-PDGFRα antibody, Signaling requirements of cCFU-F cultures**

(A) Cell cycle analysis of S^+^P^+^ cultures treated with vehicle (control) or PDGFRα inhibitor AG1296 over P4 (see Fig. 6E,F). Graph shows percent cells in G_0_ (boxed). (B,C) Cell cycle analysis and graph of percentage of cells in G_0_ of S^+^P^+^ cultures treated across P5 with 5nM and 50nM AKT inhibitor (LY294002) or 10μM, 60μM, and 100μM MYC inhibitor (10058-F4), respectively, compared to vehicle (control). (D) Treatment regime applicable to both control IgG or blocking PDGFRα (APA5) antibody, and graph showing results of colony-forming (cCFU-F) assay performed at day 14 post-treatment. (E) Long-term growth of S^+^P^+^ cultures from control IgG and PDGFRα monoclonal antbody-treated animals initiated at day 14 post-treatment. All comparisons: * p<0.05; ** p<0.01; *** p<0.001.

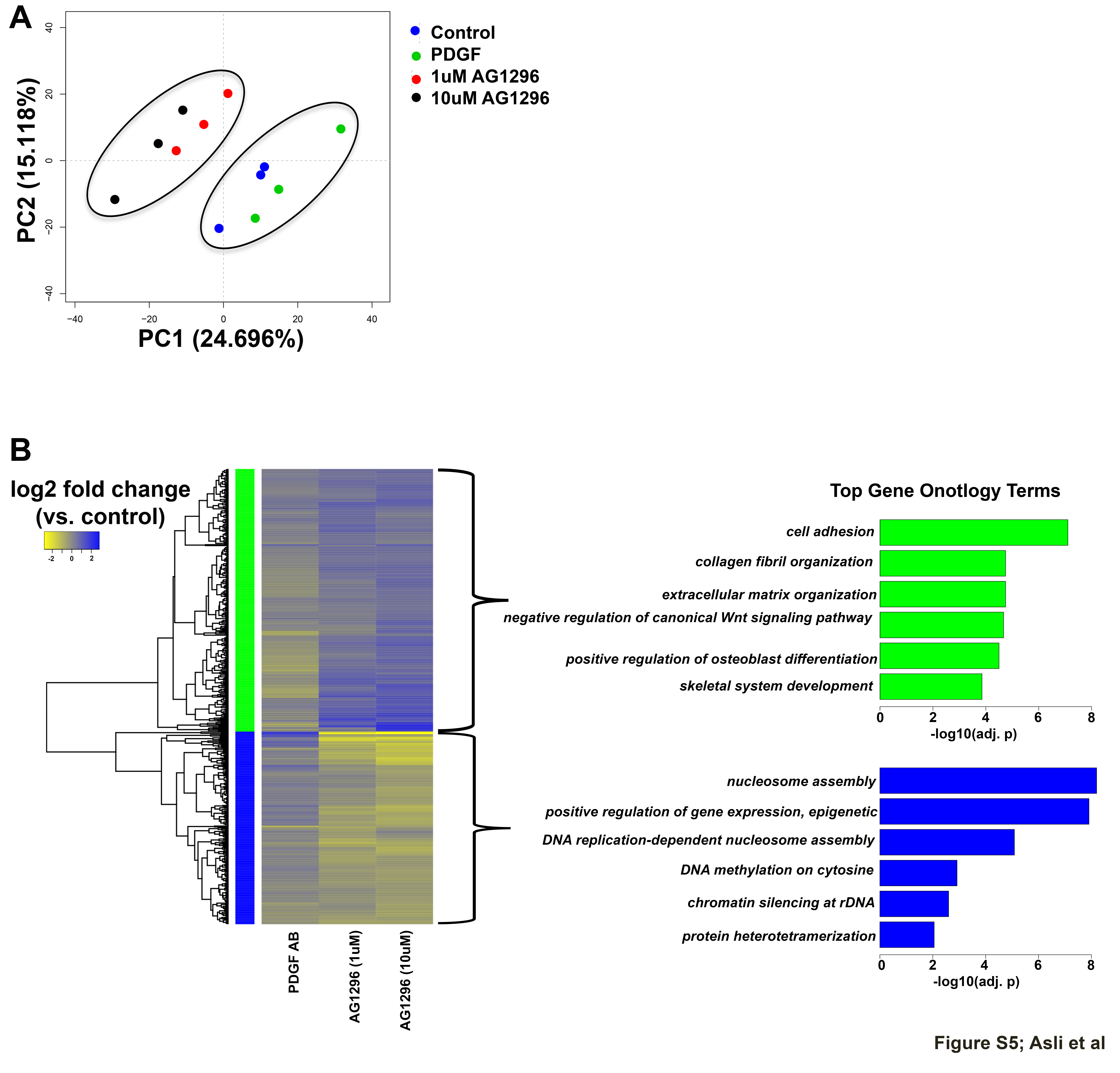

**Figure S5: Transcriptome analysis of PDGF and AG1296-treated cultures**

(A) Principal component (PC) analysis (PC1 vs PC2) of microarray transcriptome data from S^+^P^+^ cells cultured to P0 with and without PDGF-AB (100ng/ml) and AG1296 (1 and 10uM). (B) Heat map of differentially expressed genes (log2 ratio) between AG1296 (1uM), AG1296 (10uM), or PDGF-AB-treated cells versus untreated controls (adjusted p value <0.05); and top gene ontology terms for biological processes (benjamini adjusted p values) of the two distinguishing clusters (up-regulated and down-regulated with inhibitor compared to PDGF-AB-treated).

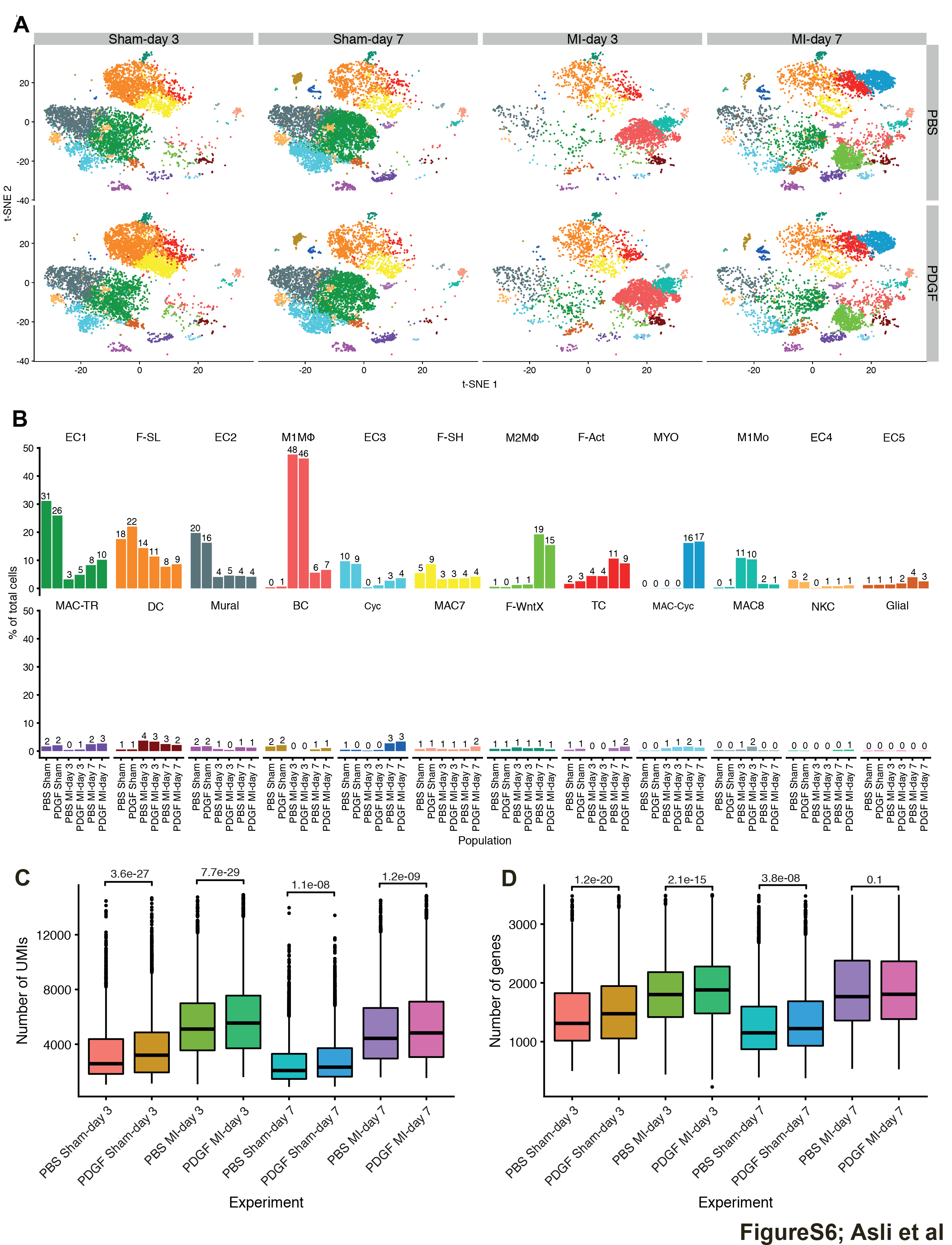

**Figure S6: Comparisons of experimental conditions across scRNA-seq from PBS and PDGF treated and injured and uninjured hearts**

(A) t-SNE visualisation of scRNA-seq cell populations for all sham and MI time-points comparing PBS (top row) and PDGF (bottom row) treatments. (B) Differential Proportion Analysis (DPA) showing percentage of total cells in each cell population according to experimental condition, where sham time-points have been merged to show an aggregate sham PBS and sham PDGF-AB. (C,D) Number of unique molecular identifiers (UMIs) (C) and genes (D) detected in TIP across all experimental conditions. Comparison values indicate P-values from Wilcoxon rank-sum tests, with number of cells per-experiment (n) ranging from 3050 (PBS MI-day 3) to 7358 (PBS sham-day 7).

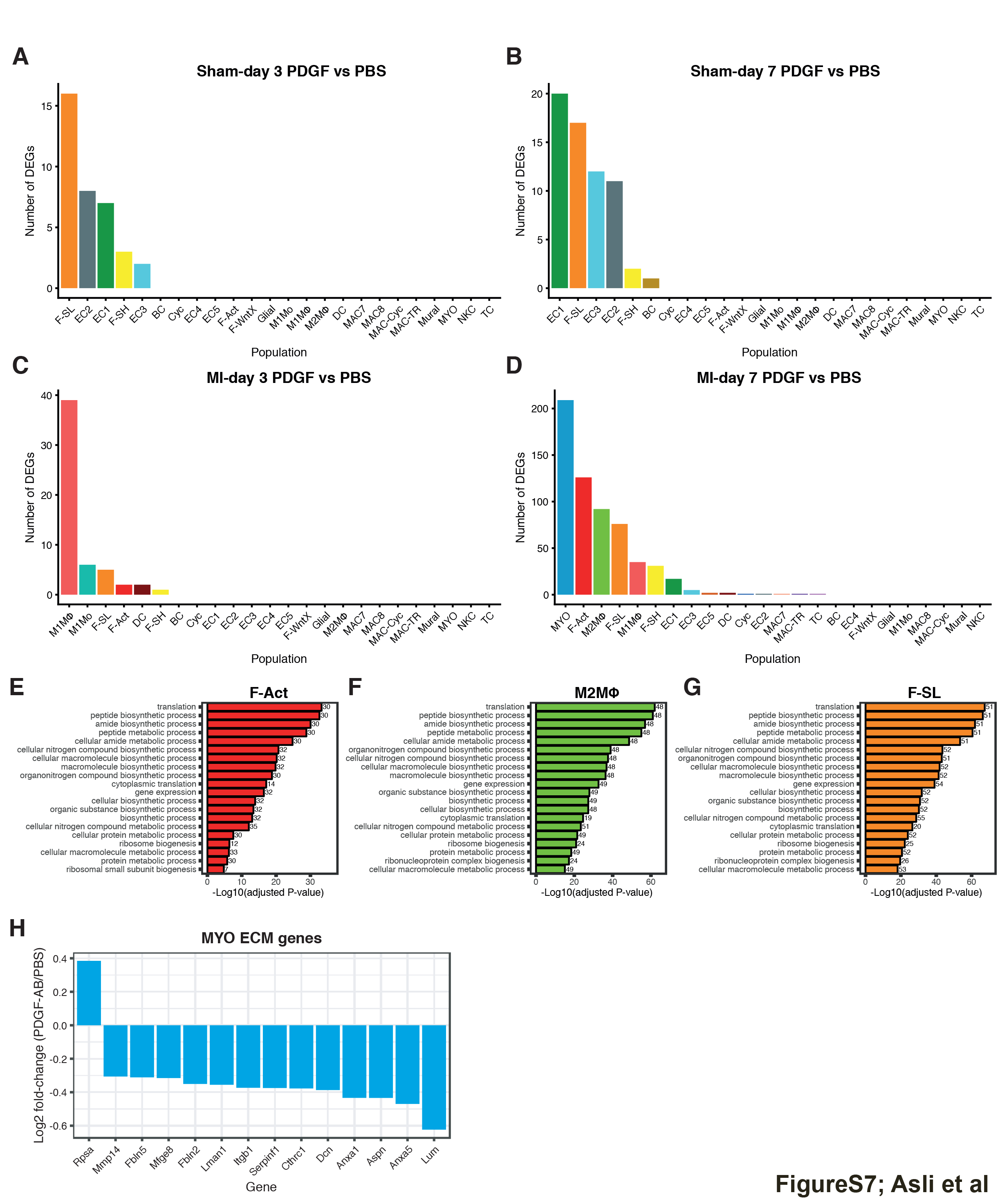

**Figure S7:** **Differential expression and gene ontology (GO) analysis of scRNA-seq cell populations comparing PDGF and PBS conditions**

(A-C) Counts of population-specific differentially expressed (DE) genes between PDGF-AB and PBS conditions at (A) sham-day 3, (B) sham-day 7, (C) MI-day 3 and (D) MI-day 7. (E-F) Top 20 over-represented GO biological process terms (PANTHER; P_adj_ < 0.05) for genes upregulated in PDGF-AB vs. PBS conditions at MI-day 7 within cell populations (E) F-Act, (F) M2 macrophages (MΦ) and (G) F-SL. (H) Average fold-change (log2) difference in differentially expressed MYO extracellular matrix (ECM) genes between PDGF-AB and PBS conditions at MI-day 7.

**Table 1. List of differentially expressed genes in MYO between PDGF-AB and PBS treated MI day7 hearts**

| **Gene** | **pct.1** | **pct.2** | **p_val_adj** | **Log2FoldChange** |
| --- | --- | --- | --- | --- |
| Rps12 | 1 | 0.982 | 9.53537E-75 | 0.65884374 |
| Klf4 | 0.69 | 0.55 | 6.42266E-11 | 0.623729711 |
| Ier3 | 0.71 | 0.527 | 1.21854E-10 | 0.53661721 |
| Rpl23 | 1 | 0.993 | 1.96589E-80 | 0.534466058 |
| Tpt1 | 1 | 0.997 | 4.83375E-80 | 0.533228339 |
| Rplp1 | 1 | 0.997 | 2.10675E-64 | 0.531423006 |
| Rps26 | 0.99 | 0.968 | 6.0721E-52 | 0.531316897 |
| Neat1 | 0.84 | 0.739 | 2.11539E-14 | 0.529670458 |
| Fosb | 0.72 | 0.556 | 1.6128E-10 | 0.526267589 |
| Rpl30 | 0.98 | 0.961 | 2.48082E-56 | 0.525892745 |
| Rps20 | 0.99 | 0.989 | 5.55498E-50 | 0.514757169 |
| Rps10 | 0.99 | 0.977 | 1.25471E-52 | 0.514604199 |
| Rpl39 | 0.99 | 0.989 | 3.3013E-50 | 0.510949214 |
| Rps15a | 0.99 | 0.996 | 2.3995E-63 | 0.510903513 |
| Atf3 | 0.34 | 0.199 | 8.72902E-08 | 0.504653602 |
| Ifi27l2a | 0.82 | 0.71 | 3.30066E-06 | 0.502446438 |
| Rpl28 | 1 | 0.988 | 8.42668E-61 | 0.500769669 |
| Rpl12 | 0.99 | 0.966 | 2.4856E-40 | 0.495344576 |
| Rps15 | 1 | 0.997 | 4.00837E-64 | 0.490449431 |
| Rps21 | 0.98 | 0.94 | 4.65864E-45 | 0.483391983 |
| Rps8 | 1 | 0.998 | 5.63271E-67 | 0.482176304 |
| Eef1a1 | 1 | 0.999 | 3.91608E-68 | 0.476179645 |
| Rpl35a | 1 | 0.994 | 1.74493E-59 | 0.47019433 |
| Rplp2 | 0.99 | 0.99 | 2.89923E-54 | 0.470028646 |
| Rps24 | 0.99 | 0.993 | 2.45244E-41 | 0.454600015 |
| Hspb1 | 0.75 | 0.664 | 1.14739E-07 | 0.447626332 |
| Fau | 1 | 0.994 | 3.91923E-58 | 0.446858069 |
| Rpl31 | 1 | 0.994 | 2.9879E-45 | 0.443506808 |
| Rpl5 | 0.98 | 0.964 | 1.03373E-35 | 0.435992535 |
| Rps12-ps3 | 0.74 | 0.544 | 2.00782E-18 | 0.429506407 |
| Gnb2l1 | 0.98 | 0.961 | 1.63054E-27 | 0.425130758 |
| Rps25 | 0.99 | 0.986 | 2.39291E-45 | 0.421714502 |
| Eef1b2 | 0.98 | 0.94 | 1.67318E-26 | 0.420116557 |
| Rpl29 | 0.99 | 0.983 | 6.9632E-35 | 0.41881437 |
| Rps2 | 1 | 0.999 | 2.62203E-35 | 0.417417842 |
| Rps4x | 1 | 0.994 | 3.64276E-37 | 0.408322721 |
| Zfp36 | 0.5 | 0.345 | 1.90102E-06 | 0.406370898 |
| Rps17 | 1 | 0.99 | 8.26439E-37 | 0.404883934 |
| Rpl21 | 1 | 0.995 | 3.58155E-41 | 0.40399087 |
| Rps13 | 0.99 | 0.993 | 5.97497E-43 | 0.403754548 |
| Rpl8 | 1 | 0.996 | 1.83566E-44 | 0.40360879 |
| Rpl10a | 1 | 0.992 | 9.839E-29 | 0.400638331 |
| Egr1 | 0.87 | 0.767 | 7.84292E-09 | 0.392845608 |
| Rpl19 | 1 | 0.993 | 5.51187E-38 | 0.388787386 |
| Rpsa | 1 | 0.994 | 3.88158E-25 | 0.384870722 |
| Pmepa1 | 0.94 | 0.89 | 9.64434E-14 | 0.383581179 |
| Rpl22 | 0.99 | 0.969 | 1.21909E-28 | 0.378219342 |
| Rpl38 | 0.99 | 0.971 | 9.775E-25 | 0.376927685 |
| Rpl36a | 1 | 0.986 | 4.33042E-31 | 0.376065026 |
| Rpl13a | 1 | 0.999 | 1.25034E-32 | 0.375649429 |
| Rpl27a | 1 | 0.997 | 1.24199E-35 | 0.375595264 |
| Rpl10 | 1 | 0.997 | 1.69267E-28 | 0.374941271 |
| Rpl7a | 0.99 | 0.983 | 8.69303E-30 | 0.368148178 |
| Rpl7 | 1 | 0.994 | 7.41357E-32 | 0.367246552 |
| Rpl18 | 1 | 0.992 | 7.88532E-29 | 0.357515987 |
| Csrp2 | 0.98 | 0.962 | 2.00898E-08 | 0.357425908 |
| Rpl27 | 0.96 | 0.902 | 5.81496E-19 | 0.355750581 |
| Rpl26 | 1 | 0.995 | 1.04752E-26 | 0.355053836 |
| Rpl23a | 1 | 0.992 | 7.23701E-28 | 0.354675659 |
| Rpl11 | 1 | 0.995 | 1.59076E-33 | 0.350523073 |
| Btg2 | 0.78 | 0.642 | 6.77548E-06 | 0.344564432 |
| Rpl17 | 1 | 0.995 | 3.10177E-22 | 0.341516356 |
| Rps7 | 0.99 | 0.994 | 5.72043E-20 | 0.340605518 |
| Igfbp4 | 0.9 | 0.852 | 6.74582E-07 | 0.339371585 |
| Rpl6 | 1 | 0.998 | 1.04668E-24 | 0.337323878 |
| Gm26532 | 0.46 | 0.299 | 6.83621E-06 | 0.334078128 |
| Rps27a | 1 | 0.997 | 9.71771E-24 | 0.333834037 |
| Rpl34 | 1 | 0.994 | 9.84442E-29 | 0.332556427 |
| Rps16 | 1 | 0.996 | 1.79402E-25 | 0.330850375 |
| Nfix | 0.91 | 0.841 | 8.4207E-18 | 0.328756619 |
| Zfos1 | 0.72 | 0.596 | 9.60532E-09 | 0.327868305 |
| Rps29 | 1 | 0.997 | 9.49898E-25 | 0.324830729 |
| Rpl32 | 1 | 0.995 | 7.13927E-19 | 0.320963298 |
| Rpl37 | 1 | 0.996 | 6.29028E-23 | 0.31649226 |
| Rps3a1 | 1 | 0.998 | 4.43385E-21 | 0.311299254 |
| Rps3 | 1 | 0.996 | 3.69935E-22 | 0.310952391 |
| Rpl9 | 0.99 | 0.996 | 4.79465E-22 | 0.300866099 |
| Hspe1 | 0.91 | 0.851 | 2.17848E-10 | 0.300857936 |
| Rpl37a | 1 | 0.997 | 2.82072E-19 | 0.291067782 |
| Eef2 | 1 | 0.988 | 3.23867E-13 | 0.290754825 |
| Ext1 | 0.61 | 0.453 | 4.47734E-07 | 0.289103427 |
| Rplp0 | 1 | 0.999 | 1.48827E-10 | 0.288502595 |
| Rps23 | 1 | 0.999 | 6.98616E-19 | 0.282824238 |
| Synpo | 0.66 | 0.521 | 1.65057E-06 | 0.280907998 |
| Rpl14 | 1 | 0.994 | 6.38654E-19 | 0.280158844 |
| Hsp90ab1 | 0.99 | 0.989 | 3.1066E-08 | 0.276033369 |
| mt-Nd2 | 0.97 | 0.91 | 6.31576E-18 | 0.272750551 |
| Atp5g2 | 0.92 | 0.852 | 4.21019E-06 | 0.264920161 |
| Rpl41 | 1 | 1 | 5.65426E-09 | 0.264656987 |
| Atp5e | 0.95 | 0.913 | 9.28223E-09 | 0.260402172 |
| Rpl23a-ps3 | 0.81 | 0.706 | 1.411E-06 | 0.254832222 |
| Erp44 | 0.54 | 0.62 | 1.26531E-06 | -0.250388069 |
| Atp5b | 0.85 | 0.909 | 2.53215E-14 | -0.252511634 |
| Lamp1 | 0.96 | 0.962 | 2.03036E-08 | -0.253526943 |
| Cnn3 | 0.86 | 0.891 | 1.90202E-08 | -0.254038594 |
| Myl12a | 0.99 | 0.998 | 6.30424E-22 | -0.255450649 |
| Morf4l1 | 0.96 | 0.966 | 7.93297E-11 | -0.256112757 |
| Scfd1 | 0.32 | 0.424 | 1.73472E-06 | -0.256357965 |
| Tspan3 | 0.81 | 0.86 | 6.85289E-08 | -0.256666542 |
| Nucb2 | 0.54 | 0.632 | 8.08659E-06 | -0.258016921 |
| Cct8 | 0.56 | 0.633 | 1.13429E-07 | -0.258023128 |
| Ldha | 0.88 | 0.915 | 1.4242E-10 | -0.259027608 |
| Rab18 | 0.7 | 0.765 | 2.6828E-07 | -0.265209129 |
| Tsn | 0.7 | 0.766 | 1.521E-07 | -0.267269545 |
| Vcp | 0.77 | 0.826 | 4.60898E-08 | -0.267296815 |
| Aga | 0.43 | 0.532 | 1.00126E-06 | -0.268643906 |
| Ssb | 0.57 | 0.669 | 2.50268E-08 | -0.269684265 |
| Eif2s1 | 0.39 | 0.49 | 2.4667E-08 | -0.272025832 |
| Ddost | 0.8 | 0.85 | 2.50263E-11 | -0.272084815 |
| Psmc3 | 0.59 | 0.675 | 2.56445E-09 | -0.274211952 |
| Cope | 0.87 | 0.906 | 5.40419E-13 | -0.276835813 |
| Atp6ap2 | 0.44 | 0.556 | 7.6948E-06 | -0.277503735 |
| Cct3 | 0.41 | 0.519 | 2.2602E-09 | -0.27838034 |
| Copz2 | 0.87 | 0.897 | 4.76374E-12 | -0.279190487 |
| Tuba1a | 0.89 | 0.928 | 6.87198E-09 | -0.28001724 |
| Mdh2 | 0.72 | 0.79 | 9.48879E-10 | -0.281036521 |
| Ywhaq | 0.77 | 0.813 | 5.52206E-09 | -0.282067196 |
| Tpr | 0.63 | 0.707 | 7.27358E-07 | -0.282196572 |
| Psmd11 | 0.53 | 0.639 | 1.83545E-09 | -0.283070042 |
| Oat | 0.67 | 0.763 | 5.13718E-11 | -0.284810133 |
| Dnajb11 | 0.62 | 0.705 | 5.63574E-08 | -0.286321399 |
| Srp54b | 0.64 | 0.734 | 7.06042E-10 | -0.288877112 |
| Pdgfrl | 0.86 | 0.909 | 9.9072E-08 | -0.290922003 |
| Arl1 | 0.85 | 0.89 | 3.2517E-12 | -0.291282642 |
| Smarca5 | 0.47 | 0.57 | 1.29983E-06 | -0.291623691 |
| Shisa5 | 0.79 | 0.862 | 5.27241E-10 | -0.291703998 |
| Hexa | 0.72 | 0.823 | 4.6799E-09 | -0.291867364 |
| Lrpap1 | 0.66 | 0.745 | 7.9989E-08 | -0.292879133 |
| Sec11a | 0.79 | 0.853 | 1.79671E-11 | -0.29338635 |
| Rbbp7 | 0.44 | 0.54 | 1.48905E-07 | -0.293610629 |
| Uba5 | 0.39 | 0.51 | 4.48675E-09 | -0.297371564 |
| Psmb2 | 0.78 | 0.84 | 1.48892E-13 | -0.297699025 |
| Hdac1 | 0.28 | 0.411 | 5.07802E-12 | -0.301289676 |
| Plpp1 | 0.63 | 0.739 | 1.60685E-07 | -0.301503391 |
| Rap1a | 0.59 | 0.687 | 1.06446E-08 | -0.301612025 |
| Mxra8 | 0.91 | 0.937 | 6.42885E-10 | -0.301798723 |
| Tecr | 0.61 | 0.726 | 5.33538E-11 | -0.30279151 |
| Copb2 | 0.56 | 0.665 | 5.26718E-11 | -0.304081254 |
| Gpx8 | 0.9 | 0.915 | 1.75289E-14 | -0.306006435 |
| Mmp14 | 0.81 | 0.894 | 7.55838E-10 | -0.30626796 |
| Kdelr3 | 0.76 | 0.806 | 6.64915E-10 | -0.307369619 |
| Dnajc3 | 0.68 | 0.754 | 6.3154E-09 | -0.307848497 |
| Ubc | 0.95 | 0.969 | 8.32126E-18 | -0.308884348 |
| Prkar1a | 0.72 | 0.806 | 2.65107E-12 | -0.309166891 |
| Fuca1 | 0.7 | 0.79 | 7.93562E-10 | -0.309620597 |
| Fbln5 | 0.88 | 0.917 | 4.03066E-06 | -0.311188744 |
| Hnrnpf | 0.73 | 0.809 | 3.48491E-14 | -0.312694243 |
| Mfge8 | 0.87 | 0.906 | 8.20608E-12 | -0.315426571 |
| Kif5b | 0.75 | 0.825 | 1.80672E-08 | -0.315554567 |
| Calm2 | 0.97 | 0.976 | 7.6076E-21 | -0.317733845 |
| Cpq | 0.53 | 0.646 | 8.99528E-07 | -0.318535245 |
| Tpm3 | 0.76 | 0.815 | 2.59993E-11 | -0.318648053 |
| Ywhah | 0.7 | 0.771 | 7.52889E-11 | -0.320637179 |
| Os9 | 0.53 | 0.624 | 5.521E-10 | -0.321683317 |
| Psmd2 | 0.47 | 0.562 | 1.37526E-12 | -0.32217444 |
| Capns1 | 0.92 | 0.959 | 2.98363E-16 | -0.322177912 |
| Rpn2 | 0.75 | 0.824 | 2.00047E-13 | -0.323240631 |
| Vapa | 0.68 | 0.775 | 3.51983E-12 | -0.324624264 |
| Cldnd1 | 0.51 | 0.617 | 3.01704E-09 | -0.329353843 |
| Rsu1 | 0.64 | 0.754 | 2.65475E-13 | -0.332785546 |
| Bcap31 | 0.59 | 0.71 | 2.65364E-13 | -0.334767356 |
| Lgmn | 0.51 | 0.607 | 8.08565E-10 | -0.335732467 |
| Tmbim6 | 0.87 | 0.933 | 2.46222E-19 | -0.335931756 |
| Tmem165 | 0.59 | 0.692 | 1.19593E-10 | -0.33792132 |
| Arl6ip5 | 0.7 | 0.782 | 4.73098E-14 | -0.339218359 |
| Tmed2 | 0.95 | 0.968 | 1.21162E-23 | -0.339362235 |
| Mdh1 | 0.58 | 0.698 | 1.29965E-13 | -0.340883242 |
| Tax1bp1 | 0.59 | 0.686 | 2.16029E-10 | -0.340992871 |
| Bzw1 | 0.61 | 0.702 | 3.99566E-14 | -0.343230608 |
| Asah1 | 0.5 | 0.638 | 2.83721E-13 | -0.343920417 |
| Calm1 | 0.96 | 0.982 | 5.735E-29 | -0.344225531 |
| Hnrnpk | 0.88 | 0.928 | 1.64619E-21 | -0.348979903 |
| Itm2b | 0.99 | 0.997 | 1.90198E-25 | -0.349996243 |
| Fbln2 | 0.94 | 0.955 | 7.29532E-13 | -0.350595664 |
| Laptm4a | 0.99 | 0.984 | 9.80337E-33 | -0.353473346 |
| Lman1 | 0.74 | 0.819 | 2.79777E-16 | -0.35547993 |
| Maged2 | 0.73 | 0.813 | 1.43972E-13 | -0.358025787 |
| Arf4 | 0.95 | 0.968 | 2.29491E-25 | -0.361015722 |
| Spcs2 | 0.69 | 0.792 | 7.00136E-19 | -0.366118717 |
| Capza2 | 0.65 | 0.764 | 5.94777E-17 | -0.37135551 |
| Itgb1 | 0.91 | 0.929 | 4.10053E-19 | -0.373537938 |
| Serpinf1 | 0.97 | 0.993 | 7.05656E-36 | -0.374935531 |
| Rsrp1 | 0.92 | 0.945 | 3.11059E-14 | -0.376088699 |
| Cthrc1 | 0.95 | 0.948 | 8.93085E-09 | -0.377549721 |
| Copb1 | 0.47 | 0.616 | 4.63734E-17 | -0.380252172 |
| Pam | 0.92 | 0.94 | 1.10264E-07 | -0.38285657 |
| Prdx1 | 0.97 | 0.986 | 3.13952E-42 | -0.384066043 |
| Ywhae | 0.82 | 0.866 | 4.1223E-25 | -0.386447499 |
| Dcn | 1 | 0.999 | 2.03487E-15 | -0.38712372 |
| Arpc2 | 0.89 | 0.944 | 2.63053E-28 | -0.389583359 |
| Sdc2 | 0.81 | 0.867 | 3.09089E-11 | -0.394951102 |
| Pcolce | 0.97 | 0.981 | 5.10577E-25 | -0.397722024 |
| Tmed9 | 0.94 | 0.965 | 2.59661E-32 | -0.397966801 |
| Eif4a1 | 0.87 | 0.917 | 3.67864E-27 | -0.399171567 |
| Ubb | 1 | 0.999 | 9.02065E-65 | -0.402427108 |
| Tmem59 | 0.92 | 0.97 | 1.55438E-28 | -0.408221431 |
| Hsp90b1 | 0.94 | 0.968 | 5.50496E-23 | -0.409059699 |
| Hspa5 | 0.92 | 0.963 | 3.52009E-25 | -0.424049837 |
| Pdia3 | 0.91 | 0.959 | 9.0292E-36 | -0.430897263 |
| Anxa1 | 0.81 | 0.89 | 1.17701E-18 | -0.433764339 |
| Aspn | 0.98 | 0.98 | 2.25051E-09 | -0.434144159 |
| Cd9 | 0.8 | 0.878 | 6.13987E-16 | -0.467408062 |
| Anxa5 | 0.93 | 0.974 | 1.85651E-41 | -0.469970716 |
| Cd63 | 0.98 | 0.997 | 6.98107E-60 | -0.482315427 |
| Cd81 | 0.99 | 0.996 | 5.28842E-48 | -0.494608542 |
| Pdia6 | 0.9 | 0.96 | 8.52236E-39 | -0.506659244 |
| Lum | 0.97 | 0.984 | 9.53977E-26 | -0.623632428 |
| Sept7 | 0.87 | 0.95 | 1.31652E-48 | -0.626455325 |
| Gsn | 0.96 | 0.987 | 1.98566E-17 | -0.74864955 |

**Table 2. List of differentially expressed genes in F-Act between PDGF-AB and PBS treated MI day7 hearts**

| **Gene** | **pct.1** | **pct.2** | **p_val_adj** | **Log2FoldChange** |
| --- | --- | --- | --- | --- |
| Eef1a1 | 1 | 0.999 | 1.10784E-36 | 0.465546727 |
| Pdia6 | 0.84 | 0.939 | 3.57297E-28 | -0.68409342 |
| Tpt1 | 0.99 | 0.988 | 4.67273E-27 | 0.430976115 |
| Cd63 | 0.93 | 0.994 | 1.71579E-26 | -0.443220913 |
| Anxa2 | 0.93 | 0.977 | 1.07986E-25 | -0.59254814 |
| Anxa5 | 0.94 | 0.981 | 3.8475E-25 | -0.521171728 |
| Hsp90b1 | 0.9 | 0.974 | 3.02427E-24 | -0.589160175 |
| Pdia3 | 0.82 | 0.952 | 7.7499E-24 | -0.557356628 |
| Sept7 | 0.82 | 0.924 | 9.28262E-24 | -0.611023171 |
| Prdx1 | 0.96 | 0.987 | 7.35127E-22 | -0.462791901 |
| Fau | 0.99 | 0.993 | 3.16457E-20 | 0.409098407 |
| Rpl23 | 1 | 0.991 | 7.03001E-19 | 0.385020536 |
| Rps12 | 0.97 | 0.961 | 2.09473E-18 | 0.454617893 |
| Myl12a | 0.97 | 0.985 | 2.39057E-18 | -0.447372846 |
| Itgb1 | 0.8 | 0.9 | 5.90599E-18 | -0.546741066 |
| Slc25a5 | 0.75 | 0.838 | 9.43192E-18 | -0.557985989 |
| Rps15 | 0.99 | 0.994 | 2.50043E-17 | 0.373674846 |
| Tmed9 | 0.9 | 0.948 | 4.69498E-17 | -0.434170203 |
| Pcolce | 0.95 | 0.987 | 1.17492E-16 | -0.404297837 |
| Anxa1 | 0.88 | 0.946 | 1.74848E-16 | -0.490069893 |
| Ywhae | 0.85 | 0.916 | 1.92432E-16 | -0.473664011 |
| Rpl21 | 1 | 0.99 | 2.01812E-16 | 0.333297419 |
| Calm1 | 0.98 | 0.98 | 3.49183E-16 | -0.426230748 |
| Rpl35a | 1 | 0.985 | 4.63563E-16 | 0.354195253 |
| Ldha | 0.85 | 0.917 | 6.03463E-16 | -0.528043575 |
| Rps25 | 0.99 | 0.99 | 1.54547E-15 | 0.364941458 |
| Eif4a1 | 0.87 | 0.939 | 1.62932E-15 | -0.452816586 |
| Rplp2 | 0.99 | 0.984 | 1.29785E-14 | 0.3683254 |
| Rpl10 | 0.99 | 0.996 | 1.73536E-14 | 0.323862523 |
| Rplp1 | 1 | 0.999 | 3.63601E-14 | 0.329166512 |
| Hspa5 | 0.92 | 0.959 | 5.04563E-14 | -0.446340524 |
| Ubb | 1 | 0.999 | 1.29511E-13 | -0.283380498 |
| Rpl8 | 1 | 0.999 | 1.65116E-13 | 0.297981191 |
| Rpl13a | 1 | 0.999 | 1.97578E-13 | 0.29443637 |
| mt-Nd4 | 0.99 | 0.987 | 2.13906E-13 | 0.374374574 |
| Rpl37a | 1 | 1 | 2.96891E-13 | 0.297503425 |
| Lmna | 0.86 | 0.92 | 1.15121E-12 | -0.427495415 |
| Ppib | 0.92 | 0.969 | 2.0812E-12 | -0.394397424 |
| Tpm3 | 0.68 | 0.82 | 4.06889E-12 | -0.463079254 |
| Bzw1 | 0.53 | 0.706 | 4.40169E-12 | -0.481236271 |
| Serpinh1 | 0.98 | 0.993 | 4.5654E-12 | -0.341151332 |
| Arl1 | 0.79 | 0.888 | 4.59396E-12 | -0.430184865 |
| Lman1 | 0.71 | 0.801 | 4.98604E-12 | -0.455627581 |
| Tmbim6 | 0.81 | 0.905 | 5.36163E-12 | -0.429830531 |
| Atp5b | 0.81 | 0.91 | 9.28321E-12 | -0.408579862 |
| mt-Nd1 | 1 | 0.994 | 1.06161E-11 | 0.310133264 |
| Hnrnpk | 0.86 | 0.932 | 1.25277E-11 | -0.376732507 |
| Rps21 | 0.91 | 0.854 | 1.65375E-11 | 0.413385515 |
| Arpc2 | 0.85 | 0.933 | 4.65742E-11 | -0.403844573 |
| Rpl39 | 0.97 | 0.978 | 5.79081E-11 | 0.306070692 |
| Tpm4 | 0.92 | 0.942 | 1.20205E-10 | -0.420500679 |
| Rps15a | 0.98 | 0.99 | 1.34132E-10 | 0.298577636 |
| Rpl6 | 1 | 0.99 | 1.48756E-10 | 0.318383171 |
| Calm2 | 0.94 | 0.962 | 1.6747E-10 | -0.349279952 |
| Rps24 | 0.99 | 0.984 | 2.40869E-10 | 0.314690978 |
| Rps26 | 0.97 | 0.942 | 2.80288E-10 | 0.370400877 |
| mt-Nd2 | 0.94 | 0.878 | 3.37218E-10 | 0.384360067 |
| Rpl28 | 0.99 | 0.978 | 5.74067E-10 | 0.310653531 |
| Nme1 | 0.7 | 0.811 | 7.25606E-10 | -0.445087694 |
| Psmb2 | 0.71 | 0.84 | 7.60465E-10 | -0.418025149 |
| Prdx4 | 0.59 | 0.745 | 9.04363E-10 | -0.463644179 |
| Egr1 | 0.85 | 0.748 | 9.70855E-10 | 0.567430062 |
| Rps20 | 0.98 | 0.984 | 1.40232E-09 | 0.304458307 |
| Prmt1 | 0.41 | 0.629 | 1.49883E-09 | -0.428287003 |
| Zbtb20 | 0.96 | 0.924 | 2.48417E-09 | 0.394135893 |
| Rpl22 | 0.96 | 0.932 | 2.86448E-09 | 0.325858127 |
| Ckb | 0.82 | 0.905 | 3.33781E-09 | -0.451224262 |
| Ranbp1 | 0.77 | 0.87 | 5.2413E-09 | -0.404447117 |
| Rpl26 | 1 | 0.997 | 5.89435E-09 | 0.262664358 |
| Arf4 | 0.9 | 0.945 | 6.49688E-09 | -0.373651696 |
| Ppa1 | 0.3 | 0.509 | 1.07871E-08 | -0.418177235 |
| Ostc | 0.84 | 0.875 | 1.10686E-08 | -0.353332895 |
| Fbln2 | 0.95 | 0.969 | 1.11607E-08 | -0.422416669 |
| Vps29 | 0.61 | 0.744 | 1.15876E-08 | -0.431340237 |
| Tmed2 | 0.95 | 0.967 | 1.2309E-08 | -0.33545454 |
| Rcn3 | 0.87 | 0.958 | 1.4442E-08 | -0.312271029 |
| Rpl10a | 0.99 | 0.984 | 1.62764E-08 | 0.293955018 |
| Rps8 | 1 | 0.996 | 2.41424E-08 | 0.26759592 |
| Rps4x | 1 | 0.991 | 2.89492E-08 | 0.275484515 |
| Erh | 0.84 | 0.902 | 3.29208E-08 | -0.379960773 |
| Rps10 | 0.97 | 0.952 | 3.52507E-08 | 0.320449823 |
| Hmgb1 | 0.94 | 0.945 | 5.57231E-08 | -0.376470369 |
| Pkm | 0.81 | 0.895 | 6.12379E-08 | -0.401460781 |
| Igfbp4 | 0.91 | 0.856 | 6.25981E-08 | 0.467177439 |
| Lgals1 | 1 | 1 | 7.3162E-08 | -0.365427103 |
| Anxa3 | 0.69 | 0.792 | 7.92627E-08 | -0.47017245 |
| Tagln2 | 0.93 | 0.943 | 8.49674E-08 | -0.397999199 |
| Reep5 | 0.76 | 0.856 | 1.03163E-07 | -0.353855889 |
| Rpl37 | 1 | 0.99 | 1.05265E-07 | 0.280095925 |
| Rpl27a | 0.99 | 0.999 | 1.31536E-07 | 0.252113451 |
| Kdelr3 | 0.66 | 0.799 | 1.56988E-07 | -0.37209575 |
| Rpl31 | 0.99 | 0.974 | 1.66934E-07 | 0.300224774 |
| Rbm3 | 0.76 | 0.846 | 1.91156E-07 | -0.44088107 |
| Rpl17 | 0.99 | 0.993 | 2.05094E-07 | 0.26230645 |
| Rpl29 | 0.97 | 0.961 | 3.0171E-07 | 0.319702825 |
| Serpinf1 | 0.9 | 0.965 | 3.29097E-07 | -0.277405063 |
| Fkbp10 | 0.55 | 0.691 | 4.25111E-07 | -0.421898626 |
| Capns1 | 0.92 | 0.939 | 4.64414E-07 | -0.282901872 |
| Actb | 0.99 | 0.988 | 6.10784E-07 | -0.418403663 |
| Eif2s1 | 0.35 | 0.531 | 8.37435E-07 | -0.391483663 |
| Hspa1a | 0.33 | 0.242 | 9.33273E-07 | 0.813719825 |
| Rpl38 | 0.96 | 0.923 | 1.2147E-06 | 0.333326384 |
| Cope | 0.79 | 0.882 | 1.21618E-06 | -0.355288415 |
| Rpl30 | 0.96 | 0.964 | 1.33361E-06 | 0.273100837 |
| Rpn2 | 0.68 | 0.806 | 1.3883E-06 | -0.377054744 |
| Ssr2 | 0.83 | 0.866 | 1.46294E-06 | -0.334667446 |
| Sec11a | 0.65 | 0.782 | 1.48875E-06 | -0.383102992 |
| Igfbp6 | 0.75 | 0.608 | 1.5745E-06 | 0.617832361 |
| Vcp | 0.72 | 0.834 | 2.15439E-06 | -0.350341899 |
| Ywhaq | 0.77 | 0.841 | 2.18848E-06 | -0.342402476 |
| Copb1 | 0.43 | 0.617 | 2.28583E-06 | -0.384315993 |
| Atp5f1 | 0.72 | 0.849 | 2.53021E-06 | -0.351568515 |
| Klf4 | 0.79 | 0.712 | 3.02967E-06 | 0.556575007 |
| Acta2 | 0.3 | 0.491 | 3.13919E-06 | -0.558283137 |
| Uqcrc1 | 0.54 | 0.687 | 3.16079E-06 | -0.38482005 |
| Spcs2 | 0.69 | 0.793 | 4.22245E-06 | -0.390550829 |
| Ubc | 0.91 | 0.951 | 4.34567E-06 | -0.343978273 |
| Pfn1 | 0.98 | 0.99 | 4.60504E-06 | -0.294376241 |
| Gpx7 | 0.7 | 0.817 | 6.42894E-06 | -0.448428786 |
| Ly6a | 0.82 | 0.69 | 7.45818E-06 | 0.543656323 |
| Eif6 | 0.5 | 0.656 | 7.86842E-06 | -0.39322806 |
| Tmem59 | 0.84 | 0.923 | 8.52035E-06 | -0.315478598 |
| Crispld2 | 0.7 | 0.626 | 8.65523E-06 | 0.466943472 |
| Fstl1 | 0.99 | 0.999 | 8.67251E-06 | -0.271073793 |
| Rpl36a | 0.98 | 0.953 | 8.80209E-06 | 0.282776868 |
| Mdh2 | 0.68 | 0.792 | 8.84669E-06 | -0.359434362 |

**Table 3. List of differentially expressed genes in M2MΦ between PDGF-AB and PBS treated MI day7 hearts**

| **Gene** | **pct.1** | **pct.2** | **p_val_adj** | **Log2FoldChange** |
| --- | --- | --- | --- | --- |
| Eef1a1 | 1 | 0.996 | 1.94313E-93 | 0.582952987 |
| Fau | 1 | 0.998 | 3.47314E-70 | 0.431276457 |
| Rpl35a | 1 | 0.992 | 2.73757E-62 | 0.434075723 |
| Rps15 | 1 | 0.987 | 2.12171E-59 | 0.464611177 |
| Rpl23 | 1 | 0.994 | 4.22842E-59 | 0.447987062 |
| Rps26 | 0.99 | 0.945 | 6.68492E-55 | 0.570531479 |
| Tpt1 | 1 | 0.988 | 3.72826E-48 | 0.443775398 |
| Rps12 | 1 | 0.959 | 4.3847E-47 | 0.5142511 |
| Rps24 | 1 | 0.993 | 1.33823E-45 | 0.415114111 |
| Rps15a | 1 | 0.99 | 2.8724E-45 | 0.445538328 |
| Rplp1 | 1 | 0.991 | 4.75207E-44 | 0.41567025 |
| Rplp2 | 1 | 0.989 | 2.53145E-43 | 0.410268562 |
| Rpl21 | 1 | 0.991 | 6.78664E-43 | 0.397589425 |
| Rpl28 | 0.99 | 0.968 | 7.19133E-40 | 0.448038944 |
| Rps8 | 1 | 0.994 | 7.30202E-40 | 0.399857742 |
| Rps25 | 1 | 0.987 | 6.11964E-39 | 0.367641302 |
| Rpl10 | 1 | 0.997 | 8.33737E-38 | 0.378426464 |
| Rps4x | 1 | 0.998 | 9.05809E-38 | 0.3671273 |
| Rpl30 | 0.98 | 0.925 | 1.18687E-35 | 0.431993822 |
| Rps16 | 1 | 0.994 | 4.36925E-32 | 0.324286602 |
| Rpl31 | 0.99 | 0.973 | 1.80782E-31 | 0.386667682 |
| Rpl29 | 0.99 | 0.956 | 1.94451E-31 | 0.42109792 |
| Rpl39 | 1 | 0.973 | 1.85776E-30 | 0.41924094 |
| Ucp2 | 0.9 | 0.761 | 4.55317E-30 | 0.5611097 |
| Rps10 | 0.99 | 0.965 | 1.99961E-29 | 0.398664913 |
| Rps13 | 0.99 | 0.99 | 1.99967E-29 | 0.343804552 |
| Rpl13a | 1 | 0.997 | 9.51868E-29 | 0.305589587 |
| Rps17 | 1 | 0.97 | 1.57806E-28 | 0.373936722 |
| Rps20 | 1 | 0.975 | 3.60004E-28 | 0.391082399 |
| Rps21 | 0.95 | 0.869 | 3.86219E-28 | 0.438363088 |
| Gnb2l1 | 0.96 | 0.871 | 1.23647E-27 | 0.456071942 |
| Lgmn | 0.99 | 0.994 | 3.06411E-27 | -0.403711143 |
| Rpl27a | 1 | 0.993 | 3.64488E-27 | 0.31652841 |
| Rpl26 | 1 | 0.993 | 9.32467E-27 | 0.29957778 |
| Rpl8 | 1 | 0.992 | 3.07066E-26 | 0.333645999 |
| Rpl12 | 0.97 | 0.888 | 1.2644E-25 | 0.437526949 |
| Rpl6 | 1 | 0.994 | 1.88789E-25 | 0.326812923 |
| Rpl11 | 1 | 0.994 | 2.64609E-24 | 0.301243131 |
| Rpl7 | 1 | 0.989 | 4.62482E-24 | 0.322458925 |
| Rpl23a | 1 | 0.99 | 6.55925E-23 | 0.279080607 |
| Rpl17 | 1 | 0.991 | 7.18388E-23 | 0.28740733 |
| Rpl19 | 1 | 0.994 | 2.8184E-22 | 0.297313558 |
| Rpl22 | 0.96 | 0.91 | 4.12721E-22 | 0.363027492 |
| Rpl7a | 0.99 | 0.935 | 9.90858E-21 | 0.381204352 |
| Hspa5 | 0.75 | 0.849 | 1.07249E-20 | -0.578890075 |
| Rsrp1 | 0.81 | 0.876 | 1.30141E-20 | -0.552312243 |
| Eef2 | 0.98 | 0.915 | 2.48925E-20 | 0.385693769 |
| Rps27a | 1 | 0.995 | 2.94424E-20 | 0.291068811 |
| Rps7 | 1 | 0.991 | 3.02335E-18 | 0.29320731 |
| Eif4a1 | 0.94 | 0.927 | 9.09033E-18 | -0.338271976 |
| Ctsc | 0.99 | 0.993 | 2.63952E-17 | -0.353353952 |
| Rpl34 | 1 | 0.987 | 6.04184E-17 | 0.272546469 |
| Tmbim6 | 0.9 | 0.921 | 6.47446E-17 | -0.329107775 |
| Rpl38 | 0.99 | 0.939 | 6.6259E-17 | 0.319842114 |
| Rpl18 | 1 | 0.992 | 6.70444E-17 | 0.261494532 |
| Anxa5 | 0.93 | 0.931 | 7.34261E-17 | -0.270884893 |
| mt-Nd4 | 0.99 | 0.971 | 7.54355E-17 | 0.323488883 |
| Rpl9 | 1 | 0.991 | 1.20732E-16 | 0.262387917 |
| Rps3a1 | 1 | 0.992 | 1.55706E-16 | 0.269434197 |
| Prdx1 | 0.97 | 0.962 | 5.96248E-16 | -0.251726312 |
| Eef1b2 | 0.88 | 0.72 | 1.05791E-15 | 0.405639692 |
| Rpl5 | 0.95 | 0.874 | 1.38002E-14 | 0.326024731 |
| Ubc | 0.99 | 0.985 | 1.62222E-14 | -0.428493967 |
| Naca | 0.98 | 0.938 | 3.07277E-14 | 0.287567577 |
| Rps2 | 1 | 0.99 | 3.69635E-14 | 0.307779573 |
| Cox7a2l | 0.95 | 0.863 | 7.34861E-13 | 0.284073229 |
| Capza2 | 0.86 | 0.891 | 7.97551E-13 | -0.3506062 |
| Rpl10a | 1 | 0.983 | 1.96079E-12 | 0.293340155 |
| Canx | 0.48 | 0.582 | 2.80561E-11 | -0.386718962 |
| Sept7 | 0.49 | 0.593 | 2.29145E-10 | -0.326747085 |
| Ms4a6d | 0.81 | 0.833 | 2.39598E-10 | -0.283467783 |
| Hsp90b1 | 0.7 | 0.758 | 2.80394E-10 | -0.349621933 |
| Pdia6 | 0.58 | 0.631 | 5.06592E-10 | -0.348467517 |
| Zfp36 | 0.8 | 0.679 | 5.12465E-10 | 0.3400825 |
| Psmd2 | 0.32 | 0.427 | 1.15069E-09 | -0.307667264 |
| Pdia3 | 0.7 | 0.737 | 2.88892E-09 | -0.289178925 |
| Mdh1 | 0.54 | 0.574 | 4.57707E-09 | -0.291152533 |
| Skap2 | 0.58 | 0.673 | 1.0824E-08 | -0.313896078 |
| Asah1 | 0.81 | 0.819 | 1.45863E-08 | -0.359264871 |
| Rpl4 | 1 | 0.981 | 2.07126E-08 | 0.260481447 |
| Rpl27 | 0.9 | 0.787 | 2.51689E-08 | 0.29771556 |
| Gsn | 0.81 | 0.818 | 2.6846E-08 | -0.299834743 |
| Rpsa | 0.99 | 0.946 | 1.82223E-07 | 0.264014886 |
| Bzw1 | 0.49 | 0.549 | 1.96914E-07 | -0.284201617 |
| Csf2ra | 0.64 | 0.695 | 3.51202E-07 | -0.342145594 |
| Atf3 | 0.83 | 0.744 | 5.05786E-07 | 0.30735749 |
| Ehd4 | 0.78 | 0.831 | 9.23916E-07 | -0.32156025 |
| Vcp | 0.53 | 0.59 | 1.72189E-06 | -0.291893868 |
| Vps35 | 0.4 | 0.455 | 5.02058E-06 | -0.284866128 |
| Spcs2 | 0.65 | 0.699 | 5.46628E-06 | -0.290473334 |
| Cd53 | 0.92 | 0.945 | 5.7134E-06 | -0.263776046 |
| Pla2g7 | 0.55 | 0.618 | 9.99466E-06 | -0.35226005 |

**Table 4. List of differentially expressed genes in F-SL between PDGF-AB and PBS treated MI day7 hearts**

| **Gene** | **pct.1** | **pct.2** | **p_val_adj** | **Log2FoldChange** |
| --- | --- | --- | --- | --- |
| Zfp36 | 0.54 | 0.35 | 7.80461E-06 | 0.694733279 |
| Eef1a1 | 1 | 1 | 1.64468E-32 | 0.60501355 |
| Rps15a | 0.98 | 0.97 | 4.3146E-27 | 0.578336104 |
| Rps12 | 0.94 | 0.88 | 1.20627E-13 | 0.559785684 |
| Rpl10a | 0.98 | 0.92 | 2.00034E-18 | 0.54672154 |
| Egr1 | 0.83 | 0.69 | 2.27636E-06 | 0.534262338 |
| Rpl39 | 0.96 | 0.86 | 2.0659E-11 | 0.520551081 |
| Rps25 | 0.98 | 0.94 | 8.54022E-18 | 0.51144446 |
| Rps24 | 0.97 | 0.95 | 7.00049E-16 | 0.507887448 |
| Rps15 | 0.99 | 0.95 | 1.23162E-16 | 0.501352313 |
| Rplp1 | 0.99 | 0.98 | 2.75798E-16 | 0.496334499 |
| Rps8 | 1 | 0.99 | 1.56919E-16 | 0.495223112 |
| Rpl8 | 0.99 | 0.98 | 9.73335E-17 | 0.490130499 |
| Rps20 | 0.98 | 0.9 | 2.34948E-11 | 0.489706958 |
| Rpl35a | 0.99 | 0.97 | 9.23632E-18 | 0.489328722 |
| Rps2 | 0.99 | 0.96 | 1.29638E-12 | 0.484276185 |
| Rpl12 | 0.9 | 0.75 | 2.55341E-07 | 0.483832195 |
| Rpl23 | 0.99 | 0.96 | 9.67853E-15 | 0.481255191 |
| Fau | 0.99 | 0.96 | 7.30181E-17 | 0.479772923 |
| Rplp2 | 0.97 | 0.93 | 5.43525E-14 | 0.477569163 |
| Rps4x | 1 | 1 | 2.465E-17 | 0.472336978 |
| Rpl29 | 0.9 | 0.8 | 0.000005372 | 0.4669417 |
| Rpl28 | 0.97 | 0.9 | 1.41385E-12 | 0.465681901 |
| Rpl21 | 0.99 | 0.95 | 3.12791E-13 | 0.463183983 |
| Rps10 | 0.91 | 0.83 | 0.000000207 | 0.461921368 |
| Rpl31 | 0.94 | 0.91 | 2.32129E-08 | 0.451117511 |
| Eef2 | 0.94 | 0.86 | 3.66439E-07 | 0.445383792 |
| Rpl6 | 0.99 | 0.97 | 6.10086E-12 | 0.444009658 |
| Rpl22 | 0.86 | 0.76 | 6.82129E-07 | 0.443059469 |
| Rpl36a | 0.96 | 0.9 | 2.12449E-07 | 0.439160986 |
| Rps17 | 0.97 | 0.91 | 0.000000005 | 0.430918528 |
| Rpl37 | 0.99 | 0.97 | 1.43863E-13 | 0.42759118 |
| Rpl17 | 0.98 | 0.97 | 7.82174E-13 | 0.426989952 |
| Rpl30 | 0.91 | 0.83 | 1.13263E-06 | 0.417599854 |
| Rpl32 | 1 | 0.98 | 4.87037E-12 | 0.41713688 |
| Rpl13a | 1 | 0.99 | 1.66849E-12 | 0.406020896 |
| Rps7 | 0.98 | 0.98 | 5.28699E-12 | 0.406002991 |
| Rpl37a | 0.99 | 0.98 | 2.27287E-11 | 0.401651355 |
| Rplp0 | 1 | 0.99 | 6.43689E-10 | 0.394874427 |
| Rpl24 | 0.99 | 0.96 | 4.82124E-09 | 0.389686431 |
| Tpt1 | 0.99 | 0.98 | 9.63554E-13 | 0.389125992 |
| Rps3a1 | 0.99 | 0.98 | 3.18355E-08 | 0.386020916 |
| Rps3 | 0.99 | 0.97 | 9.64136E-09 | 0.385407854 |
| Rpl26 | 0.99 | 0.98 | 4.81641E-10 | 0.385238459 |
| Rpl14 | 0.99 | 0.99 | 2.57094E-13 | 0.38394072 |
| Rpl7a | 0.96 | 0.87 | 6.74171E-06 | 0.375063972 |
| Rpl19 | 0.98 | 0.96 | 2.29752E-09 | 0.375015805 |
| Rps27 | 0.99 | 0.97 | 4.3919E-09 | 0.372127178 |
| Rps27a | 1 | 0.99 | 5.00838E-11 | 0.369816478 |
| Rps29 | 0.99 | 0.98 | 7.76317E-08 | 0.368366906 |
| Rpl7 | 0.98 | 0.95 | 9.55892E-08 | 0.3665426 |
| Rps16 | 1 | 0.99 | 8.29527E-11 | 0.362351584 |
| Rpl23a | 0.99 | 0.97 | 1.98764E-08 | 0.362175936 |
| Rpl34 | 0.99 | 0.96 | 9.48697E-07 | 0.350043005 |
| Rpl10 | 1 | 0.99 | 1.80675E-06 | 0.344950902 |
| Rps6 | 1 | 0.99 | 0.000007059 | 0.34215626 |
| Rpl9 | 1 | 0.99 | 4.45036E-10 | 0.341643368 |
| Rpl27a | 0.99 | 0.97 | 1.61573E-06 | 0.328550296 |
| Rpl11 | 0.97 | 0.97 | 0.000000046 | 0.311498887 |
| mt-Nd4 | 1 | 0.98 | 1.2585E-07 | 0.308208416 |
| Rps23 | 1 | 0.99 | 4.235E-08 | 0.287595851 |
| Rpl18a | 1 | 1 | 0.000000003 | 0.286661034 |
| Rps14 | 1 | 1 | 3.51189E-09 | 0.283058408 |
| Rpl13 | 1 | 1 | 0.000002589 | 0.271952993 |
| Ubb | 1 | 1 | 2.9939E-11 | -0.269764521 |
| Cd63 | 0.96 | 0.99 | 2.98587E-08 | -0.283805532 |
| Anxa5 | 0.92 | 0.96 | 1.67192E-06 | -0.285867521 |
| Prdx1 | 0.93 | 0.97 | 1.14879E-08 | -0.319029022 |
| Hspa5 | 0.79 | 0.86 | 0.000001027 | -0.39149414 |
| Hsp90b1 | 0.79 | 0.87 | 0.000000052 | -0.421800167 |
| Eif4a1 | 0.78 | 0.85 | 0.00000004 | -0.439814965 |
| Lum | 0.94 | 0.98 | 4.14778E-10 | -0.441675854 |
| Gsn | 1 | 1 | 1.53418E-09 | -0.528748585 |
| Pdia6 | 0.64 | 0.79 | 1.4368E-12 | -0.531175592 |
| Pdia3 | 0.69 | 0.83 | 4.04112E-14 | -0.540398927 |
| Sept7 | 0.68 | 0.84 | 7.82522E-18 | -0.656819333 |

**Detailed Methods and Materials**

***In vivo* animal studies:** All animal experiments were approved by the St Vincent’s Hospital and Garvan Institute of Medical Research Animal Ethics Committee. Mice were bred and housed in the BioCORE facility of the Victor Chang Cardiac Institute. Rooms were temperature and light/dark cycle controlled. Standard food was provided *ad libitum*. Soggy and/or high nutrient food was provided after surgery. Mouse strains used in experiments include C57Bl6J, *Pdgfra*-GFP, *Pdgfra*-merCremer and tdTomato (please see Key Resources Table for details). Both male and female mice aged 8-12 weeks (young) to 18-20 weeks (aged) were used. Surgical models included acute myocardial infarction (MI), catheterized minipump implantation, minipump implantation, tamoxifen treatment and oral gavage. Tissues were collected both from healthy and surgical mice for establishment of primary cell cultures, flow cytometry, immunohistochemistry and gene expression studies. Surgical procedures were performed in the BioCORE sugery suites between 9:30am to 3:30pm.

**Primary cell culture**: Primary cells were isolated from the mice and cultured at 37^o^C in 5% CO_2_ and normoxia as described in the relevant sections.

**Cardiac CFU-F isolation and long term culture:** Whole hearts were collected from mice that were either untreated, had undergone sham or MI surgeries, were treated with PDGF-AB ligand or vehicle via minipump and/or PDGFRα signaling inhibitor or vehicle via oral gavage, and cCFU-F assays performed as described in relevant sections. Once collected, hearts were minced and enzymatically digested as previously described (Chong, Chandrakanthan et al. 2011). The labelled cells were sorted by FACS for S^+^P^+^ population and plated at the density of 5000-cells/35cm^2^ dish or 20,000 cells/75cm^2^ flasks for colony long-term growth assays in 20% fetal bovine serum as described (Chong, Chandrakanthan et al. 2011). Colonies were scored as large, small and micro colonies according to their sizes following a crystal violet staining. For clonal analysis, single large, small and micro-colonies were trypsinised within O-ring Cloning Cylinders (Corning), and then cells were pooled, centrifuged and plated separately. Bulk cultured cells were split every 12 days until passage 4, then every 8 days thereafter. Cumulative growth curves were plotted on a logarithmic scale, following standard cell number calculations. In brief, cumulative cell numbers for each passage comprised the total cell number harvested, multiplied by the previous cumulative number divided by number of cells plated.

***In vitro* treatment of ligands, inhibitors and monoclonal antibodies**

Bulk or clone cultures of flow-sorted cardiac S^+^P^+^ cells isolated from healthy hearts were treated with PDGF-AB (50 or 100ng/ml) ligand, PDGFRα/β chemical inhibitor AG1296 (0.5-10µM), AKT (5-50nM) or cMYC (10-100μM) inhibitors, or control or PDGFRα-blocking monocolonal IgGs (0.02-10μg/ml), adding the reagents to culture media and refreshing with each media change.

**Cell cycle analysis, BrdU, TUNEL and mitochondrial assays**

Cells sorted from mouse hearts with and without *in vivo* or *in vitro* treatments were used for cell cycle analysis. Staining of the cells with PyroninY and 7AAD was performed with minor modifications to the previously described method (Joun, Fonoudi et al. 2017). In brief, cells were fixed in ice-cold 80% ethanol and stored at -20^o^C at least overnight. Cells were permeabilized by the Cytoperm Plus buffer (Becton Dickinson) for 5 mins and washed with 1x Cytoperm wash buffer before staining. Cellular DNA was stained with 7AAD (Becton Dickinson) and RNA with PyroninY (Sigma) before visualization by flow cytometry (Becton Dickinson, CantoII). Data analysis was performed using FlowJo software. BrdU, EdU and TUNEL assays were performed using the APC-BrdU assay kit (Becton Dickinson), the Click-iT EdU assay kit (Thermofisher Scientific) and the DeadEnd™ Colorimetric TUNEL System (Promega), respectively, following manufacturers instructions. For mitochondrial assays, cells were harvested following standard methods, washed once with PBS and resuspended in culture medium containing 200nM of Mitotracker Green FM (Thermofisher Scientific). The staining mix was incubated for 15min at 37^o^C and analysed on a FACS CantoII (Becton Dickinson).

**Live cell imaging and single-cell tracking of cCFU-F**

Live cell imaging was performed using a Leica live cell imaging microscope (DMI6000B) equipped with x-y-z controller and hardware autofocus as described (Cornwell, Hallett et al. 2016). Phase contrast images were acquired every 15 minutes for 12 days (288 hours). Raw images were exported to Matlab for removal of background noise, enhancement of contrast, and to stitch contiguous frames of view using custom-written scripts. Custom-written software implemented in Matlab (Nordon’s Tracking Tool) was used to manually track cell nuclei through consecutive frames, to build trajectories and record fate outcomes (division, death, and right censored [lost or not recorded]). At least 200 cells from at least 20 individual clones per condition were tracked from two separate wells. Statistical analyses of live cell tracking data were implemented in custom-written scripts in Matlab. Kaplan-Meier analysis was used to generate empirical cumulative distribution functions for cell division. Cox regression was used to test for group differences and to estimate the relative risk of division relative to untreated controls. The relative frequency of observed fates in each treatment group was quantified by counting the number of cells with distinct fate outcomes in each group. Student’s *t*-tests were used to test for significant differences in the proportion of observed fates in each treatment group. All Matlab code will be made available upon request.

**Quantitative RT-PCR, Western blots**

Quantitative RT-PCR, was performed using the SYBR master mix (Roche) following manufacturer’s instructions. PCR reactions were run on LightCycler480 (Roche). Expression values were calculated using the Delta Ct method. Western blotting and was performed following standard protocols. For a comprehensive list of primers and antibodies used, refer to the “Key Resources Table”.

**Microarray sample preparation and analysis**

For studies with inhibitor and ligand (P0), three independent experiments were performed for each condition. Total RNA was extracted using RNAeasy micro RNA isolation kit (Qiagen) and the quality of RNA was assessed using the Nanodrop (Thermofisher Scientific; 230/260>1.8, 260/280>1.8) and Bioanalyzer (Agilent, RIN>8.0). The RNA was further processed for microarray analysis by the Ramaciotti Center for Genomics (University of New South Wales, Sydney). Total RNA (100ng) of was labelled using the Affymetrix WT kit (Ambion). 3.52 μg of fragmented labelled ssDNA was hybridised to Mouse Gene 1.0 ST Gene Arrays for 16 hours at 45°C in the Affymetrix GeneChip Oven. Arrays were washed on the Affymetrix Fluidics 450 instrument as per manufacturer’s protocol using the Affymetrix GeneChip Hybridization Wash & Stain kit for cartridge arrays. Arrays were scanned on the Affymetrix GeneChip 3000 7G Scanner instrument as per manufacturer’s protocol and the Affymetrix GeneChip Command Console (AGCC) software was used to generate the CEL files from the DAT image files. RAW and normalised microarray data are available at NCBI’s Gene Expression Omnibus (GSE106778; GSE106779) (Edgar, Domrachev et al. 2002)

Microarray data were processed and analyzed in R/Bioconductor (Gentleman, Carey et al. 2004) Heatmaps were generated using the gplots R package and where clusters were identified, were split by cutting the tree at the reported height. Gene ontology analyses were performed using DAVID (Huang da, Sherman et al. 2009) against the *Mus musculus* background. Quality control was first performed using the ‘affy’ and ‘affyPLM’ R packages to assess the distribution of probe-level effects via the RLE and PLM methods as well as pseudoimage decomposition of the microarrays (Gautier, Cope et al. 2004). No artefacts or outliers were revealed. Data were then normalised at the transcript level using Robust Multichip Average (RMA) normalisation. Principle component analysis was performed using the pcaMethods R package (Stacklies, Redestig et al. 2007) using singular value decomposition. For determination of differentially expressed genes, a linear model with Bayes variance shrinkage was fit using the limma R package (Smyth 2005) to determine moderated t-statistics for all pair-wise comparisons. P-values were then corrected for multiple hypothesis testing using the Benjamini and Hochberg method (Benjamini and Hochberg 1995). Unless reported otherwise, a corrected p-value less than or equal to 0.05 was reported as significant. Where multiple probe sets were assigned to the same annotated gene, the gene with the highest average expression was reported for further analyses.

**Immunohistochemistry and confocal microscopy in tissues**
Cryosections (8µm) of paraformaldehyde (PFA; 4%) perfusion fixed and sucrose treated hearts were cut and incubated with one or more primary antibodies overnight at 4^o^C (listed in Table S2) after incubating with a blocking solution including 1% bovine serum albumin, 3% serum (from the species that the secondary antibodies were derived), 0.03% triton X-100 in PBS for 30 minutes at room temperature. Nuclear counter-staining was performed after visualization of the markers by incubating sections with appropriate fluorochrome-conjugated secondary antibodies. For a comprehensive list of antibodies used, refer to the “Key Resources Table”.

**Immunohistochemistry and confocal microscopy on cells**
Cells were fixed with 4% PFA for 5 minutes at room temperature, washed with PBS and incubated with one or more primary antibodies for 1 hour at 4^o^C after incubating with a blocking solution including 1% bovine serum albumin, 3% serum (from the species that the secondary antibodies were derived), 0.03% triton X-100 in PBS for 30 minutes at room temperature. Nuclear Hoechst counter-stains performed after visualization of the markers by incubating the sections with appropriate fluorochrome-conjugated secondary antibodies. Images were acquired by using a Zeiss upright microscope or Zeiss confocal microscope. For a comprehensive list of antibodies used, refer to the “Key Resources Table”.

**Systemic delivery of ligands, inhibitors and monoclonal antibodies**

PDGF-AB ligand (R&D, 20µg/ml) or vehicle (PBS) was administered systemically by a subcutaneous minipump (Alzet), inserted to the right external jugular vein via 13mm of 28G catheter (Alzet) through a 1mm incision. The ligand/vehicle was continuously released at a concentration of 0.5µl/hr for 5 days. Imatinib was delivered via oral gavage once daily at a concentration of 20mg/kg daily for 5 days. To block PDGFRα *in vivo,* mice were injected with one dose of 400ug (2μg/ul) on day 1, 200μg each on days 4 and 8, before either MI or sham surgeries.

**Acute myocardial infarction**

Mice were anaesthetised by intraperitoneal injection of ketamine (100mg/kg) and xylazine (20mg/kg), and intratracheally intubated. The hearts were exposed via a left intercostal incision, the left anterior descending coronary arteries were ligated just before the first diagonal branching point, lungs inflated and the wounds suture closed. Sham operations were performed without the ligation of the coronary artery. The mice were given intra-muscular injection of Bupernopherine for 3 days. Hearts were collected for immuno-histochemistry, FACS analysis or echocardiography 5-28 days after surgery as indicated for individual experiments.

**Chronic cardiac injury models**

*Aldosterone mediated cardiac fibrosis***:** Anaesthetised mice were implanted with a 14 day mini-pump (Alzet 1002*)* containing either aldosterone (0.15g/hr) or vehicle (polyethylene glycol) through a dorsal skin incision (Kageyama and Bravo 1988). Mice were fed with water containing 1% NaCl**.** The osmotic pumps were replaced after 2 weeks and hearts analyzed on 28 days. Excess aldosterone generates hypervolemic hypertension and cardiac fibrosis by stimulating water and sodium reabsorption, and release of vasopressin (Brunner, Chang et al. 1972).

*Aortic banding assays:* The aorta was exposed through a mediastinum incision in anaesthetised and intubated mice. The aortic root was isolated and a 4-0 constrictor suture was placed by using a template wire for diameter control. The incision was then closed and the animals recovered. Hearts were analysed at day 28 post surgery.

**Echocardiography**

Mice were anaesthesized using 5% isoflurane delivered by nose cone and echocardiography was performed on supine mice using a Vevo770 (Visualsonics Inc., Toronto, Canada) under 1-2% isoflurane on a warming pad set to 37°C. B-mode images were taken of the LV long axis (LAX) of the heart followed by four short axis (SAX) images, starting at the mid-papillary muscle level and at 1, 2 and 3mm towards the apex. Using Vevo770 Workstation software the following data were obtained by a blinded operator from the echocardiography images: LV length was measured from the apical dimple to the base of the aortic valve leaflets from the LAX images at end diastole (ED) and end-systole (ES). The areas boarded by the endocardium were planimetrically measured on all four SAX images at ED and ES, and from this a modified Bullet formula was used to derive ED and ES LV volumes (LVvol = L x (mean endocardial area) x 5/6).

**Quantification of cellularity and vascularity**

CD31 and DAPI stained sections of infarcted and sham-operated hearts were imaged using Zeiss 700 upright confocal microscope at 20 times magnification to cover entire left ventricle. Images were segmented as infarct, border and remote zones. The images were imported into the Image J to analyze total cell and vessels numbers in the respective areas.

**Heart weight/body weight and infarct size measurement**

Mice were weighed postmortem at post-surgical day 28, then hearts were resected and weighed separately. Percentage heart weigh/body weight was calculated. Hearts were embedded in OCT and frozen. 10μM sections were stained using a Masson’s Trichrome staining kit (Sigma) and stained slides were scanned using an Aperio ePathology scanner (Leica). The area size and length of the left ventricles were measured using Image J.

**Flow cytometry analysis of immune cell influx and time course analysis of GFP^medium^ population**

For immune influx analysis, the total interstitial population isolated by FACS was subjected to further fractionation with monocyte makers. Within the monocyte/macrophage population (CD45^+^ CD11b^+^ Ly-6G^low^), type 1 monocytes (m1) were defined as Ly-6C^high^[F4/80/I-A^b^/CD11c]^low^, and monocyte type 2 (m2) as Ly-6C^low^[F4/80/I-A^b^/CD11c]^low^ (see Fig. S6A,B). In time course analyses (2 hrs to 5 days), *Pdgfra*-GFP mice were used to separately quantify the *Pdgfra*-GFP^high^ stromal cells and *Pdgfra*-GFP^medium^ myofibroblasts within the S^+^P^+^ population in sham and MI hearts (see Fig. 5G).

***In vivo* labelling of BrdU and tamoxifen induction**

BrdU (50mg/ml) was delivered through subcutaneously implanted minipumps. 7-day pumps with a delivery rate of 0.5µl/ml were used for the 5 day assays. For labelling of the PDGFRα lineage positive cells *in vivo*, adult *Pdgfra-merCREmer x tdTomato* mice were given tamoxifen (40mg/ml, Sigma) via oral gavage (300mg/kg) for 2 days, MI surgeries were performed 1 week after the tamoxifen treatment, and hearts were subjected to IHC or FACS analysis at day 5 and day 28 post-MI surgeries.

**Production of APA5 antibody**

The blocking monoclonal antibody against PDGFRα was produced in house (Centre For Targeted Therapy, Garvan Institute of Medical Research) from the APA-5 hybridoma cell line (Takakura, Yoshida et al. 1996, Takakura, Yoshida et al. 1997). IgG was purified from the supernatant using Protein L Sepharose resin (Genscript) as previously described (Dudgeon, Rouet et al. 2012)and concentrated to 2mg/mL in PBS using an Amicon Ultra-15 Centrifugal filter unit (Millipore). The purity of the antibody preparation was validated by reducing and non-reducing SDS-PAGE and endotoxin contamination was assessed using the QCL-1000™ Endpoint Chromogenic LAL Assay (Lonza) according to manufacturer’s instructions. Control IgG from Rat IgG2a isotype control clone 2A3 was sourced from BioXcell.

**ELIZA assay**

Serum was collected from 8wk old healthy and PDGF-AB-treated mice. Briefly, blood was collected via cardiac puncture after euthanasia and let clot at room temperature for 30mins before centrifugation at 1500rpm for 10mins. Serum concentration of mouse PDGF-AB and human PDGF-AB was measured using a Mouse/Rat PDGF-AB Quantikine ELISA Kit and a Human PDGF-AB Quantikine ELISA Kit (R&D Systems), respectively, following manufacturer’s instructions and analysis was performed using the ELISA Centro Luminescence Microplate Reader (Bio-Rad).

**Single cell transcriptomics** The single cell library preparation was performed using a commercially available droplet method, the 10x Chromium System (10x Genomics Inc. San Francisco, CA). The number of cells loaded on the system was calculated based on the desired number of captured cells (5000 per-sample) following manufacturer’s instructions. scRNA-seq libraries were generated following capture. Samples were sequenced on the Illumina NovaSeq 6000 platform.

### **Processing of 10x Genomics Chromium scRNA-seq data** Raw scRNA-seq data was processed using the 10x Genomics CellRanger software (version 2.2.0). The BCL files obtained from the Illumina NovaSeq platform were processed to Fastq files using the Cell Ranger *mkfastq* program. The Fastq files were then mapped to the mm10 version 1.2.0 reference, downloaded from the 10x Genomics website. The Cell Ranger *count* program was run on individual Fastq data-sets from the different conditions. The *aggr* program was run to generate an aggregate count matrix for the 8 individual samples.

### **Filtering,** **dimensionality reduction and clustering of scRNA-seq data** Bioinformatics processing of the scRNA-seq data was performed in R using the *Seurat* package (Butler, Hoffman et al. 2018) and visualisation performed with ggplot2 (Wickam 2016). R code containing the steps used for processing and clustering the aggregate scRNA-seq data can be provided on request**.** Initial quality control filtering metrics were applied as follows: Cells with fewer than 200 detected expressed genes were filtered out. Genes that were expressed in less that 10 cells were filtered out. In order to control for dead or damaged cells, cells with over 10% of raw unique molecular identifiers (UMIs) mapping to mitochondrial genes were filtered out. To further control for potential doublets in our data, we visualized the distribution of expressed genes and unique molecular identifier (UMI) numbers and filtered out cells with clear outliers.

UMIs were normalized to counts-per-ten-thousand, log-transformed, and a set of 2018 highly variable genes was identified by gating for mean expression level and dispersion level. The log-normalised data was scaled, with variation due to total number of UMIs regressed out using a linear model. Principal component analysis was run on the scaled data for the set of previously-defined highly variable genes. In order to identify the number of principal components (PCs) to use for clustering, we ran the JackStraw procedure implemented in *Seurat* that identifies statistically significant PCs. Based on running the JackStraw procedure with 1,000 permutations, we defined 64 significant PCs as those up to P<0.001, which were used as input to the *Seurat* graph-based clustering program, *FindClusters*. The resolution parameter for *FindClusters,* which determined the number of returned clusters, was set at 0.8 after considering clustering output from a range of resolutions. The cells and clusters were visualized on a t-SNE dimensionality reduction plot generated on the same set of 64 PCs used for clustering.

We inspected the clusters for hybrid gene expression signatures that could indicate captured cell doublets, contaminating cell types, or a signature of stress/apoptosis that could indicate cells damaged during the process of cell sorting and capture in the microfluidics device. Within the scRNA-seq data, our initial clustering analysis returned 26 clusters; within these we identified 1 minor cluster exhibiting hybrid gene expression signatures (Fibroblast-endothelial cell intermediate; F-EC) and a minor cluster indicating an erythroid identify. These two clusters were removed and all subsequent analysis (e.g. differential expression analysis) was performed on the remaining 24 clusters.

### **Differential expression** For calculating DE, we first identified genes expressed in at least 25% of cells for at least one of the conditions being compared and with an absolute log2 fold-change difference of 0.25 (including a pseudo-count of 1). We then assigned P-values using the ‘MAST’ test for DE (Finak, McDavid et al. 2015) implemented in the *Seurat FindMarkers* program. A Bonferroni-adjusted P-value of 1e-05 was used to determine significantly DE genes.

### **Gene Ontology testing** Over-representation of GO terms in gene lists was calculated using the PANTHER web-service (Mi, Huang et al. 2017). The set of expressed genes from the scRNA-seq was used as background. A false-discovery rate cut-off of 0.05 was used to determine statistical significance.

### **Network analysis** Network analysis of DEGs was performed using protein-protein association connections obtained from the STRING (Szklarczyk, Morris et al. 2017) version 11 data-base, considering connections with a combined score of at least 700. Differentially expressed genes between PDGF-AB and PBS conditions within sub-populations MYO, M2MΦ, F-Act and F-SL were used as input to the STRING web-server, with network connections visualised using Cytoscape (Shannon, Markiel et al. 2003).

**Quantification and statistical analysis**

Statistical analyses of live cell tracking data were implemented in custom-written scripts in Matlab. Kaplan-Meier (KM) analysis was used to generate empirical cumulative distribution functions for cell division. Cox regression was used to test for group differences and to estimate the relative risk of division (relative to untreated controls). The relative frequency of observed fates in each treatment group was quantified by counting the number of cells with distinct fate outcomes in each group. Student’s *t*-tests were used to test for significant differences in the proportion of observed fates in each treatment group. All Matlab code will be made available upon request. For all other data, Student’s *t*-tests were used to determine significance between triplicate experiments.

**Data and software availability**

RAW and normalised microarray data are available at NCBI’s Gene Expression Omnibus (GSE106778; GSE106779) (Edgar, Domrachev et al. 2002).

Raw and processed PDGF scRNA-seq data is available on ArrayExpress under accession ID E-MTAB-7971 with username Reviewer_E-MTAB-7971 and password dgbcctze.

**Key Resources Table**

| **REAGENT or RESOURCE** | **SOURCE** | **IDENTIFIER** |
| --- | --- | --- |
| Antibodies | | |
| Rb | Abcam | ab6075-1 |
| PhosphoRb (Ser780) | Cell Signaling | D59B7 |
| P130 | Abcam | 76234 |
| P27^kip^ | BD Biosciences | 610242 |
| H2B | Abcam | Ab18977 |
| CyclinA2 | Cell Signaling | 4656P |
| CyclinB1 | Cell Signaling | 4138P |
| CyclinE1 | Cell Signaling | 4129P |
| CyclinD1 | Cell Signaling | 2978P |
| c-Myc | Santa Cruz | sc-788 |
| Akt (pan-Akt) | Cell Signaling | 4691 |
| p-Akt308 | Cell Signaling | 2965 |
| p-Akt473 | Cell Signaling | 4060 |
| p-FOXO1 | Cell Signaling | 9461 |
| FOXO1 | Cell Signaling | 2880 |
| SMA | Sigma-Aldrich | A2547 |
| Calponin | Abcam | ab46794 |
| BrdU | Abcam | ab6326 |
| PDGFRα | Cell Signaling | 3164 |
| PDGFRβ | Santa Cruz | Sc-432 |
| Lectin | Sigma-Aldrich | L3759 |
| Isolectin | Thermo Fisher Scientific | I21411 |
| Collagen IV | Abcam | ab19808 |
| Collagen VI | Abcam | ab6588 |
| cTnT | Thermo Fisher Scientific | MS-295-P1 |
| Ly-6C PE | eBioscience | 12-5932-80 |
| F4/80 FITC | eBioscience | 11-4801 |
| CD11c FITC | BD Biosciences | 553953 |
| I-A^b^ FITC | BD Biosciences | 553456 |
| PE Sca1 (Ly6A/E) | BD Biosciences | 5531108 |
| CD45- APCCy7 | BD Biosciences | 557659 |
| CD31-PECy7 | eBioscience | 25-0311-82 |
| PDGFRα (CD140a)-APC | eBioscience | 17-1401-81 |
| Ly-6G (Gr-1)-PerCPCy5 | BD Biosciences | 45-5931-80 |
| CD11b-APC | BD Biosciences | 553312 |
| mPDGFRa | R&D System | AF1042 |
| mPDGFRb | R&D System | AF1062 |
| CD146 | Abcam | Ab75769 |
| NG2 | EMD Millipore | AB5320 |
| **Chemicals, Peptides, and Recombinant Proteins** | | |
| PDGF-AB | R&D System | 222-AB |
| AG1296 | Cayman Chemical | 146535-11-7 |
| Imatinib | LC Laboratories | I-5508 |
| cMyc inhibitor (10058-F4) | Sigma-Aldrich | F3680 |
| Akt inhibitor (LY294002) | Cell Signaling | 9901 |
| PyroninY | Polysciences | 18614-5 |
| 7AAD | BD Biosciences | 51-68981E |
| Mitotracker Green FM | Thermo Fisher Scientific | M7514 |
| Hoechst | Invitrogen | H369 |
| DAPI | Thermo Fisher Scientific | 62247 |
| Aldosterone | Sigma-Aldrich | 234362 |
| Tamoxifen | Sigma-Aldrich | T5648 |
| BrdU | Sigma-Aldrich | B5002 |
| **Critical Commercial Assays** | | |
| Click-iT EdU assay kit | Thermo Fisher Scientific | C10337 |
| DeadEnd™ Colorimetric TUNEL System | Promega | G736 |
| Masson Trichrome Staining kit | Sigma-Aldrich | HT-15 |
| Mouse/Rat PDGF-AB Quantikine ELISA Kit | R&D Systems | MHD00 |
| Human PDGF-AB Quantikine ELISA Kit | R&D Systems | DHD00C |
| **Deposited Data** | | |
| Microarray data repository | https://www.ncbi.nlm.nih.gov/geo/query/acc.cgi?acc=GSE106779- reviewer token: ifczsmwwhzkfxwb |  |
| Single-cell RNA-seq data repository | [https://protect-au.mimecast.com/s/Fu09CE8kz9tX5GJRcNUCuO?domain=ebi.ac.uk](https://protect-au.mimecast.com/s/Fu09CE8kz9tX5GJRcNUCuO?domain=ebi.ac.uk" \t "_blank). ID E-MTAB-7971 |  |
| **Experimental mice** | | |
| *Mouse, Pdgfra-*GFP, KI GFP, KO | Jackson Laboratory | *Pdgfra ^tm11(EGFP)Sor^* |
| *Mouse, Pdgfra^fl^,* TG | Jackson Laboratory | B6.Cg-Pdgfratm8Sor/EiJ |
| *Mouse, PdRa-MCM,* KI | RIKEN Center for Developmental Biology, Kobe, Japan | *Pdgfra-merCremer* |
| *Mouse, tdTomato Red,* TG | Jackson Laboratory (Imported from ARMI) | *B6.Cg-Gt(ROSA)26Sor^tm14(CAG-tdTomato)Hze^/J* |
| **Primers** | | |

*PolG*	F:CACGGACCTCCTGCCTAAG	R:TCATCTAGTCGAGGGGCAGAG

*PolG2*	F:AGCCAATAGAAACCCTGTGG	R:CACTTACAGACAGAACGCAACG

*Tfam*	F:TCAGGAGGCAGTTATTG	R:CAGGAAGTCTTCACGTTGACC

*Smooth Muscle Actin*	F:GCTGACAGAGGCACCACTGA	R:CATCTCCAGAGTCCAGCACA

*Calponin*	F:GAACCCTGTGGACCTGTTTGA	R:GCGTCGTCAAAGTTCCTCTC

| *Embryonic Smooth Muscle* F:CGCTTTGGCAAGTTTATCCG R:CAGCACTGAAGACACGACTT | | |
| --- | --- | --- |
| *Fibronectin (EIIA)* F:CCCACTGTGGAGTACGTGG R:GAGTCCTGACACAATCACCG | | |
| *Fibronectin (EIIB)* F:GGGGACCTCTCTGGAAGAAGTGG R:GTCCCAGGCAGGAGATTTG | | |
| *Nkx2-5* F:CCCAAGTGCTCTCCTGCTTTC R:TCCAGCTCCACTGCCTTCTG | | |
| *Islet-1* F:TCATCCGAGTGTGGTTTCAA R:CCATCATGTCTCTCCGGACT | | |
| *Tbx-5* F:ATGGTCCGTAACTGGCAAAG R:TTTCGTCTGCTTTCACGATG | | |
| *Gata-4* F:GCCCGGGCTGTCATCTCACTATG R:GCTGGCCTGCGATGTCTGAGTG | | |
| *Tbx-20* F:GCAGCAGAGAACACCATCAA R:CAGAGCTCCTTCGTTTCCAG | | |
| *DDR-2* F:CTGTCGGATGAGCAGGTTAT R:CTCGGCTCCTTGCTGAAGAA | | |
| *Vimentin* F:TACCAGGACACTATTGGCCG R:CTGTTGCACCAAGTGTGTGC | | |
| *Desmin* F:GCTATCAGGACAACATTGCG R:GTTGTTGCTGTGTAGCCTCG | | |
| *18S rRNA* F:AGGGGAGAGCGGGTAAGAGA R:GGACAGGACTAGGCGGAACA | | |
| *GAPDH* F:GGTCCTCAGTGTAGCCCAAG R:AATGTGTCCGTCGTGGATCT | | |
| *HPRT* F:AGTGTTGGATACAGGCCAGAC R:CGTGATTCAAATCCCTGAAGT | | |
| Software and Algorithms | | |
| CellRanger | 10x Genomics | https://support.10xgenomics.com/single-cell-gene-expression/software/downloads/latest |
| Seurat | PMID: 29608179 | <https://satijalab.org/seurat/>; RRID: SCR_007322 |
| PANTHER | PMID: 27899595 | <http://www.pantherdb.org>; RRID:SCR_004869 |
| Cytoscape | PMID: 14597658 | <https://cytoscape.org>; RRID:SCR_003032 |
| Differential Proportion Analysis | PMID: 30912746 | https://elifesciences.org/articles/43882 |

**References of the supplementary materials**

Benjamini, Y. and Y. Hochberg (1995). "Controlling the false discovery rate: a practical and powerful approach to multiple testing." Journal of the Royal Statistical Society, Series B **57**: 289-300.

Brunner, H. R., P. Chang, R. Wallach, J. E. Sealey and J. H. Laragh (1972). "Angiotensin II vascular receptors: their avidity in relationship to sodium balance, the autonomic nervous system, and hypertension." J Clin Invest **51**(1): 58-67.

Butler, A., P. Hoffman, P. Smibert, E. Papalexi and R. Satija (2018). "Integrating single-cell transcriptomic data across different conditions, technologies, and species." Nat Biotechnol **36**(5): 411-420.

Chong, J. J., V. Chandrakanthan, M. Xaymardan, N. S. Asli, J. Li, I. Ahmed, C. Heffernan, M. K. Menon, C. J. Scarlett, A. Rashidianfar, C. Biben, H. Zoellner, E. K. Colvin, J. E. Pimanda, A. V. Biankin, B. Zhou, W. T. Pu, O. W. Prall and R. P. Harvey (2011). "Adult cardiac-resident MSC-like stem cells with a proepicardial origin." Cell Stem Cell **9**(6): 527-540.

Cornwell, J. A., R. M. Hallett, S. A. der Mauer, A. Motazedian, T. Schroeder, J. S. Draper, R. P. Harvey and R. E. Nordon (2016). "Quantifying intrinsic and extrinsic control of single-cell fates in cancer and stem/progenitor cell pedigrees with competing risks analysis." Scientific Reports **6**: 27100.

Cornwell, J. A., R. E. Nordon and R. P. Harvey (2018). "Analysis of cardiac stem cell self-renewal dynamics in serum-free medium by single cell lineage tracking." Stem Cell Research **28**: 115-124.

Dudgeon, K., R. Rouet, I. Kokmeijer, P. Schofield, J. Stolp, D. Langley, D. Stock and D. Christ (2012). "General strategy for the generation of human antibody variable domains with increased aggregation resistance." Proc Natl Acad Sci U S A **109**(27): 10879-10884.

Edgar, R., M. Domrachev and A. E. Lash (2002). "Gene Expression Omnibus: NCBI gene expression and hybridization array data repository." Nucleic Acids Res **30**(1): 207-210.

Finak, G., A. McDavid, M. Yajima, J. Deng, V. Gersuk, A. K. Shalek, C. K. Slichter, H. W. Miller, M. J. McElrath, M. Prlic, P. S. Linsley and R. Gottardo (2015). "MAST: a flexible statistical framework for assessing transcriptional changes and characterizing heterogeneity in single-cell RNA sequencing data." Genome Biol **16**: 278.

Furtado, M. B., M. W. Costa, E. A. Pranoto, E. Salimova, A. R. Pinto, N. T. Lam, A. Park, P. Snider, A. Chandran, R. P. Harvey, R. Boyd, S. J. Conway, J. Pearson, D. M. Kaye and N. A. Rosenthal (2014). "Cardiogenic genes expressed in cardiac fibroblasts contribute to heart development and repair." Circulation Research **114**(9): 1422-1434.

Gautier, L., L. Cope, B. M. Bolstad and R. A. Irizarry (2004). "affy--analysis of Affymetrix GeneChip data at the probe level." Bioinformatics **20**(3): 307-315.

Gentleman, R. C., V. J. Carey, D. M. Bates, B. Bolstad, M. Dettling, S. Dudoit, B. Ellis, L. Gautier, Y. Ge, J. Gentry, K. Hornik, T. Hothorn, W. Huber, S. Iacus, R. Irizarry, F. Leisch, C. Li, M. Maechler, A. J. Rossini, G. Sawitzki, C. Smith, G. Smyth, L. Tierney, J. Y. Yang and J. Zhang (2004). "Bioconductor: open software development for computational biology and bioinformatics." Genome Biol **5**(10): R80.

Huang da, W., B. T. Sherman and R. A. Lempicki (2009). "Systematic and integrative analysis of large gene lists using DAVID bioinformatics resources." Nat Protoc **4**(1): 44-57.

Joun, G., H. Fonoudi and N. S. Asli (2017). "Cancer Cell Properties Shape Along Metabolic Activity in Human Epithelial Carcinoma." Journal of Stem Cell: Advanced Research and Therapy **4**: 1-5.

Kageyama, Y. and E. L. Bravo (1988). "Hypertensive mechanisms associated with centrally administered aldosterone in dogs." Hypertension **11**(6 Pt 2): 750-753.

Mi, H., X. Huang, A. Muruganujan, H. Tang, C. Mills, D. Kang and P. D. Thomas (2017). "PANTHER version 11: expanded annotation data from Gene Ontology and Reactome pathways, and data analysis tool enhancements." Nucleic Acids Res **45**(D1): D183-D189.

Noseda, M., M. Harada, S. McSweeney, T. Leja, E. Belian, D. J. Stuckey, M. S. Abreu Paiva, J. Habib, I. Macaulay, A. J. de Smith, F. al-Beidh, R. Sampson, R. T. Lumbers, P. Rao, S. E. Harding, A. I. Blakemore, S. E. Jacobsen, M. Barahona and M. D. Schneider (2015). "PDGFRalpha demarcates the cardiogenic clonogenic Sca1+ stem/progenitor cell in adult murine myocardium." Nature Communications **6**: 6930.

Shannon, P., A. Markiel, O. Ozier, N. S. Baliga, J. T. Wang, D. Ramage, N. Amin, B. Schwikowski and T. Ideker (2003). "Cytoscape: a software environment for integrated models of biomolecular interaction networks." Genome Res **13**(11): 2498-2504.

Smyth, G. K. (2005). Iimma: Linear Models for Microarray Data. Bioinformatics and Computational Biology Solutions Using R and Bioconductor. Statistics for Biology and Health. R. Gentleman, V. J. Carey, W. Huber, I. R.A. and S. Dudoit. New York, NY, Springer.

Stacklies, W., H. Redestig, M. Scholz, D. Walther and J. Selbig (2007). "pcaMethods--a bioconductor package providing PCA methods for incomplete data." Bioinformatics **23**(9): 1164-1167.

Szklarczyk, D., J. H. Morris, H. Cook, M. Kuhn, S. Wyder, M. Simonovic, A. Santos, N. T. Doncheva, A. Roth, P. Bork, L. J. Jensen and C. von Mering (2017). "The STRING database in 2017: quality-controlled protein-protein association networks, made broadly accessible." Nucleic Acids Res **45**(D1): D362-D368.

Takakura, N., H. Yoshida, T. Kunisada, S. Nishikawa and S. I. Nishikawa (1996). "Involvement of platelet-derived growth factor receptor-alpha in hair canal formation." J Invest Dermatol **107**(5): 770-777.

Takakura, N., H. Yoshida, Y. Ogura, H. Kataoka, S. Nishikawa and S. Nishikawa (1997). "PDGFR alpha expression during mouse embryogenesis: immunolocalization analyzed by whole-mount immunohistostaining using the monoclonal anti-mouse PDGFR alpha antibody APA5." J Histochem Cytochem **45**(6): 883-893.

Wickam, H. (2016). ggplot2: Elegant Graphics for Data Analysis, Springer.
